# Frugivory and seed dispersal by chelonians: A review and synthesis

**DOI:** 10.1101/379933

**Authors:** Wilfredo Falcón, Don Moll, Dennis Hansen

## Abstract

In recent years, it has become clear that frugivory and seed dispersal (FSD) by turtles and tortoises is much more common than previously thought. Yet, a review and synthesis is lacking. We here review published and unpublished records of chelonian FSD, and assess the role of chelonians as seed dispersers, from individual species to the community level. We first discuss the distribution of chelonian FSD and the characteristics of the fruit and/or seed species eaten and dispersed by chelonians. We then use the seed dispersal efficiency framework to explore the quantitative and qualitative components of seed dispersal by tortoises and turtles, embarking on a journey from when the fruits and/or seeds are consumed, to when and where they are deposited, and assess how efficient chelonians are as seed dispersers. We finally discuss chelonian FSD in the context of communities and chelonians as megafauna. We found that a substantial proportion of the world’s aquatic and terrestrial turtles and a major part of testudinid tortoises (70 species in 12 families) include fruits and/or seeds in their diet, and that furits of at least 588 plant species in 120 families are ingested and/or dispersed by chelonians. For some chelonians, overall or in certain seasons, fruit may even form the largest part of their diet. Contrary to seed dispersal by lizards, the other major reptilian frugivores, chelonian FSD is not an island phenomenon in terms of geographic distribution. Nevertheless, on islands especially tortoises are often among the largest native terrestrial vertebrates—or were, until humans got there. We synthesize our knowledge of chelonian FSD, and discuss the relevance of our findings for conservation and restoration, especially in relation to rewilding with large and giant tortoises.

**Resumen:** En años recientes, se ha hecho claro que la frugivoría y dispersión de semillas (FDS) llevada a cabo por tortugas (quelónidos) es más común de lo antes pensado. No obstante, todavía carecíamos de una revisión y síntesis sobre este tema. En este artículo, revisamos récords (publicados y no publicados) sobre FDS por quelónidos, y evaluamos su rol como dispersores de semillas, desde el nivel de individuos, al nivel de comunidades. Primero, discutimos la distribución de FDS por quelónidos, y las características de las especies de frutos y/o semillas consumidas y dispersadas por tortugas. Luego hacemos uso del concepto de la eficiencia de dispersión de semillas como marco de referencia para explorar los componentes cualitativos y cuantitativos de la FDS por quelónidos, embarcándonos en un viaje desde cuando los frutos y/o semillas son consumidas, hasta cuando son depositadas. También evaluamos cuán eficientes son los quelónidos como dispersores de semillas. Finalmente procedemos a discutir la FDS por quelónidos en el contexto de comunidades, y como ‘megafauna’. Encontramos que una proporción substancial de las tortugas acuáticas del mundo y la mayor parte de las tortugas testudínidas (70 especies en 12 familias) incluyen frutos y/o semillas en su dieta que abarcan al menos 588 especies de plantas en 120 familias. En algunas especies, en general o en algunas estaciones, la mayor parte de su dieta está conformada por frutas y/o semillas. Más importante aún, y contrario a las lagartijas, que son otro grupo importante de reptiles que incurre en FDS, la frugivoría y dispersión de semillas por quelónidos no es un fenómeno de islas solamente, en términos de distribución geográfica. Empero, en islas, especialmente las tortugas terrestres, están entre los vertebrados nativos de mayor tamaño–o lo estuvieron, hasta que los humanos llegaron a ellas. En este artículo, hacemos una síntesis de las lecciones aprendidas hasta ahora sobre la FDS por quelónidos, y discutimos la relevancia de nuestros hallazgos para la conservación y restauración, especialmente en relación a proyectos de resilvestrar (‘rewilding’) con tortugas gigantes o de gran tamaño.

## I. Introduction

Animal-mediated seed dispersal is the process by which animals disperse the seeds away from the mother plant (Fig. 1), and is an important ecological function that has profound ecological and evolutionary implications in ecosystems (Howe & Smallwood, 1982; Rezende *et al.*, 2007; Stoner & Henry, 2008). The distribution and ecology of frugivory and seed dispersal (FSD) in most major vertebrate taxa has been thoroughly investigated and results synthesised (Estrada & Fleming, 1986; Levey, Silva, & Galetti, 2002), most recently for lizards (Iverson, 1985; Olesen & Valido, 2003; Valido & Olesen, 2007; Whitaker, 2011), and a start has even been made for crocodilians (Platt *et al.*, 2013). However, a thorough overview and synthesis is still missing for chelonians.

**Figure 1:**
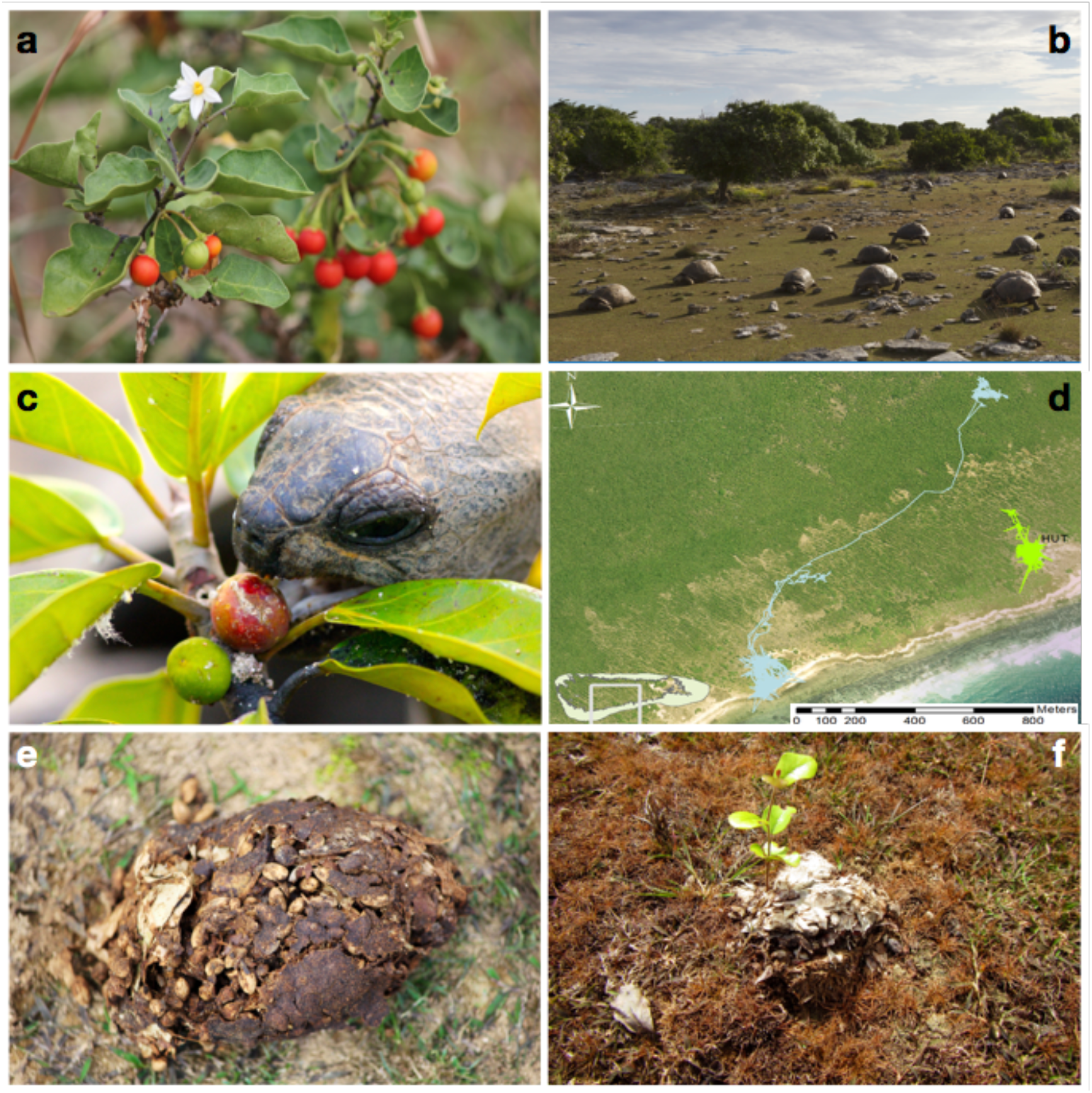
The process and outcome of chelonian-mediated seed dispersal, here exemplified by Aldabra giant tortoises (*Aldabrachelys gigantea*) on Aldabra Atoll, Seychelles. Fruiting plants like the Aldabra tomato (*Solanum aldabrense*) attract giant tortoises (a), which occur at high densities on the atoll (b). Fruits are a large component of the diet of giant tortoises, and they have often been observed eating ripe fruits, while ignoring green ones (e.g., of Ficus nautarum; c). After ingestion, seeds are retained for an average of 15 days in the guts of the tortoises; a time period during which tortoises can move considerable distances across the landscape (d; movement paths of two individuals on the south of the atoll). Once defecated, a single scat of giant tortoises can contain over 150 seeds, and often results in germination (e-f; seeds and a seedling of *Terminalia bovinii*).

Reviewing the origin and rise of frugivory and seed dispersal through deep time, Tiffney (2004) established that plants had the necessary morphological features for vertebrate dispersal by the Late Carboniferous (323.2–298.9 Ma), and that by the middle of the Mesozoic (252–66 Ma), several reptile lineages could have established specific FSD associations with plants. Given the long evolutionary history of chelonians, and the generally broad diet of many extant chelonians could have been among the early dispersers and ‘first movers’ in the evolutionary ecology of fruits (Ridley, 1930; van der Pijl, 1969; Tiffney, 1986; 2004). Perhaps the earliest example of frugivory by chelonians comes from a Campanian (83.6–72.1 Ma) coprolite that likely originated from a turtle, and that contained ca. 200 achenes of a Ranunculaceae sp. (Rodriguez-de la Rosa, Cevallos-Ferriz, & Silva-Pineda, 1998). Two fossilised specimens of *Stylemys* tortoises from the Oligocene (33.9–23.03 Ma) in South Dakota contained hackberry (*Celtis*) seeds (Marron & Moore, 2013). On the Bahamas, two out of three extremely well-preserved individual carapaces of the recently extinct (4,200–1,200 BP) giant tortoise (*Chelonoidis alhuryorum*) contained many seeds of two large-fruited species (wild mastic, *Mastichodendron foetidissimum*, and satinleaf *Chrysophyllum oliviforme;* both Sapotaceae) (Steadman *et al.*, 2007; Franz & Franz, 2009). In historical times, now-extinct giant tortoises (*Cylindraspis* spp.) of the Mascarene Islands were observed by early settlers to include fruit in their diet. In Mauritius in the late 1600s the tortoises were reported to eat ‘apples’ (= endemic ebony *Diospyros*, Ebenaceae, and Sapotaceae fruits) (Hume & Winters, 2016). On nearby Rodrigues Island, the exiled French Huguenot François Leguat and his men ate many fruits from the forest, but “left the dates [= palm fruits, Arecaceae] for the turtles [= giant tortoises, *Cylindraspis* spp.]” (Leguat, 1708).

One of the first modern, experimental FSD studies was Rick & Bowman’s (1961) classic paper on how the germination rate of an endemic Galápagos tomato was dramatically improved by passing through the gut of the endemic giant tortoises. Additionally, Hnatiuk’s (1978) study of germinating seeds from the feces of Aldabra giant tortoises, and Iverson’s (1987) discussionof the likely frugivore mutualistsof the highly specialized Tambalacoque tree (*Sideroxylon grandis*; Sapotaceae), are two early examples of seminal thinking about the potential for seed dispersal interactions and germination enhancement of tortoises and island plants. It is thus ironic that, despite several calls for studies of turtles as seed dispersers (e.g., Moll & Jansen, 1995; Pérez-Emán & Paolillo, 1997), our understanding of chelonian FSD has progressed very little since then. In this review, we aim to summarise published and unpublished information about chelonian FSD in the wild, and synthesise and discuss the role of chelonians as frugivores and seed dispersers. We first present an overview of the taxonomical distribution of chelonian FSD, as well as of the taxonomical distribution of plants consumed by chelonians. We then use the concept of seed dispersal effectiveness (SDE) (Schupp, 1993; Schupp, Jordano, & Gómez, 2010) to discuss the quantitative and qualitative aspects of chelonian seed dispersal. We progress to discuss the function of chelonians as megafaunal seed dispersers in the FSD community, and their role in conservation and restoration efforts.

## II. Methods and data

To synthesise data on FSD by chelonians, we performed a comprehensive literature search that included scientific articles, books, monographs, and theses. We used Google Scholar (http://scholar.google.com/; Google Inc.), as it has been found to include and exceed the results of other commonly used literature databases (specifically WoS and Scopus; see Svenning *et al.*, 2016). We used the following search terms: ‘diet’, ‘frugivory’, ‘seed dispersal’, in combination with the Latin genera of chelonians (from van Dijk *et al.*, 2014), or the keywords ‘chelonian’, ‘tortoise’, or ‘turtle’. No constraints on the year of publication or language were imposed (i.e., we found some articles in other languages, e.g., Spanish). We filtered the search results by reading the abstracts, and also went through the references of each text found to identify other potentially suitable articles. We added literature known by the authors to include diet information, but which did not appear in our search (mostly books). In addition, we added unpublished data based on our own observations and those shared by various researchers. For diet data, we only included information based on wild chelonians. For germination and gut passage experiments we included studies using captive chelonians conducted with fruits found in the natural habitat of the species. To give a more complete overview of some of the main variables that determine the outcome of seed dispersal, we reviewed information on gut retention time (GRT), and on movement ecology and habitat range of chelonians. We used the same approach as above, using each of the search terms, ‘gut retention time’, ‘movement’, ‘activity’ and ‘home range’ together with ‘tortoise’, ‘turtle’ or ‘chelonian’. See Supplementary Materials S1 for the resulting reference lists.

We followed van Dijk *et al.* (2014) for chelonian taxonomy, and the iPlant Collaborative for plant taxonomy (Boyle *et al.*, 2013; http://tnrs.iplantcollaborative.org/). We used the amniote life-history database for data on chelonian body mass (Myhrvold *et al.*, 2015). When studies only showed results graphically, we extracted the data from figures using WebPlotDigitizer ver. 4.0 (Rohatgi, 2017; https://automeris.io/WebPlotDigitizer/). We analysed and visualised the data using R v. 3.3.3 (R Core Team, 2017) and the package ‘ggplot2’ (Wickham, 2016).

We were able to extract data from a total of 167 studies on chelonian FSD, germination, GRT, and movement. We found a total of 106 studies containing data on FSD by wild chelonians. These arose from either focused FSD studies (i.e., studies focusing directly on the role of chelonians as frugivores and/or seed dispersers; n = 24), partial FSD studies (i.e., studies that examine diet in relation to/in a framing of FSD or examine gut passage, but not germination; n = 70), or diet studies (not framed in an FSD context; n = 12). The studies used several methods to obtain data on FSD by chelonians, including direct observation, camera traps (e.g., Wang *et al.*,2011), stomach flushing (e.g., Legler, 1977), or analysis of collected faeces (e.g., Nogales *et al.*, 2017). Faecal collection methods ranged from simple picking up, to more creative approaches, such as collection with a miniature wheeled barrow mounted behind the animal (Josseaume, 2002), and, for marine turtles, collecting in cloaca-mounted bags (Amorocho & Reina, 2008). Determination of the seed content in the faeces was done with either direct counts of seeds, or counting any seeds that germinated from the dung (e.g., Hnatiuk, 1978). For chelonian GRT, we found 37 studies, which were conducted by feeding fruits and/or artificial particles. Finally, we found 24 studies on chelonian movement and home ranges.

There are inherent biases associated with the different methodologies when estimating chelonian FSD. In dietary studies, seeds might often be overlooked, or underreported/not specifically mentioned as plant diet components. For example, Mouden *et al.* (2006) have a long list of plants recorded in scat of the spur-thighed tortoise (*Testudo graeca*), many, but not all of which, overlap with those of Cobo & Andreu (1988), who specifically studied seeds dispersed by *T. graeca*. Also, Kabigumila (2001) and Hansen, Johnson, & Van Devender (1976) provide a long list of food plants found in the scat of the leopard tortoise (*Stigmochelys pardalis*) and the Mojave desert tortoise (*Gopherus agassizii*), respectively, but they did not specify whether these were fruits, seeds or other plant parts. Faecal analysis alone may provide a biased account of a species’ diet. For example, de Lima Magnusson, & da Costa (1997) describe the red side-necked turtle (*Rhinemys rufipes*) as a major frugivore “palm specialist” based on faecal analysis, but a subsequent study by Caputo & Vogt (2008), using stomach flushes, found relatively larger amounts of animal food items. Thus, faecal analysis tends to record more plant matter, while it can grossly underrepresent the importance of animal matter in the diet (Caputo & Vogt, 2008). However, using only stomach flushing may underestimate frugivory, as large seeds are hard to dislodge (Kennett & Tory, 1996; de Lima *et al.*, 1997). A combination of both approaches, where possible, would seem to be ideal. Another aspect which can bias the available data is the seasonality in the diet of some chelonians, where fruit may only be a major part of the diet in some season(s). Short-term studies that do not span different seasons may underestimate fruit consumption and thus the potential for seed dispersal. This is important to into consideration, because only 9% of the studies we found considered seasonality in the diet of chelonians. All these factors underscore the need for a more comprehensive dietary sampling when considering the feeding type of chelonians and their role as frugivores and seed dispersers.

## III. Distribution of chelonian FSD

### (1) Taxonomical distribution

Chelonians comprise about 335 species, of which 275 are turtles and 60 are tortoises, spanning 94 genera in 14 families (van Dijk *et al.*, 2014). We documented FSD in a total of 72 species with, distributed across all major chelonian phylogenetic groups (Table 2; except for Dermochelyidae, with the marine leatherback turtle, *Dermochelys coriacea*, as the only extant species).

There was a notable gap in FSD in the branches containing *Platemys platychephala* to *Acanthochelys* spp. (Chelidae), *Pelochelys* spp. and *Chitra* spp. (Trionichydae), and containing from *Orlitia borneensis* to *Pangshura smithii*(Geomydidae). However, FSD was recorded in other species within these three families. This pattern is likely due to the lack of focused dietary or FSD studies on these species. Moreover, as we will see below, habitat and seasonal influences on the diet of these groups may influence the levels of FSD in different locations and times of the year, therefore affecting sampling results. The few other chelonian species without any reported FSD have been described as purely carnivorous. Thus, frugivory is widespread in Testudines, with most taxa having at least one frugivorous representative at the genus level.

### (2) Geographical distribution

Chelonians are widely distributed across the world, inhabiting habitats from tropical to temperate, from continents to islands and oceans, and they include terrestrial, aquatic and semi-aquatic, as well as marine species (see van Dijk *et al.*, 2014 for individual species distributions). Chelonian species richness peaks in the southeastern USA, the Ganges Delta, Southeast Asia, and northern South America (Fig. 2a; data from Roll *et al.*, 2017, provided by Y Itescu). Thus, unlike FSD by lizards (Olesen & Valido, 2003), FSD by chelonians is not restricted to islands, and they can thus potentially play a major role in continental and island ecosystems alike. The geographic distribution of species richness of chelonian species that engage in FSD is concentrated in the south-eastern USA and northern South America, highlighting the underrepresentation of studies for especially south-east Asia (Fig. 2b).

**Figure 2:**
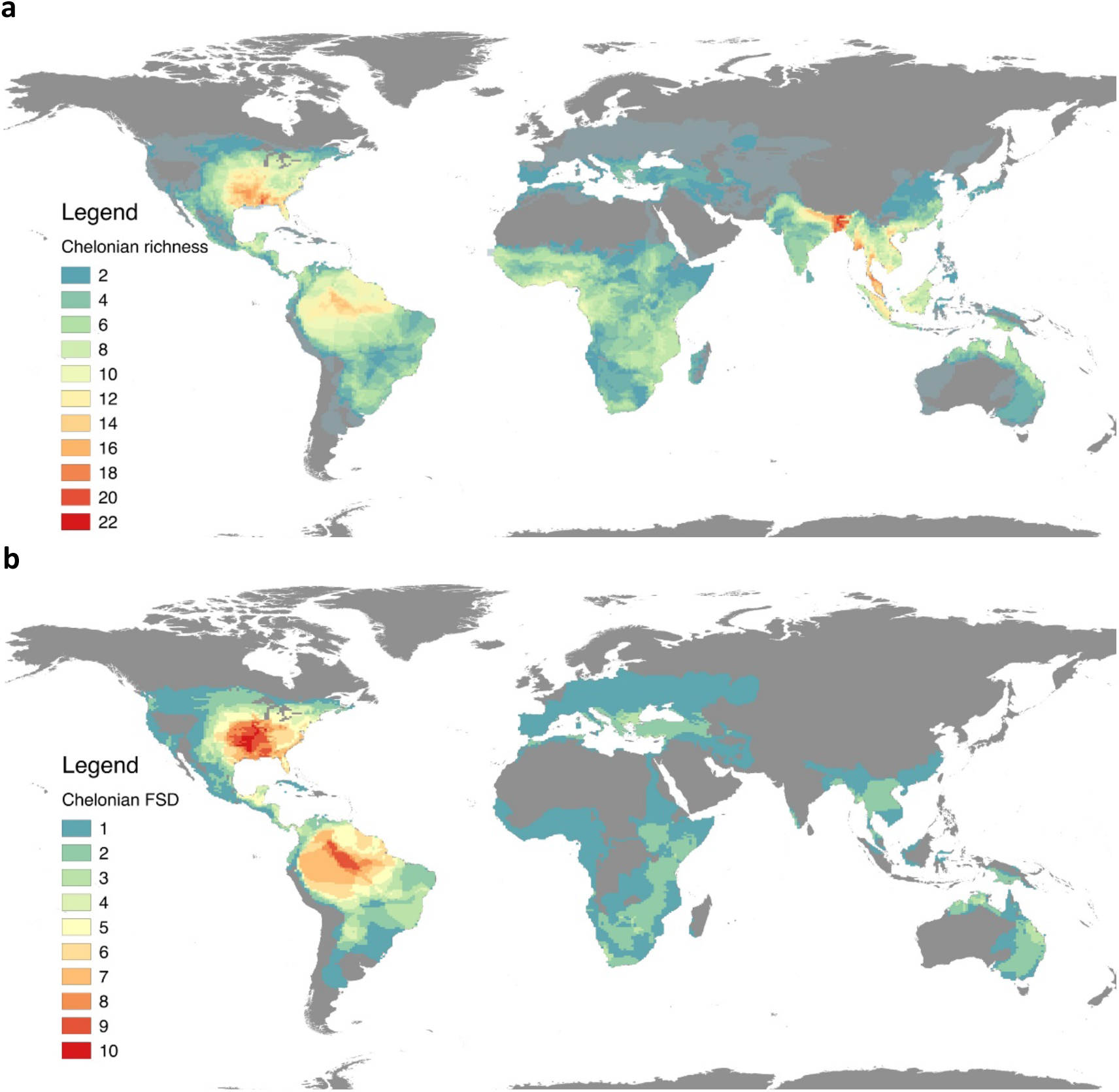
Overall global chelonian species richness (a), and the geographic distribution of chelonians for which we found records of frugivory and/or seed dispersal (b), excluding marine species. Note the difference in magnitude in the colour gradients of the legend.

### (3) Patterns of generalisation and specialisation

Specialisation or generalisation on fruits varies depending on the chelonian species, as expected by the different main feeding types (herbivores, carnivores or omnivores). Frugivorous tortoises can vary from generalist, specialist to opportunistic frugivores. For example, *Chelonoidis* tortoises in South America are generalist frugivores, consuming fruits having a variety of traits (Moskovits, 1985; Guzman & Stevenson, 2008). At the opposite end of the spectrum, we have the highly specialised Gibba turtle (*Mesoclemys gibba [Phrynops gibbus]*) that has been found to feed almost exclusively on palm fruits (*Mauritia flexuosa*, Aracaceae) during part of the year in the Rio Negro Basin in Brazil (Caputo and Vogt, 2008). Other species such as the common snapping turtle (*Chelydra serpentina*) are omnivorous, incorporating roughly the same amount of plant material (including fruits) and animal material in their diet (Ernst & Lovich, 2009). Lastly, there are species that are mostly carnivorous, which will eat fruits opportunistically, such as Blanding’s turtle (*Emydoidea blandingii*) (Rowe, 1992) and hinge-back tortoises (*Kinixys* spp.) (Luiselli, 2003). Overall, most frugivorous chelonian species are generalist frugivores that also include other plant material in their diet; this is especially true for tortoises (Testudinidae).

### (4) Functional traits in relation to FSD

Frugivore species have inter- and intraspecific differences in functional traits, such as habits, size and age, sex, cognition and preferences, which may result in large differences in the seed dispersal services they provide (Jordano *et al.*, 2007; Zwolak, 2017). Knowledge about these traits will help us understand the role of specific characteristics of frugivores in their effectiveness as seed dispersers.

#### (a) Habitat

Chelonians are a diverse group of vertebrates whose different habits, such as terrestrial, semi-aquatic, aquatic and marine, have allowed them to exploit many habitats and resources. Terrestrial plants and tortoises are perhaps the first that come to mind when thinking about seed dispersal in this group. With their preponderance of fleshy fruits, terrestrial plants could be considered more zoochorous than their aquatic counterparts, and because of their habitat, tortoises and terrestrial –or semi-aquatic–turtles are more likely to encounter fruits and disperse their seeds within terrestrial habitats. However, as we found, seed dispersal is also carried out by mainly aquatic species, both on land and in water, for terrestrial and aquatic plants, and even for coastal and marine plants in marine ecosystems. Yet, most studies of chelonian FSD have focused on terrestrial chelonians andlargely ignored the role of aquatic and marine species in seed dispersal (Moll and Jansen 1995). Ultimately, the habitats of both chelonians and of the plants they encounter will determine which fruits are available to each species, and whereto the seeds can be dispersed.

#### (b) Size and age

Tortoises and turtles exhibit great inter- and intraspecific size variation. Size generally increases with age in chelonians (Carr, 1952; Ernst and Lovich, 2009). From the perspective of FSD, the size of chelonians limits the size and the number of fruits and/or seeds they can swallow and pass through their guts. Furthermore, size may affect gut passage time (see section on mouth and gut passage treatment) and volume of the scat. Thus, size is expected to substantially affect the ability and effectiveness of chelonians as seed dispersers (see also section on chelonians as megafaunal seed dispersers).

Ontogenetic changes in diet may also occur in chelonians, with vegetation becoming more important as chelonians age and become larger (Moll, 1976); this seems to be common in omnivorous turtles (Clark & Gibbons, 1969; Georges, 1982; Hart, 1983; Sung, Hau, & Karraker, 2016). In the case of the omnivorous red sidenecked turtle (*Rhinemys (Phrynops) rufipes*), de Lima *et al.* (1997) found that most of the scat volume was palm seeds, and that the frequency of palm seeds increased with turtle size. These ontogenetic changes in diet may be accompanied by changes in gut morphology, as found in the green sea turtle (*Chelonia mydas*), with the ratio of long to short intestines increasing from 0.45 in post-hatchlings to 2.5 in adults, possibly reflecting a higher proportion of animal matter in the diets of young individuals (Davenport, Antipas, & Blake, 1989).

#### (c) Sex

Sexual dimorphism is common in chelonians, but the direction of sexual dimorphism depends on the species and even on habitat. For example, males of angulate tortoises (*Chersina angulata*) are larger than females, whereas females of leopard tortoises (*Stigmochelys pardalis*) are larger than males within the same habitat (Mason *et al.*, 2000). In the case of Aldabra giant tortoises (*Aldabrachelys gigantea*), the population exhibits no sexual dimorphism on the east of Aldabra Atoll, but males gradually attain larger sizes compared to females towards the western side of the atoll (Turnbull *et al.*, 2015). As described above, size is expected to have a differential effect on seed dispersal, and there may thus be differences in the seed dispersal provided by males and females, respectively. For example, where sexual dimorphism is present, the larger males of Aldabra giant tortoises are able to extend their necks to reach higher vegetation and fruits than the smaller females can (WF & DMH, pers. obs). Males and females may also exhibit different behaviours, e.g., habitat selection, which can affect the outcome of FSD. For example, most of the stomach contents of the omnivorous female smooth softshell turtle (*Apalone mutica*) were aquatic items, whereas stomach content of males was mostly terrestrial items and included more fruits (Plummer & Farrar, 1981). These sexual differences in terms of diet were attributed to the different microhabitat preferences (females forage in deep water, whereas males forage in the interface between aquatic and terrestrial habitats). Furthermore, males and females of some species may show differences in home range size and displacement distances (see below). Difference in habitat selection, home range size and displacement distances are not only expected to affect the ability of chelonians to exploit fruits, but also their effectiveness as seed dispersers.

#### (d) Cognition and behaviour

Chelonians, as other animals, rely on cognitive processes to acquire knowledge about their environment through their senses, leading to learning and memory creation. The sensory features of fruits play an important role in attracting frugivorous birds and aid in their selection (Schaefer, Spitzer, & Bairlein, 2008b), and this is expected to be the case for chelonians as well. Sight and olfaction are the sensory faculties that aid turtles and tortoises in the recognition of food sources. Chelonians can perceive images and distinguish colours in the human-visible spectrum (Granda & Stirling, 1965; Baylor & Fettitplace, 1975; Schwartz, 1975; Neumeyer & Jäger, 1985; Arnold & Neumeyer, 1987; Ammermüller, Muller, & Kolb, 1995; Ventura *et al.*, 2001; Twig & Perlman, 2004; Mathger, Litherland, & Fritsches, 2007; Pellitteri-Rosa *et al.*, 2010), and some have been shown to also have sensitivity to the ultraviolet spectrum (Ammermüller *et al.*, 1998; Ventura *et al.*, 1999; Zana *et al.*, 2001). Additionally, chelonians have a highly developed olfactory (vomeronasal) system (Manteifel, Goncharova, & Boyko, 1992; Murphy, Tucker, & Fadool, 2001; Fadool, Wachiowiak, & Brann, 2001), which they can use to detect volatile chemicals excreted by plants from long distances (King, 1996), and also to smell fruits at close range, possibly to evaluate ripeness (WF, DMH, DM, pers. obs.).

Learning and memory of frugivores has an important impact on seed dispersal, because decision-making based on previous experiences can determine which plants and which fruits are selected and consumed, and ultimately where seeds are dispersed (reviewed in John *et al.*, 2016). A model by John *et al.* (2016) testing frugivores with different spatial memory skills suggested that those with longer spatial memory are able to relocate food sources more efficiently, survive longer and disperse larger amounts of seeds. The also moved less at random around the landscape, which led to differences in terms of the spatial distribution of seeds dispersed compared to animals with shorter memory. Captive red-footed tortoises (*Chelonoidis carbonarius*) can navigate efficiently in their environment, and they can remember spatial location of food for at least two months (Soldati, 2015). Moreover, they were able to anticipate food availability over periods of 24 h, discriminating between the quality and quantity of food, and remembering these attributes for at least 18 months. In relation to large-scale movement patterns, individuals of both Galápagos (Blake *et al.*, 2013) and Aldabra giant tortoises (Baxter 2015) have been shown to follow the same movement patterns in different years, implying that they have a persistent spatial memory.

Chelonians may use landmarks and different stimuli to orient themselves and find suitable food sources. For example, when tested in a T-maze, sulcata tortoises (*Geochelone sulcata*) and leopard tortoises (*Stigmochelys pardalis*) could discriminate between colours and shapes, and associate these features with navigation to food sources (Janisch, 2013). Red-footed tortoises (*Chelonoidis carbonarius*) can navigate between known localities where fallen fruits are available at certain seasons (Josseaume, 2002). Also, fallen fruits encountered are often from foraging activity of arboreal/aerial frugivores (Moll & Jansen, 1995), and it is thus possible that chelonians can use cues from other species to find food. This seems to be the case in in Malaysia, where painted terrapins (*Batagur borneoensis*) have been observed clustering in the water under a troop of leaf monkeys in trees above to eat berembang fruits (*Sonneratia caseolaris*, Lythraceae) that the monkeys were throwing into the water (Moll, 1980b).

By navigating the landscape based on previous experiences, chelonians can identify and exploit fruits. For example, (Legler, 1976) noted that the gulf snapping turtle (*Elseya lavarackorum*) in Australia exploits windfall fruits of fig trees, with large congregations of these turtles found around this resource. In addition, other aquatic species such as the black river turtle (*Rhinoclemmys funerea*) (Moll & Jansen, 1995) and the Central American river turtle (*Dermatemys mawii*) (Moll, 1989) have been observed clustering and waiting in water beneath fruiting *Ficus glabrata*(Moraceae) trees, and the painted terrapin (*Batagur (Callagur) borneoensis*) displays similar clustering in the water under falling berembang fruits in Malaysian rivers (Moll, 1980b). Similarly, the Travancore tortoise in India (*Indotestudo travancoria*)(Bonin, Devaux, & Dupré, 2006; Kanagavel & Raghavan, 2012), and in Brazil, the red-footed tortoise (*Chelonoidis carbonarius*) (Moskovits & Bjorndal, 1990) congregate beneath favoured fruiting trees to exploit these food resources. Notably, the tree *Spondias testudinis* (Anacardiaceae) was named for the yellow-footed tortoises (*Chelonoidis denticulatus*) (Mitchell & Daly, 1998) that flock beneath fruiting trees to feed on the large, yellow-brown fruits (D Daly, pers. comm.). Furthermore, aquatic Antillean sliders (*Trachemys decussata*) in Cuba will emerge onto the land in great numbers after rains to feed on fallen jobo (*Spondias lutea*) and Bagá(*Anona palustris*, Annonaceae) fruits that have fallen from riparian trees (Barbour & Carr, 1940). Thus, chelonians possess a landscape-scale spatial awareness of plants providing fruits.

#### (e) Fruit preferences

Animals rely on their ability to detect differences in food quality by using sensory adaptations, which allows them to circumvent some of the costs associated with foraging (Borges *et al.*, 2011). Frugivores can establish and maintain preferences based on colour, odour and taste (Sorensen, 1983; Levey, 1987; Willson, Graff, & Whelan, 1990; Willson & Comet, 1993). As discussed above, chelonians have highly developed visual and olfactory systems, and are known to be attracted by smell and colour (see Harless & Morlock, 1979 for a review), which may lead to the establishment of preferences. Indeed, studies focusing on colour preferences in tortoises have found preferences for distinct visual stimuli. Probably the first study that explored colour preferences in chelonians was done by Grant (1960) on Texas tortoises (*Gopherus berlandieri*), which exhibited a strong preference for red, selecting food items dyed red after having initially rejected them (i.e., when the same food items had other colours). Subsequent studies using spur-thighed tortoises (*Testudo graeca*) (Pellitteri-Rosa *et al.*, 2010), yellow-footed tortoises (*Chelonoidis denticulatus*) (Passos, Santo Mello, & Young, 2014), and Aldabra giant tortoises (*Aldabrachelys gigantea*) (Spiezio, Leonardi, & Regaiolli, 2017; DMH, unpublished) have shown a prevalent preference for yellow, and/or red colours. Furthermore, chelonians have been shown to discriminate between odours to identify potential mates and conspecifics (e.g., Auffenberg, 1965; Galeotti *et al.*, 2009; Polo-Cavia, López, & Martín, 2009), and they also use scent to find food items (Germano *et al.*, 2014). Although chelonians are also known to discriminate shapes (Janisch, 2013), we did not find any studies examining food or fruit shapes as visual stimuli, nor did we find any studies on taste discrimination.

Plants are known to employ visual and scent cues to signal ripeness in fruits to attract seed dispersers, which use these cues to assess their nutritional value (Brady, 1987; Kalko, Herre, & Handley, 1996; Schlumpberger, Clery, & Barthlott, 2006; Schaefer, McGraw, & Catoni, 2008a). Unripe fruits often have chemical compounds that make them unpalatable to seed dispersers (Sherburne, 1972; Schaefer, Schmidt, & Winkler, 2003), who may learn to associate visual and scent cues with unpalatability. Therefore, different colour and smell preferences may ultimately lead to distinct preferences for certain fruit traits. For example, many fruits are green when unripe, and yellow or red when ripe, and the ripening process is usually accompanied by the release of scents. Consequently, we can expect chelonians to have different preferences for different fruit species, be able to discern between ripe and unripe fruits, and show a preference for ripe ones, especially those that become yellow and red.

The degree to which chelonians act as valid seed dispersers rather than only as frugivores depends on the selection of fruits with viable seeds (usually ripe). Moskovits & Bjorndal (1990) showed that the red- (*C. carbonarius*) and yellow-footed tortoises (*C. denticulatus*) preferred fruits over other food items, and preferred fruits that were predominantly red or yellow and were fragrant while rejecting unripe fruits. Chelonians have been observed smelling ripe and unripe fruits at close proximity before eating or apparently rejecting them. For example, this behaviour has often been observed in Aldabra giant tortoises (WF and DMH, pers. obs.; Fig. 1c). Similarly, the eastern box turtle (*Terrapene carolina carolina*)seems to be able to distinguish between ripe and unripe fruits, preferring the ripe ones (Allard, 1948). However, it should be noted that Hermann’s tortoises (*Testudo hermanni*) consumes unripe green fruits of *Ruscus aculeatus* (Asparagaceae) when seasonally available (Del Vecchio *et al.*, 2011), probably limiting their effectiveness as seed dispersers.

The only experimental study that we are aware of that simultaneously evaluated the perception of colour, olfaction and taste was by Grant (1960), studying the Texas tortoise. He proposed, based on feeding trials, that vision, olfaction, and taste, in that order, were used to by the tortoises to select food items. Thus, rather than just relying on one or the other, chelonians use sight and olfaction and taste to discriminate between possible food sources (Grant, 1960; Fitch, 1965; Pellitteri-Rosa *et al.*, 2010), and when fruits and seeds become available in their habitat, they are probably effective at finding them (Moll & Jansen, 1995).

It is worth mentioning that, although it is often assumed that tortoises benefit from seed dispersal interactions with plants by obtaining food resources, it appears that some chelonians do not derive nutritive benefits from these interactions. For example, alligator snapping turtles (*Macrochelys temminckii*) eat many acorns (such as those of the willow oak, *Quercos phellos*), and significantly enhance their germination (Elbers *et al.* 2011). However, the acorns pass through the gut seemingly unscathed and unchanged, so why are they ingested by the turtles in large numbers? Are they covered by nutritious microorganisms after they have soaked in water and that benefit the turtles, or perhaps act as roughage to help grind up other ingested foods? More research is needed to determine the mechanisms by which chelonians are attracted to seemingly unnutritious fruits and/or seeds, and whether they truly present nutritional or other benefits to these reptiles.

## IV Plants eaten and dispersed by chelonians

### (1) Taxonomical distribution

Chelonians consume the fruits and/or seeds of a great number of plants, including at least 588 species belonging to 368 genera in 121 families. These plant species are distributed across the phylogenetic tree of angiosperms. These plant species occur in many different habitats, with a variety of growth habits, and possess fruits and seeds with a myriad of traits (see Supplementary Materials S2 for the list of plant fruit and/or seed species consumed and/or dispersed by chelonians). Only 18% of all plant families, however, had more than 10 species whose fruits and/or seeds are consumed and/or dispersed by chelonians (Table 1), with 27% of families represented by only a single plant species.

**Table 1:**
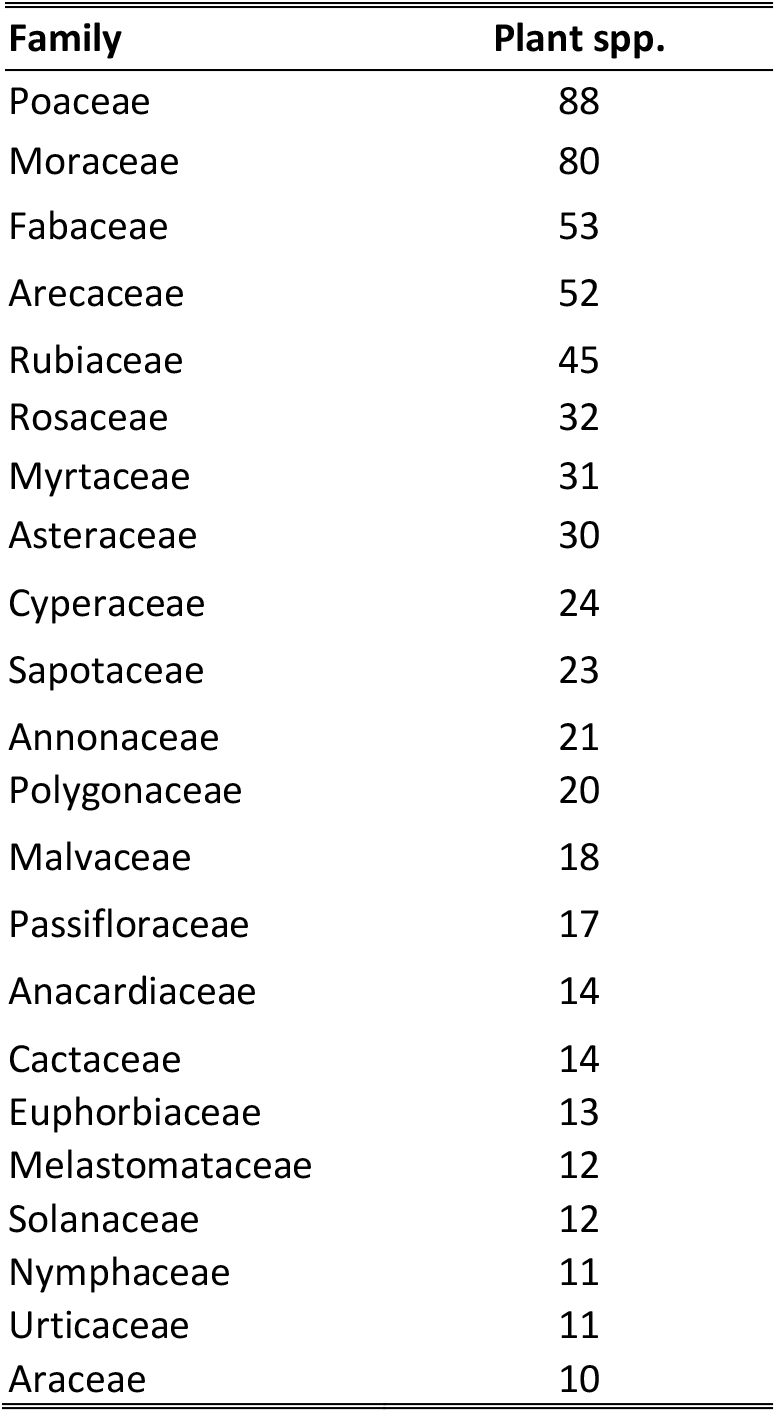
Plant families of fruits and/or seeds most commonly eaten by chelonians.

### (2) Modes of dispersal

There are two modes of chelonian seed dispersal: endozoochory (dispersal of seeds through the ingestion of fruits and/or seeds), and epizoochory (dispersal of seeds on external body parts). Of these, endozoochory is by far the most common mode, forming the majority of cases reviewed in our study. It occurs in terrestrial, aquatic, and even in marine ecosystems. During the process of endozoochory, the handling behaviour, gut treatment and location of defecation all affect the ultimate quality of seed dispersal (see below). Epizoochory is a passive way of dispersal where seeds are stuck on the external parts of the animals until they are subsequently dropped, and other than movement away from the mother plant, the fruits or seeds are not affected further. Epizoochory has only been observed in two species of chelonians. The terrestrial Aldabra giant tortoises (*Aldabrachelys gigantea*), which disperse the sticky seeds of *Plumbago aphylla* (Plumbaginaceae) that adhere to their carapaces, and secondarily disperse seeds of various plant species on their carapaces after birds defecate on them (e.g., *Ficus* spp., Moraceae; WF and DH, pers. obs.). In Australia, the aquatic eastern long-necked turtle (*Chelodina longicollis*), disperses several wetland-associated plants whose seeds lodge on its carapacial algal mats (Burgin & Renshaw, 2008).

### (3) Diversity of seeds

The species diversity of seeds potentially dispersed by chelonians varies by chelonian species and/or studies. Overall, frugivorous chelonians covered in our review each potentially disperse a high diversity of seeds, with a mean of 13.0 plant species per chelonian species (± 23.6; range: 1–123; see Supplementary Materials S2 for species dispersed), and for some chelonians fruits and seeds were major parts of their diets. For example, the Gopher tortoise (*Gopherus polyphemus*) disperses more than 50 species of seeds in pine savannah in the southestern USA (Birkhead, Guyer, & Hermann, 2005). For the big-headed Amazon river turtle (*Peltocephalus dumerilianus*) fruits and seeds were the most diverse components in the diet, with a total of 19 species found in the stomachs, and with Aracaceae (palm) seeds as the most common ones (Pérez-Emán & Paolillo, 1997). In the northern giant musk turtle (*Staurotypus triporcatus*), the large seeds of *Diospyros digyna* (Ebenaceae) comprised 63% of the volume of their stomach contents (Vogt & Guzman, 1988). It should be noted that although careful studies have documented many dry-seeded species dispersed –and potentially dispersed–by chelonians (e.g., Hnatiuk, 1978; Cobo & Andreu, 1988; Milton, 1992; Birkhead *et al.*, 2005), there is likely an underestimation in the amount and diversity of such seed species when compared to fleshy-fruited species due to the difficulty of detection and/or identification.

### (4) Plants only/mostly dispersed by chelonians

van der Pijl (1969) suggested that fruits dispersed by reptiles (saurochory) should be coloured, smelly, and borne near the ground or drop at maturity. Although strong FSD relationships have been documented between plants and some chelonians, there is a lack of evidence of any form of coevolution that has resulted in a chelonian seed dispersal syndrome (Herrera, 1985). As mentioned above (see section on preferences), although they may show preferences, chelonians potentially disperse fruits with a wide variety of s sizes, colours, and scents. For example, although they show preferences for certain fruits, *Chelonoidis* tortoises consume fruits with a variety of colours, including both fragrant and odourless ones (Moskovits, 1985; Guzmán & Stevenson, 2008).

However, certain plants may rely disproportionally on chelonians for seed dispersal. For example, while rodents and birds destroy the seeds of *Pandanus aquaticus* (Pandanaceae), northern Australian snapping turtles (*Elseya dentata*)defecate the seeds intact (Kennett & Russell-Smith, 1993). Similarly, European pond turtles (*Emys orbicularis*) disperse most of the seeds of *Nymphaea alba*(Nymphaceae) intact, while ducks, coots and fish destroy the seeds after gut passage (Calviño-Cancela, Ayres Fernández, & Cordero Rivera, 2007, and references therein). Wang *et al.* (2011) found that red-footed tortoises (*Chelonoidis carbonarius*) may be an important seed disperser of *Syagrus flexuosa* (Arecaceae), because the seeds were often defecated undamaged but are rarely found at all in the scat of other animals. Furthermore, Moll & Jansen (1995) suggested the black river turtle (*Rhinoclemmys funerea*) as an important seed disperser of *Ficus glabrata* (Moraceae) and *Dieffenbachia longispatha* (Araceae). This turtle is very abundant, practices “windfall” feeding in water under riparian fig trees, and emerges on riverbanks and defecates seeds while on land along shorelines in optimal growing locations for these plants. Tortoises may also be especially important for the dispersal of large-seeded plant species on islands (Heleno *et al.* 2011; Blake *et al.* 2012; Falcón 2018), which has important implications at the ecosystem level (see section on chelonians as megafaunal seed dispersers).

Grasslands (composed of grasses and sedges) are an important food source in the diet of different terrestrial chelonians (e.g., eastern Hermann’s tortoise, *Testudo hermanni boettgeri*) (Rozylowicz & Popescu, 2013). In the case of Aldabra giant tortoises on Aldabra, grasslands are the most preferred habitat (Walton *et al.*,in review), where the high grazing pressure led to the evolution of a specialised ‘tortoise turf’ plant community, whose seeds they disperse (Merton, Bourn, & Hnatiuk, 1976; Hnatiuk, 1978). For the green sea turtle (*Chelonia mydas*) in the Great Barrier Reef (Australia), seagrass is an important dietary component and it disperses its seeds (Tol *et al.*, 2017). The only other known seed disperser in the Great Barrier Reef is the dugong (*Dugong dugon*) (Tol *et al.*, 2017), which is considered vulnerable and occurs in low numbers, so turtles may be more important in terms of quantity. Additionally, the diamondback terrapin is also known to be a seed disperser for the eelgrass (Zosteraceae), a type of seagrass, in the Lower Chesapeake Bay (Tulipani & Lipcius, 2014).

Although chelonians do not necessarily seek for grass seeds per se (but see Kimmons and Moll, 2010, turtles may eat floating grass seeds from water surface), and rather act mainly as herbivores, grasses, sedges and seagrasses in general have traits that facilitate chelonian seed dispersal, and it could be important for the maintenance of such communities (Merton *et al.*, 1976; Hnatiuk, 1978; Tol *et al.*,2017). As Janzen (1984) puts it, “the foliage is the fruit”, and the role of chelonians as seed dispersers in grass communities is likely to be of great importance in places where they reach high densities and levels of biomass, like in island ecosystems or in some places in Africa, especially when compared to other seed dispersers (e.g., Coe, Bourn, & Swingland, 1979; Branch, 2008). It should be noted, however, that the six-tubercled Amazon river turtle (*Podocnemis sextuberculata*) seems to be a predator of Poaceae and Cyperaceae seeds in the Amazonas, which constituted 92% of their stomach volume contents (Fachín-Terán & Vogt, 2014). In all cases, proper viability, germination and recruitment studies are necessary to determine whether effective seed dispersal occurs.

## V Chelonian seed dispersal efficiency

The ultimate definition of efficient animal-mediated seed dispersal is that a dispersal event results in the successful establishment of new reproducing plant individuals. This, however, is far from always the case, as different frugivore species do not provide the same dispersal services to plants. The seed dispersal effectiveness (SDE) framework (Schupp, 1993; Schupp *et al.*, 2010) provides a way to estimate the contributions of individual dispersal agents to the overall dynamic of plant populations Essentially, it quantifies the number of seeds dispersed by a frugivore multiplied by the probability that a dispersed seed produces a new adult plant. As such, the SDE framework has two components: a quantitative and a qualitative one, which, in turn, have many variables, demographic parameters and subcomponents.

The SDE framework can thus be used as a valuable organising tool to study the ecological and evolutionary implications of seed dispersal. Below we discuss chelonian FSD in the context of the SDE framework.

### (1) Quantitative component

The quantitative component of SDE can be reduced to the number of foraging visits a chelonian makes to a fruiting plant multiplied by the number of seeds dispersed for each visit (Schupp *et al.*, 2010). The former can be affected, for example, by the local abundance of both plants and chelonians, and the chelonian’s degree of frugivory, while the latter is influenced by the numbers of fruits and/or seeds handled per visit, handling behaviour, and body size (for body size, see section on chelonian functional traits).

#### (a) Local biomass and density

Perhaps the most comprehensive work to date on chelonian biomass and density is that of Iverson (1982), who argued that despite the important role that reptiles play in terms of the energetics at the ecosystem level, the study of chelonian abundance and biomass was a neglected subject. He calculated biomass of chelonians based on population density estimates, and analysed those data in terms of habit, habitat, and trophic position. He found that typical values of chelonian biomass are at least one order of magnitude higher than those of other ectotherm species. He also found indications that herbivorous chelonians, which often include fruits as part of their diet, appear to have higher biomass than omnivorous or carnivorous species. Finally, he found that annual production estimates in chelonians (with a maximum of 528 kg ha^-1^ yr^-1^) are similar to most other vertebrate groups, except for fishes; and that the maximum biomass for individual tortoise species could be as high as 586 kg ha^-1^. In terms of density, studies have provided estimates for several species; for example, 0.15–0.31 individuals ha^-1^, for the highly frugivorous yellow-footed tortoise in the Amazon (Guzmán & Stevenson, 2008), 0.85 tortoises ha^-1^ for leopard tortoises and 0.12 individuals ha^-1^ for angulate tortoises in South Africa (Mason *et al.*, 2000).

In some species, chelonian biomass may be higher than that of many classes of larger mammals. For example, Branch (2008) indicated that the leopard and angulate tortoise biomass is about 13% that of all mammalian herbivores in South Africa’s Eastern Cape province, where tortoises can reach high densities (Mason *et al.*, 2000). He posited that this meant that the total biomass of tortoises there almost equalled the combined biomass of kudu, buffalo, eland, and bushbuck, only being exceeded by that of elephants! Moreover, Coe *et al.* (1979) estimated the biomass of Aldabra giant tortoises to range between 253.42–353.87 kg ha^-1^ on Aldabra Atoll, which is much higher than that exhibited by large mammalian herbivores on Africa. However, it should be noted that chelonian biomass is limited by different factors, such as habitat type (e.g., in mesic vs. xeric habitats) (McMaster & Downs, 2006), and can differ between co-occurring species (Mason *et al.*, 2000). Nevertheless, in general, we can expect the total numbers of seeds dispersed per hectare per year to be large for chelonians (see section on quantity of seeds dispersed), especially when considering the number of large seeds dispersed (Jerozolimski, Ribeiro, & Martins, 2009).

#### (b) Degree of frugivory

The degree of frugivory in chelonians varies between species, and within species it can vary at the population and at the individual level. For example, in Mexican giant mud turtles (*Stauratypus triporcatus*), fruits and seeds were the most important dietary component across two sites in Los Tuxtlas (Mexico), but the occurrence of frugivory ranged from 38–100% between populations, and fruits and seeds represented values between 55–82% of the stomach content volume examined (Vogt & Guzman, 1988). The degree of frugivory can also vary depending on the size of chelonians. For example, Sung *et al.* (2016) found a positive relationship between the size of big-headed turtles (*Platysternon megacephalum*) and the occurrence of fruits in their diet. Moreover, diet can vary much over short distances. Another aspect to take into consideration is the changes in diet depending on which habitat chelonians inhabit, and depending on seasons. Geoffroy’s sidenecked turtle (*Phrynops (Rhinemys) geoffroanus*) may have different diets depending on whether it inhabits clean or polluted rivers (Medem, 1960, cited in Fachín-Terán, Vogt, & Gomez, 1995); Souza & Abe, 2000), and depending on season (e.g., fruits of Myrtaceae and Sapotaceae were only found in its stomach during the season of rising water levels) (Fachín-Terán *et al.*, 1995). Likewise, the Gibba turtle (R. *gibbus*)has been found to feed almost exclusively on palm fruits (buriti) only during part of the year in the Rio Negro Basin in Brazil (RC Vogt, pers. comm.). Similarly, inclusion of fruits in the diet can shift seasonally in the smooth softshell turtle (*Apalone mutica*) (Plummer & Farrar, 1981) and the Mexican mud turtle (Kinosternon integrum) (Macip-Rios *et al.*, 2010). In addition, changes in diet can occur at the same location (e.g., a river) over time, as the habitat and food resources change over time (e.g., river changes from clean to polluted) (Moll, 1980a).

#### (c) Quantity of seeds dispersed

Propagule pressure influences the establishment of plants, and the number of seeds dispersed can thus determine the dynamics of plant recruitment. Studies on chelonians indicate that tortoises and turtles are capable of dispersing a high number and diversity of seeds. For example, in the red-footed tortoise (*Chelonoidis carbonarius*), Wang *et al.* (2011) reported that a single scat sample contained high numbers of seeds, ranging from 22 to 765 seeds. Moreover, Lagler (1943) found 11,065 seeds of *Nymphaea alba* in the digestive tract of one individual of the common snapping turtle (*Chelydra serpentina*). Combining information on density estimates and information on their diet and seed dispersal ecology, Guzmán & Stevenson (2008) estimated that yellow-footed tortoises disperse 160.70 seeds ha^-1^ per year.

### (2) Qualitative component

The qualitative component of SDE can be reduced to the probability that a dispersed seed survives handling by chelonians in a viable condition (quality of treatment in the mouth and gut) multiplied by the probability that a viable dispersed seed will survive, germinate, and produce a new adult (quality of deposition) (Schupp *et al.*, 2010).

#### (a) Mouth and gut passage treatment

Lacking teeth, most chelonians tend to swallow fruits and seeds whole (“gulpers”), rather than chewing them as other vertebrate groups do (Moll & Jansen, 1995). They use ‘lingual prehension’, which is the behaviour of using the tongue to touch food items to insert them into their mouths, and this is obligatory for tortoises (Wocheslander, Hilgers, & Weisgram, 1999; Bells *et al.*, 2008). Amphibious emydids and geoemydids use their jaws to grasp food items in terrestrial habitats, a behaviour known as ‘jaw prehension’ (Heiss, Plenk, & Weisgram, 2008; Natchev *et al.*, 2009; 2015). Moreover, and different from birds and monkeys, tortoises do not regurgitate/spit seeds. Thus, damage to seeds by the mouthparts of chelonians was minimal in the studies evaluated. For example, most of the large numbers of seeds of *Nymphaea alba* (Nymphaceae) found in the digestive tract of the common snapping turtle were mature, and very few of the coats were ruptured (Lagler, 1943). However, some chelonian species can damage seeds with their mouths before gut passage. For example Caputo & Vogt (2008) reported that seeds of several plant species were never recovered whole from stomach flushing in the red side-necked turtle (*Rhinemys (Phrynops) rufipes*). Similarly, seeds of two species of plants were found crushed inside the stomachs of the giant South American river turtle (*Podocemis expansa*) (Goulding, 1980).

After consuming the fruits or seeds, they pass to the stomach and through the gut before being defecated. The overall effect on seeds can vary, depending on digestion efficiency and gut retention time (GRT; the time seeds take to pass through the guts until being defecated). Food intake rates may differ among food types in herbivorous chelonians, which have a flexible dietary response, with the ability of switching between cell wall fermentation and extraction of cell contents depending on the diet (Bjorndal, 1989). Moreover, digestive efficiency is inversely related to food intake in tortoises (Meienberger, Wallis, & Nagy, 1993). In some instances, digestive efficiency can depend on the degree of herbivory the species considered, and upon the types of fruits consumed (e.g., in the box turtles *Terrapene carolina* and *T. ornata*) (Stone & Moll, 2009), while in others, such as yellow- and red-footed tortoises (*Chelonoidis denticulatus* and *C. carbonarius*, respectively), for a given diet, neither digestibility nor mass-specific intake varied between species, and neither did they vary by sex or body mass within each species (Bjorndal, 1989).

Chelonians seem to submit digesta to a similar degree of ‘gut washing’ as mammalian herbivores do (Franz *et al.*, 2011). However, although herbivorous reptiles have similar digestibilities as mammalian herbivores (Bjorndal, 2012), overall chelonians are said to be inefficient feeders because their performance at digesting cellulose is lower when compared to mammalian herbivores, and they need to eat large quantities of food to satisfy their energy demands (Branch, 2008). As a result, plant items in their scat are often recognisable, and seeds often pass undamaged.

Compared to the other vertebrate groups, chelonians have relatively longer GRTs, with a mean of 7.65 days (± 5.89; for all species examined combined, Fig. 3), due to their low metabolic rates and food intake (Stevens & Hume, 2004; Franz *et al.*, 2011). Gut retention times in chelonians may be affected by a myriad of factors. For example, GRT tends to vary across seasons, especially in habitats where there are wet and dry periods (e.g., *Aldabrachelys gigantea*) (Coe *et al.*, 1979). Temperature also plays a role in regulating GRT, with increasing temperature leading to faster passage (Sadeghayobi *et al.*, 2011). Moreover, GRT depends strongly on fruit species consumed and on overall diet composition (Bjorndal, 1989; Stone & Moll, 2006). For birds, secondary metabolites in fruits are known to affect GRTs (Murray *et al.*, 1994; Wahaj *et al.*, 1998), which is likely the case in chelonians as well. Furthermore, tortoises show variation in their intestinal morphology according to their feeding habits, and the length ratio of large to small intestines is positively related with GRT (Hailey, 1997). Also, chelonians may exhibit selective food retention based on particle size (Hatt *et al.*, 2002), with coarser food being retained for longer (Hailey, 1997). Lastly, chelonians may exhibit antiperistalsis in the large intestine (i.e., contents are carried upwards) (Naitoh, Hukuhara, & Kameyama, 1975), which also likely affects GRT.

**Figure 3:**
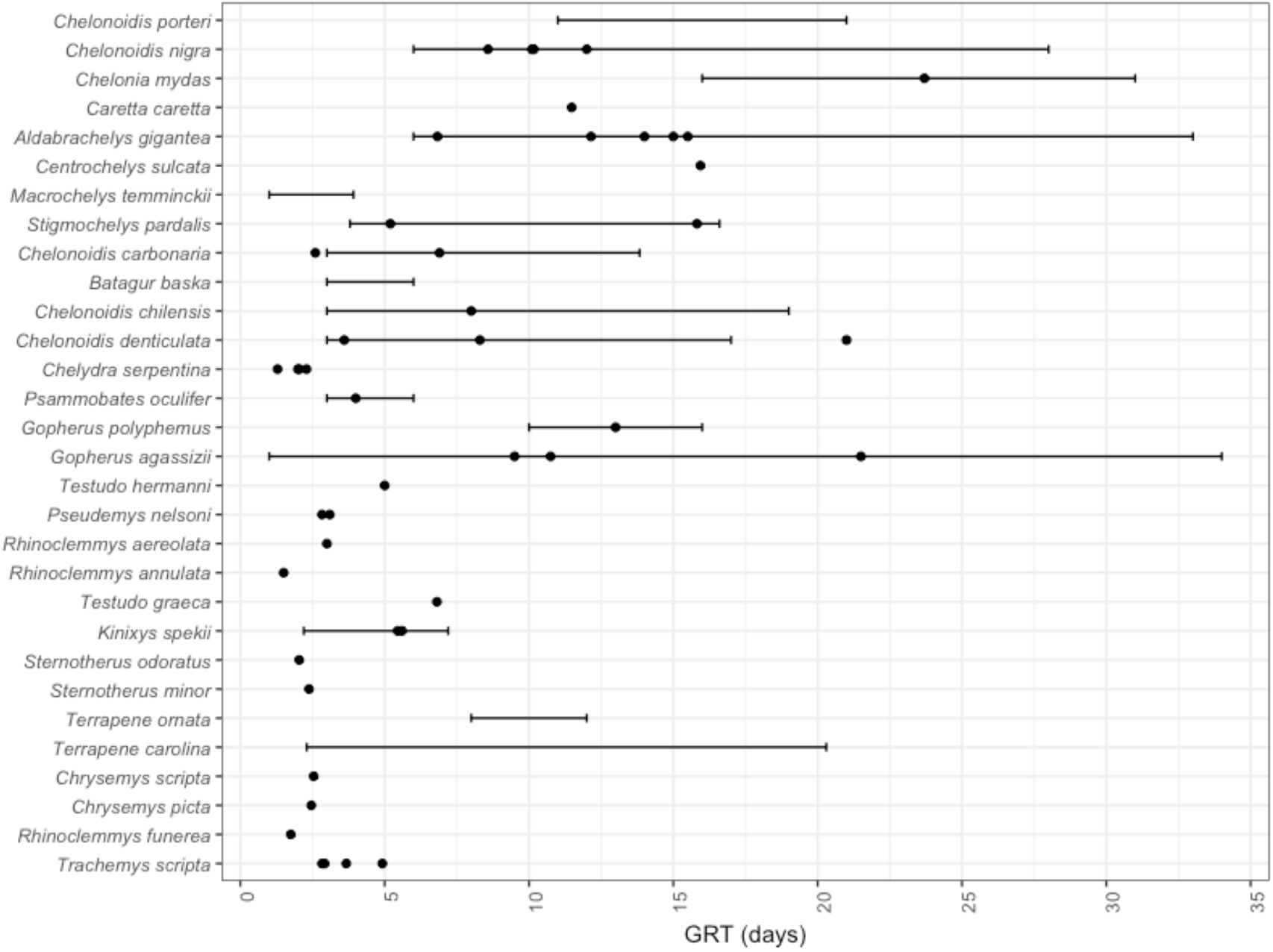
Gut retention times (in days) of 30 species of chelonians. Species are ordered by ascending mean body mass (bottom to top). Points represent the mean gut retention times (GRT) reported for each species by different studies, and bars represent the ranges of GRT reported (minimum and maximum). See Supplementary Materials S3 for references.

Overall, mean GRT seems to increase with species size (Fig. 3), likely due to the increasing length of digestive tracts (Hatt *et al.*, 2002). However, although mean GRT scales with body mass across different tortoise taxa, Franz *et al.* (2011) reported that this relationship was not significant when looking only at tortoises with body mass > 1 kg. The reported effects of chelonian size on GRT varied by species in the studies reviewed. Body size did not influence GRT in the red- and yellow-footed tortoises (Bjorndal, 1989). When comparing GRT of hatchlings with that of adults of the aquatic Florida red-bellied turtle (*Pseudemys nelsoni*), Bjorndal & Bolten (1992) reported that although adults were, on average, 250 times larger, GRT was only 1.4 longer when compared to that of hatchlings.

Potentially muddying the waters, studies on the effect of tortoise size on GRT in Galápagos and Aldabra giant tortoises that used different methods yielded different results. Sadeghayobi *et al.* (2011) found no effect of size on GRT of Galápagos giant tortoises (carapace width range: 0.84–1.53 m) when fed artificial seeds. However, Hatt *et al.* (2002), using n-alkanes particles as GRT markers, reported that mean GRT was shorter for smaller Galápagos giant tortoises (mass range: 7–38 kg vs. 100–210 kg in adults). Similarly, in Aldabra giant tortoises, Falcón *et al.* (in revision) reported no effect of tortoise size (mass range: 0.6–104 kg) on GRT (mean GRT 15 days ± 4) when fed artificial seeds, whereas Waibel *et al.* (2013) reported that sub-adults (20–30 kg) had shorter mean GRT (13 days ± 1) when compared to adult individuals (75–80 kg; 18 days ± 2) when fed fruits of different plants. Thus, other factors such as differences in diet, hydration, food intake and temperature may be more relevant in determining chelonian GRTs within species.

Although seed size can also affect GRT in frugivores (e.g., Fukui, 2003; Figuerola *et al.*, 2010), this does not seem to be the case for chelonians. Braun & Brooks (1987) found that seed size did not influence the GRT of the small, box turtle (*Terrapene carolina*) when fed fruits of different wild plants found in their habitat. Also, in larger chelonians such as the Chaco tortoise (*Chelonoidis chilensis*) (Varela & Bucher, 2002), the Galápagos giant tortoise (Sadeghayobi *et al.*, 2011) and the Aldabra giant tortoise (Falcón *et al.*in revision), seed size does not affect GRT.

Overall, the GRT data suggests that within chelonian species, seeds of different sizes can be dispersed to similar distances.

#### (b) Seed deposition

After being consumed, fruits and seeds are processed in the gut and transported until they are eventually defecated. The state in which seeds are deposited by frugivores is affected by the combination of the mouth and gut treatments. In general, after handling and passage through chelonian guts, seeds are defecated without pulp, but this can be plant-species dependent as some seeds can pass with little physical change and still be covered with pulp (Rick & Bowman, 1961; Varela & Bucher, 2002; Hansen, Kaiser, & Müller, 2008; Waibel *et al.*, 2013). Within the same species of plants, there may be differences in terms of seed damage depending on the species of chelonian that consumes them (Kimmons & Moll, 2010).

Damage to seeds tends to be minimal after defecation. For example, Rick & Bowman (1961) found that less than 1% of recovered seeds of *Solanum cheesmaniae*(Solanaceae) showed any signs of damage after gut passage. Similarly, virtually all the seeds of *Solanum aldabrense* were recovered intact from a single Aldabra giant tortoise scat (WF, pers. obs.). Also, painted turtles (*Chrysemys picta*) pass 99% of seeds intact (Padgett, Carboni, & Schepis, 2010). In addition, 90% of gut-passed seeds were intact for *Chelonoidis carbonarius* in the Pantanal (Wang *et al.*, 2011). Moreover, most seeds were intact for after gut passage in *C. denticulatus* in the Brazilian Amazonia (Jerozolimski *et al.*, 2009). Even for soft seeds without endocarp, like *Syzygium mammilatum* (Myrtaceae), substantial amounts of seeds survive gut passage undamaged (Hansen *et al.*, 2008). As a result of the minimal damage experienced by seeds after chelonian gut passage, many of them remain viable. For example, studies reported between 90–100% of viability of seeds in the faeces of red-footed tortoises (Strong & Fragoso, 2006; Wang *et al.*, 2011).

The location of seed deposition, and perhaps especially the distance from the source, are two key factors for determining what happens to seeds after defecation. This is largely affected by the frugivores’ movement ecology in combination with the GRTs. Only very rarely have chelonian FSD studies specifically included movement ecology (Moll & Jansen, 1995; Strong & Fragoso, 2006; Guzman & Stevenson, 2008; Jerozolimski *et al.*, 2009). We therefore here include information on the movement ecology of chelonians as it affects seed deposition, germination success and ultimately plant recruitment.

Turtles and tortoises have varied home range sizes and movement distances, and these may vary depending on species and individuals within species. There is high variation of home range size between species, with the mean home range size generally increasing with species size (Fig. 4a). Overall, chelonians have a mean home range size of 14.8 ha (± 24.2; n = 41). There is a high within-species variation in home range size (Fig. 4a). Furthermore, chelonians show overall mean daily displacements of 103.9 m day^-1^ (± 114.3; n = 22), but displacement distances do not seem to be related to chelonian size (Fig. 4b). As for home ranges, there is a high variation within species.

**Figure 4:**
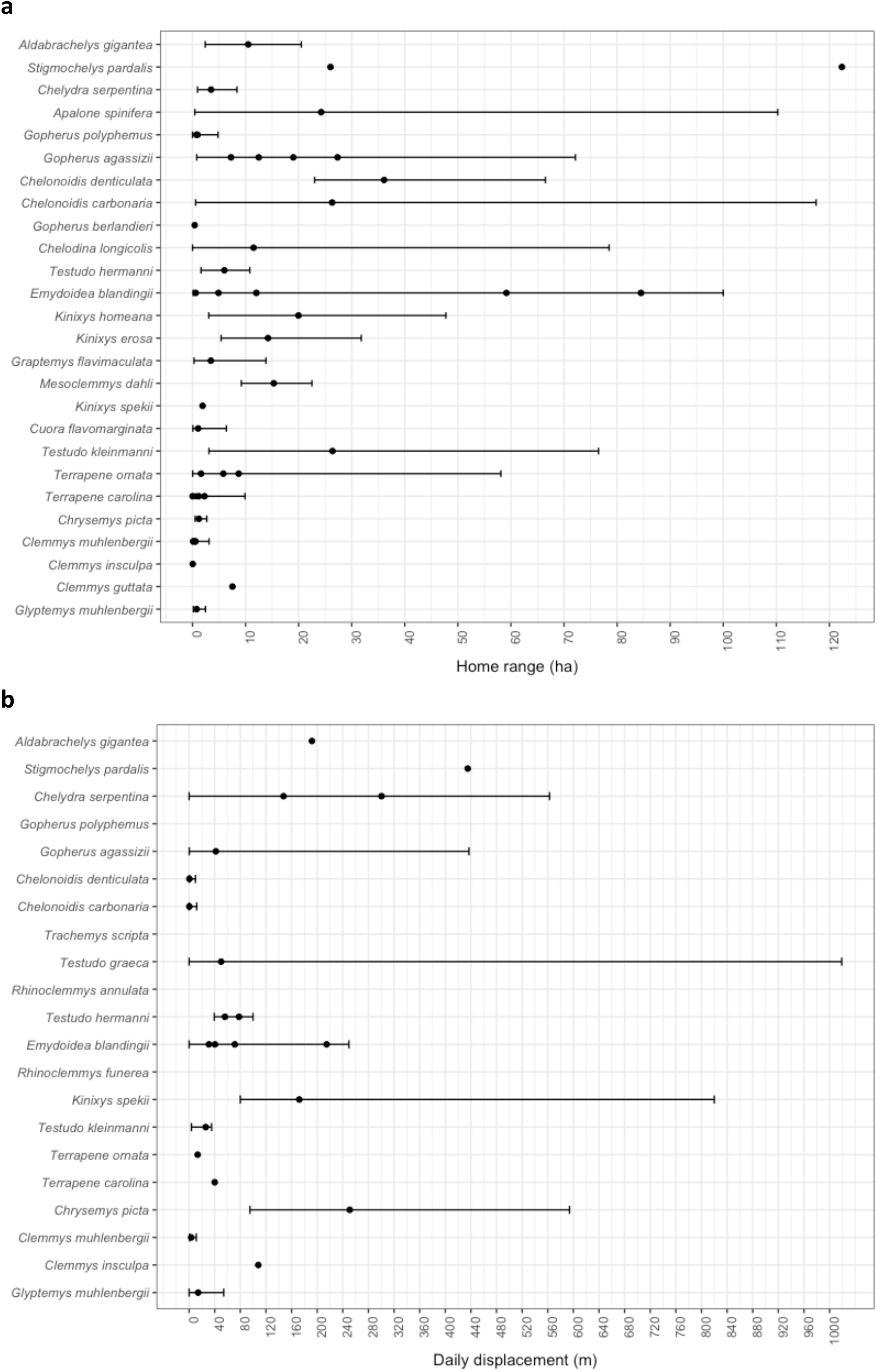
Home ranges (ha) and daily displacement distances (m day^-1^) of certain species of chelonians. Points represent the mean home range and daily displacements reported for each species by different studies, and bars represent the reported ranges (minimum and maximum). See supplementary material S4 (a) and S5 (b) for references.

In contrast to many other frugivores, turtles and tortoises are mostly solitary and thus disperse seeds scattered across the landscape (Varela & Bucher, 2002). Additionally, they often frequent areas expected to be of high recruitment probability for seeds growing into plants. For example, tortoises frequent tree gaps in forested areas to bask in the sun, and such gaps are very suitable recruitment areas for many plant species. A model parameterised with red-footed tortoise cognitive data suggested that the active use of gaps by tortoises enhances the probability of seed deposition in gaps and deforested areas (Soldati, 2015). Indeed, the congeneric yellow-footed tortoise (C. *denticulatus*), which is a major seed disperser, often deposits seed-rich dung in open habitats and treefall gaps (Josseaume 2002, cited in Bonin *et al.*, 2006). In the wild, yellow- and red-footed tortoises favour microsites in open areas that are important for seed germination for resting, such as areas of debris piles, with fallen branches, vines or trees, where they presumably defecate more often than other sites (Moskovits & Bjorndal, 1990; Strong & Fragoso, 2006). Brown wood turtles (*Rhinoclemmys annulata*), are also known to frequent tree gaps (Moll & Jansen, 1995). Open areas are also often used by the gopher tortoise (*Gopherus polyphemus*), which are important areas of plant recruitment in pine savannah in the southestern USA (Birkhead *et al.*, 2005). The European pond turtles, which disperses the seeds of the aquatic waterlily (*Nymphaea alba*, Nymphaceae), effectively disperse seeds between ponds, aiding in maintaining population connectivity and meta-population dynamics of the waterlily (Calviño-Cancela *et al.*, 2007). Moreover, even aquatic species often spend time out of the water, increasing the probability of dispersing plants to suitable habitats (rather than in the water). For example, the black river turtle (*Rhinoclemmys funerea*) in Costa Rica regularly defecates on land (Jansen 1993, cited in Moll & Jansen, 1995).

#### (c) Seed and seedling fate

Seed deposition after zoochory has both spatial and temporal aspects, both of which affect the ultimate fate of seeds. Spatially, the Janzen–Connell model proposed that seeds that are dispersed away from maternal plants have a higher probability of survival as they can escape distance- and density-dependent seed- and seedling predation (Janzen, 1970; Connel, 1971). Both of these are ubiquitous interactions that result in strong establishment limitations for plants (Crawley, 2000; Wright, 2002; Paine & Harms, 2009). Temporally, Guzmán & Stevenson (2011) proposed that escape in time via endozoochory by animals with low metabolic rates and long GRTs, such as chelonians, may aid seeds by basically allowing them to ‘time travel’ into the future to escape from periods with high-intensity seed predation.

After being deposited in suitable habitats, viable seeds that escape predation and pathogens may eventually germinate, and a proportion of these survive and are recruited as adult plants. One of the factors that can affect germination percentage and rates of seeds consumed by chelonians is the gut treatment. For example, gut washing by the digestive fluids of frugivores may be an important mechanism which aids in increasing seed endocarp permeability, and thus enhance germination (Traveset, 1998). Germination percentage and rates can vary within plant genera and between plant species and on the frugivore species after gut passage (reviewed in Traveset, 1998). Effects on seed germination after gut passage can go from positive (enhanced germination), neutral (no effect), to negative (decreased germination). In the studies reviewed here, chelonian gut passage had a mixed effect, depending on the species of chelonian and of fruits/seeds consumed (Table 3). Compared to controls (depulped seeds), 29% of the cases, gut passage had a negative effect on germination, the effect was neutral for 39% of the cases, and in 32% of the cases, seed germination was enhanced.

**Table 2:** Species of chelonians that engage in frugivory and/or seed dispersal.

| **Family** | **Genus** | **Chelonian spp.** |
| --- | --- | --- |
| Carettochelyidae | *Carettochelys* | 1 |
| Chelidae | *Chelodina* | 1 |
| Chelidae | *Elseya* | 2 |
| Chelidae | *Emydura* | 2 |
| Chelidae | *Mesoclemmys* | 2 |
| Chelidae | *Phrynops* | 2 |
| Cheloniidae | *Chelonia* | 1 |
| Chelydridae | *Chelydra* | 1 |
| Chelydridae | *Macrochelys* | 1 |
| Dermatemydidae | *Dermatemys* | 1 |
| Emydidae | *Actinemys* | 1 |
| Emydidae | *Chrysemys* | 1 |
| Emydidae | *Emys* | 2 |
| Emydidae | *Graptemys* | 2 |
| Emydidae | *Malaclemys* | 1 |
| Emydidae | *Terrapene* | 2 |
| Emydidae | *Trachemys* | 2 |
| Geoemydidae | *Batagur* | 2 |
| Geoemydidae | *Heosemys* | 1 |
| Geoemydidae | *Rhinoclemmys* | 3 |
| Geoemydidae | *Vijayachelys* | 1 |
| Kinosternidae | *Kinosternon* | 5 |
| Kinosternidae | *Staurotypus* | 1 |
| Kinosternidae | *Sternotherus* | 1 |
| Platysternidae | *Platysternon* | 1 |
| Podocnemididae | *Peltocephalus* | 1 |
| Podocnemididae | *Podocnemis* | 3 |
| Testudinidae | *Aldabrachelys* | 1 |
| Testudinidae | *Astrochelys* | 1 |
| Testudinidae | *Chelonoidis* | 5 |
| Testudinidae | *Chersina* | 1 |
| Testudinidae | *Gopherus* | 3 |
| Testudinidae | *Homopus* | 1 |
| Testudinidae | *Indotestudo* | 3 |
| Testudinidae | *Kinixys* | 1 |
| Testudinidae | *Psammobates* | 1 |
| Testudinidae | *Pyxis* | 1 |
| Testudinidae | *Stigmochelys* | 1 |
| Testudinidae | *Testudo* | 2 |
| Trionychidae | *Apalone* | 1 |
| Trionychidae | *Trionyx* | 1 |

**Table 3:**
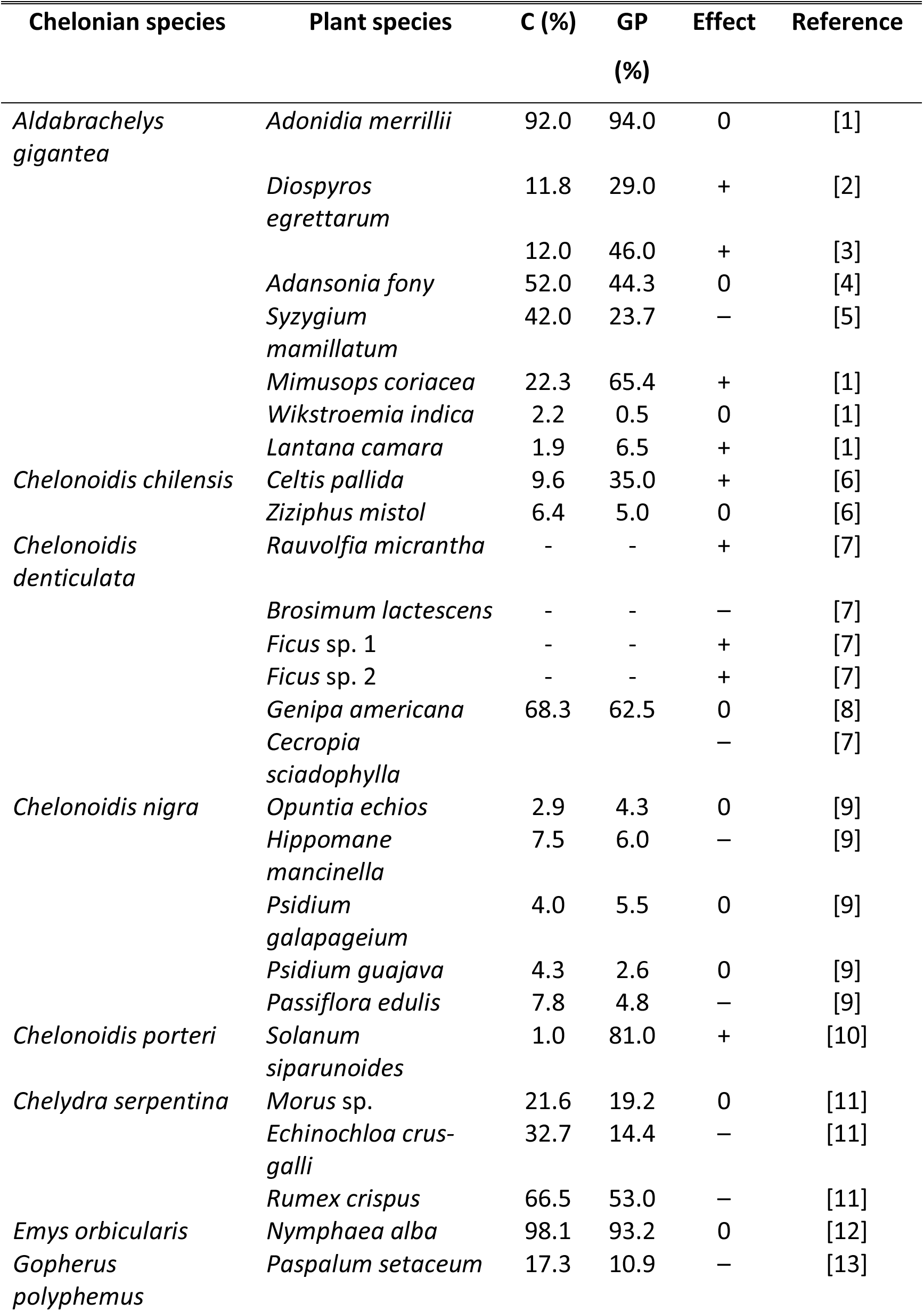
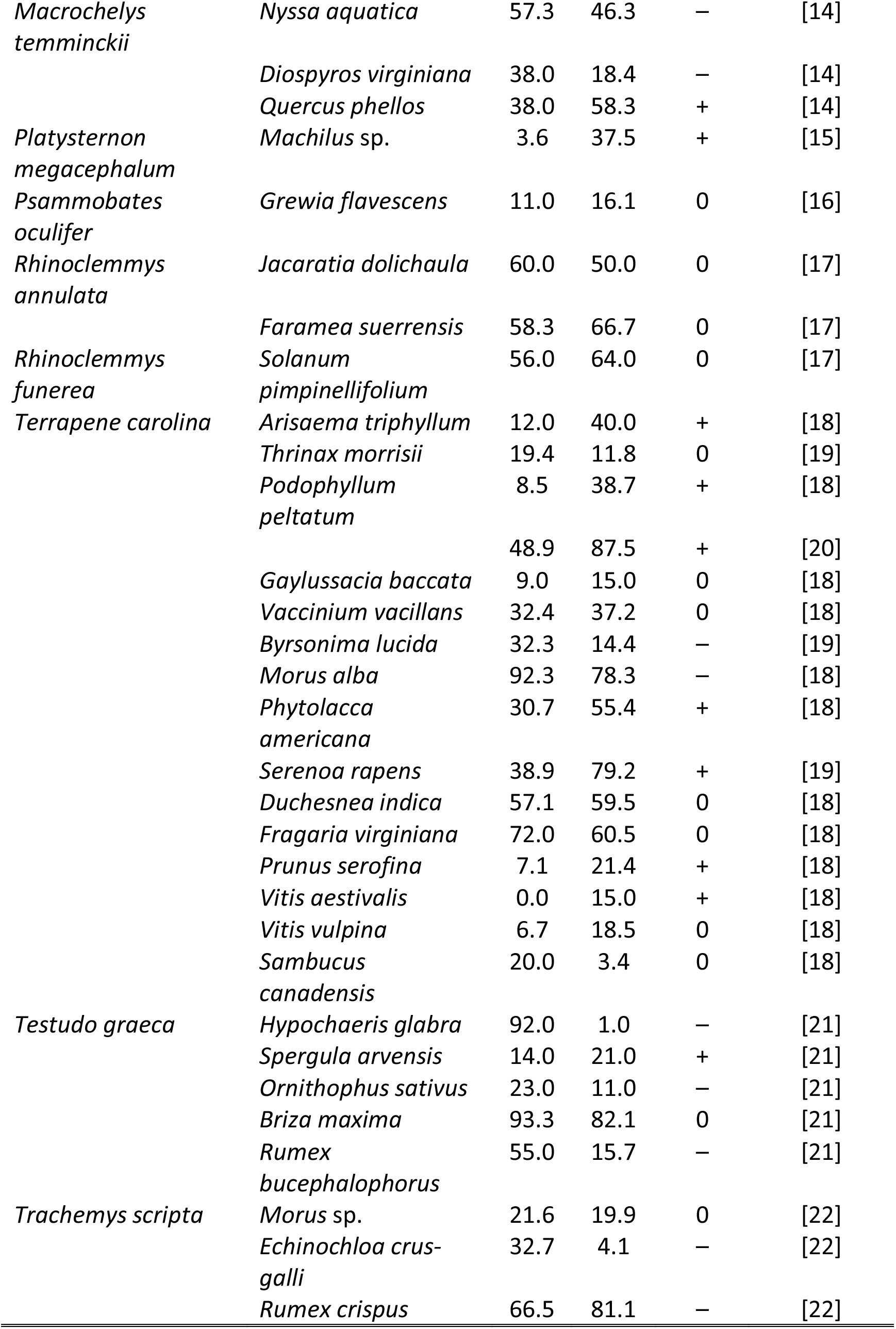
Effects of chelonian gut passage on the germination percent of different plant species. Effects, compared to controls, can go from positive (+) for enhanced germination, neutral (0) to negative (−). Chelonian species are ordered alphabetically. Treatments are depicted as gut passage (GP) and controls (C). Only control treatments of depulped seeds are considered here. See Supplementary Materials S6 for references.

In addition to depending on the species of chelonians and plants, factors such as chelonian ontogeny, seed size, within-species variation in seed dormancy, and external stimuli may affect seed germination. For example, tortoise age, which correlates with size, can affect the likelihood of seed germination after passage through the guts of Aldabra giant tortoises (*Aldabrachelys gigantea*), with smaller sub-adults increasing the probability of germination of some plant species when compared to larger adult tortoises, and this was attributed to the shorter GRTs of sub-adults (Waibel *et al.*, 2013). Braun & Brooks (1987) found that after gut passage through the box turtle (*Terrapene carolina*), seed germination increased with increasing seed size. Plant species may also have different degrees of seed dormancy that may affect seed germination after gut passage (Rick & Bowman, 1961). External stimuli, such as the availability of light has been shown to have a differential effect on aquatic seed germination, with delayed germination after gut passage in light conditions (but with equal total germination to controls), and delayed germination during the first year, with subsequent increased germination speed and percentage in the long term in dark conditions (Calviño-Cancela *et al.*, 2007). The authors suggest that in their natural habitat, the differential effect of gut passage in combination with light stimuli is expected to affect seed germination in turbid vs. clear bodies of water. Similarly, we can expect that seeds inside the dung of terrestrial species, with no direct light, to have a delayed germination, as dung disintegrates, and thus escape predators in time (assuming that the dung does not attract predators).

In terms of seedling growth and vigour, the few studies we found reported a positive effect of chelonian gut passage. For example, in the case of *Syzygium mammilatum* (Myrtaceae), gut passage through *A. gigantea* had negative effects on seed germination rate, but positive effects on seedling growth and health when grown ‘in shitu’ (i.e., grown in scat) (Hansen *et al.*, 2008). In addition, Elbers & Moll (2011) found that common persimmon (*Diospyros virginiana*, Ebenaceae) and water tupelo (*Nyssa aquatica*, Nyssaceae) seeds had lower proportions of germinating seeds (compared to controls) after passage through the guts of alligator snapping turtles (*Macrochelys temminckii*), while the acorns of the willow oak (*Quercus phellos*, Fagaceae) had a higher proportion of germination after gut passage compared to controls. Passage of seeds of the grass *Briza maxima* (Poaceae) through the gut of the spur-thighed tortoise (*Testudo graeca*) led to seedlings growing larger and faster, although this may have been due to filtering of seed size, as only larger seeds were recorded passing through the gut (Cobo & Andreu, 1988).

#### (d) Secondary seed dispersal

Secondary seed dispersal is the process by which seeds that have been initially dispersed by a frugivore via endozoochory are consumed by a second disperser, for example, through coprophagy. Some chelonian species have been observed acting as potential secondary seed dispersers. For example, giant tortoises frequently eat each other’s scat on Aldabra Atoll (WF & DMH, pers. obs.), and red- and yellow-footed tortoises (*Chelonoidis carbonarius* and *C. denticulatus*, respectively) have been observed eating tortoise scat in Brazil (Moskovits and Bjorndal 1990). Also, Young (2003), states that tortoises (without specifying which species) are partial to eating dung from camels, sheep, and goats, who themselves are potential seed dispersers (Kuiters & Huiskes, 2010; Mancilla-Leytón, Fernández-Alés, & Vicente, 2011; Root-Bernstein & Svenning, 2016). Juvenile Central American river turtles (*Dermatemys mawii*) eat the scat of adults, presumably to obtain cellulolytic bacteria to aid in digestion of plant foods (Legler & Vogt, 2013). Forsten’s tortoise (*Indotestudo forsteni*) has been observed eating monkey scat, which contained fruit pulp (Ives *et al.*, 2008), and thus likely also contained seeds. In addition, deer faecal pellets were found in the scat of the box turtle (*Terrapene carolina bauri*) (Platt *et al.*, 2009). Although it is possible that they secondarily ingested some seeds from the deer scat, the authors stated that the contribution to the overall number of seeds found in the turtle’s dung is likely to be minimal. Also, North American box turtles (*Terrapene carolina* and *T. ornata*) regularly eat cow dung which often contains seeds (DM, pers. obs.)

Seeds in tortoise scat can also be potentially secondarily dispersed by non-chelonian species. For example, turtle doves (*Streptopelia picturata*) have been observed eating the contents of giant tortoise scat on Aldabra Atoll (WF, pers. obs). Moreover, dung beetles, which feed on scat and usually bury it, have been recorded amassing and dispersing scat of red- and yellow-footed tortoises in Brazil (Strong & Fragoso, 2006). In addition, land crabs (*Cardisoma carnifex*) and coconut crabs (*Birgus lastro*) have been observed eating giant tortoise scat containing grass and *Ficus* sp. seeds on Aldabra Atoll (WF, pers. obs.). We are unaware of any studies addressing the effects of secondary seed dispersal by chelonians, or other species consuming chelonian scat, on plant germination and/or recruitment and it thus remains to be seen whether effective secondary seed dispersal occurs in, or is promoted by, chelonians.

## VI Chelonian FSD in a community context

The interactions between plants and frugivores do not occur in a vacuum, but are embedded in the ecological network of seed dispersal interactions between all plant species and all frugivores in the community (Bascompte & Jordano, 2007).

Therefore, if we truly want to know the role of chelonians as seed dispersers, we must look at their role in a community context. Studies on the role of chelonians as frugivores and seed dispersers at the community level are scarce (we found only four studies, described below) yet they provide valuable insights about their role in relation to other frugivores.

Donatti *et al.* (2011) studied seed dispersal interactions at the community level in the Brazilian Pantanal using bipartite interaction network analysis. For mutualistic plant–animal interactions, a bipartite network consists of nodes (vertices) and links (edges), which are represented by trophic levels (i.e., frugivores and plants in this case) and the interactions between them (interactions within trophic levels are not possible). The Pantanal seed dispersal network was hyper-diverse, with 46 species of frugivores interacting with 46 species of plants. In the network, the red-footed tortoise (*Chelonoidis carbonarius*), was the sixth most important frugivore in terms of the number of interactions (Fig. 5a). Given the diversity and the complexity of the network, based on the number of interactions in comparison to other frugivores and the fact that they are capable of dispersing large-seeded plants, red-footed tortoises are probably one of the most important dispersers in the Pantanal community.

**Figure 5:**
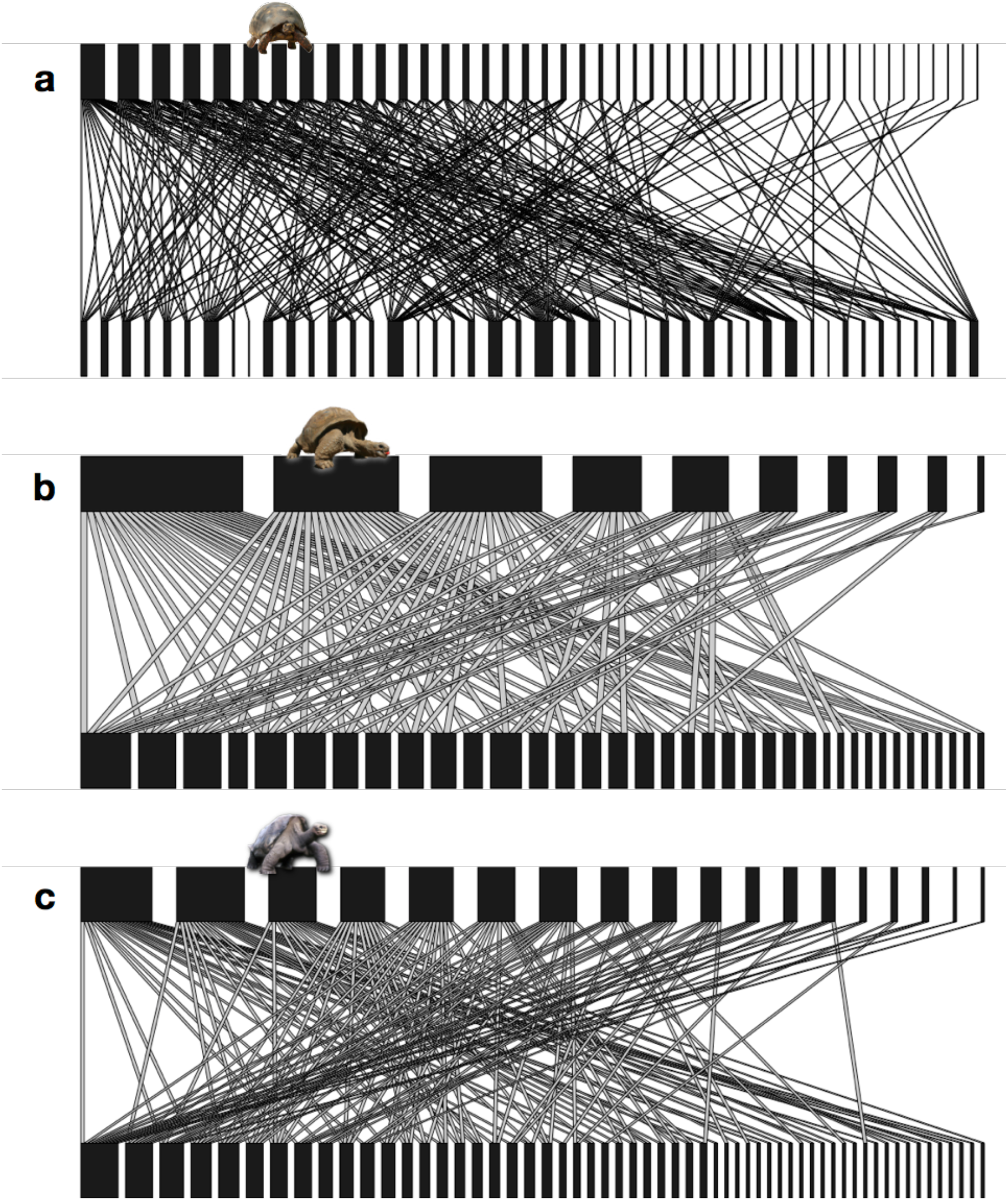
The role of chelonians as frugivores and seed dispersers in a community context, based on the seed dispersal networks of Pantanal (a; *Chelonoidis carbonaria*), Aldabra Atoll (b; *Aldabrachelys gigantea*), and Galápagos (c; *Chelonoidis nigra*). Networks are qualitative (i.e., the strength of the interactions are not considered) and the size of the boxes represent the number of interactions for each frugivore (top; organised from largest to smallest) and each plant (bottom) present in the community. Networks drawn from data available in Donatti *et al.* (2011; a), Falcón (2018; b), and Heleno *et al.* (2013; c).

Falcón (2018) studied seed dispersal interactions in the smaller plant–frugivore community of Aldabra Atoll (with ten frugivores and 37 plant species), home to Aldabra giant tortoises (*Aldabrachelys gigantea*). The network was highly generalised, and tortoises were the second most important seed dispersers in terms of the number of interactions. In total, *A. gigantea* dispersed the seeds of at least 20 fleshy-fruited plant species (grasses and sedges were not included; Fig. 5b), including large-seeded ones such as *Cordia subcordata* (Boraginaceae) and *Guettarda speciosa*(Rubiaceae). Moreover, he found that the network was most vulnerable to the loss of three particular frugivores, one of them being the giant tortoises. This study highlighted the importance of tortoises as megafaunal seed dispersers and suggests that the many recently extinct giant tortoises in the Indian Ocean (see Hansen *et al.* 2010) had a similarly pivotal role in their communities before being exterminated.

In Galápagos, Heleno *et al.* (2013) used network analysis to investigate the impact of alien plants on the seed dispersal networks in two islands, one of which harboured giant tortoises (*Chelonoidis nigrus*). They looked at the seed dispersal of both native and introduced plants by the different island frugivores. Giant tortoises here were the third most important seed disperser in terms of the number of interactions; Fig. 5c), and were especially important for fleshy-fruited plants. They also performed an analysis of the quantitative seed dispersal network, and stated that tortoises played an important role as seed dispersers based on the strength of interactions, and that the extirpation of tortoises on other islands in the Galápagos must have resulted in a negative impact on seed dispersal function at the community level.

Also in the Galápagos, Nogales *et al.* (2017) took a step further and studied the direct contributions delivered by different groups of frugivores, including giant tortoises, lizards, and three groups of birds, to the number of seeds dispersed, and the effect on germination. Frequency of occurrence of seeds was the highest in the scats of giant tortoises and medium-sized passerine birds, but the number of seed deposited per unit area was lowest for tortoises and lizards. In terms of seed emergence after gut passage, only a small proportion of seeds from all scat samples germinated (19%) within the study period, but those that originated from tortoise scat showed the highest emergence frequency compared to seeds dispersed by all the other disperser guilds. Based on the large frequency of occurrence and number of seeds found in the scat, as well as seed germination after gut passage, they concluded that Galápagos giant tortoises play a key role as seed dispersers in the Galápagos Islands.

## VII Chelonians as megafaunal seed dispersers

On many islands worldwide, large and giant tortoises were present until recently, and were often the largest vertebrates in their respective faunas (Hansen *et al.*, 2010). Giant tortoises on islands function as megafauna, capable of dispersing even very large seeds (Hansen & Galetti, 2009). Surprisingly, there is evidence that medium-sized tortoises in continental ecosystems can disperse unexpectedly large seeds. In Amazonia, (Jerozolimski *et al.*, 2009) found that yellow-footed tortoises (*Chelonoidis denticulatus*), a tortoise with a mean length of 40 cm, dispersed seeds of the palm *Attalea maripa* (Arecaceae) of up to 40 × 17 mm, and (Mitchell & Daly, 1998) described how *C. denticulatus* tortoises easily swallowed the 50–60 mm large fruits of *Spondias testudinis* (Anacardiaceae), thus presumably capable of dispersing the ca. 40 × 30 mm large seeds. The two Brazilian *Chelonoidis* species may thus act as some of the last surviving heirs to several of the many large-seeded fruits left orphaned by late Pleistocene megafauna extinctions (Guimarães, Galetti, & Jordano, 2008), and *Spondias mombin* is thus perhaps not yet entirely “culturally deprived in [mammalian] megafauna-free forest”(*sensu* Janzen, 1985).

## VIII Chelonian FSD and conservation/restoration

Chelonians are the most endangered of the major groups of vertebrates, exceeding birds, mammals, fishes and amphibians (van Dijk *et al.*, 2014). Factors that affect the conservation of chelonians include habitat destruction, exploitation, and climate change. On a more positive note, chelonians have shown themselves to be key players in habitat restoration projects.

### (1) Chelonian conservation

Roll *et al.* (2017) found that the distributional overlap of the range of chelonians with protected areas is only ca. 10%, which puts them at great risk, especially if they are habitat specialists. For example, the Northern Australian snapping turtle (*Elseya dentata*)resides in riverine habitats, and their diet consist mainly of fruits of riparian rainforest trees, so they are particularly vulnerable to changes in land management that may have negative effects on riparian forest habitats (Kennett & Tory, 1996). Thus, habitat modification and destruction not only affect chelonian populations, but can also affect the availability of fruit resources, which can lead to the loss of seed dispersal mutualisms.

Exploitation is another factor threatening the conservation of chelonian species, and the main causes are consumption as food resources, traditional medicine, and the pet trade. Known frugivorous chelonians are not exempt from suffering from exploitation, and for example, species of the turtle genera *Trachemys* and *Pseudemys* are the most exported turtles in the USA, with individuals being taken directly from the wild, or taken from the wild and subsequently bred in captivity (Moll & Moll, 2004; Mali *et al.*, 2014). Similarly, species such as the radiated tortoise (*Astrochelys radiata*) (Leuteritz, 2003) and the spur-thighed tortoise (*Testudo graeca*), are prone to exploitation from their native habitat (Walker & Rafeliarisoa, 2012). Exploitation of chelonian populations may have important implications for seed dispersal as the reduction of frugivore populations can result in the functional extinction of seed dispersal mutualisms, even before the species of frugivore itself goes extinct (e.g., McConkey & Drake, 2006).

In addition, changes in temperature and precipitation due to anthropogenically-induced climate change are poised to affect many ectothermic species, including chelonians, harder than endothermic ones (Walther *et al.*, 2002; Deutsch *et al.*, 2008; Clusella-Trullas, Blackburn, & Chown, 2011; Ihlow *et al.*, 2012). For example, turtle and tortoise species may respond strongly to precipitation, and their activity and movements decrease with increasingly dry periods (Luiselli, 2005; Baxter, 2015; Falcón *et al.*, 2018). In addition, increasing droughts can affect the habitats of chelonians (Haverkamp *et al.*, 2017), and potentially reduce shade availability, which is an important resource for thermoregulation (Merton *et al.*,1976; Moulherat *et al.*, 2014). Moreover, increasing temperatures have been shown to decrease the activity of chelonians, and they may be particularly vulnerable to increases in air temperature in terms of thermoregulation (Lambert, 1981; McMaster & Downs, 2013; Falcón *et al.*, 2018). Thus, the magnitude and outcome of chelonian FSD is very likely to be negatively affected by climate change.

### (2) Rewilding and restoration

Overall, because frugivorous chelonians in general are efficient seed dispersers they are ideal candidates for rewilding and restoration efforts that have the resurrection of extinct seed dispersal interactions as a major focus. This is especially the case in island ecosystems, where many of the large-bodied frugivores have gone extinct (Heinen *et al.*, 2017), and where giant tortoises are in general considered to be ecosystem engineers (Hansen *et al.*, 2010). The best-studied example of this is the introduction of Aldabra giant tortoises to islands in the Mascarenes to restore the function left behind by the extinction of the endemic *Cylindrapsis* giant tortoises (Griffiths *et al.*, 2010; Hansen *et al.*, 2010). Here, they effectively disperse the seeds of several endemic and endangered plant species (Hansen *et al.*, 2008; Griffiths *et al.*, 2010), including the large-seeded *Diospyros egrettarum* (Ebenaceae) (Griffiths *et al.*, 2011). Moreover, these tortoises have also been shown to have potential as seed dispersers of the huge fruits of Baobab trees (*Adansonia rubrostipa*, Malvaceae) in Madagascar, where giant tortoises also used to occur (Andriantsaralaza *et al.*, 2013), and may soon find themselves being deployed as ecological restoration agents in Madagascar, too (Pedrono *et al.*, 2017).

The perhaps most ‘extreme’ functional substitution can be found in Hawai’i, where recently extinct herbivorous and frugivorous giant flightless ducks and geese have been replaced by the large African spurred tortoise (*Centrochelys sulcata*) in the Makauwahi Cave Reserve on the island of Kauai (Burney *et al.*, 2012). Although neither terrestrial nor fresh water chelonians ever reached Hawai’i by natural means, based on their ecology, the authors posited that the spurred tortoises could act as ecological substitutes for the extinct endemic frugivore-herbivores.

Rewilding with tortoises does not have to be necessarily limited to islands, and according to Sobral-Souza *et al.* (2017), the continental northern Atlantic Forest of Brazil, which is heavily defaunated and fragmented, and whose fragments are too small to reintroduce large mammalian frugivores, is another potential tortoise rewilding region. Based on studies highlighting the role of yellow- and red-footed tortoises as seed dispersers, especially for large-seeded plants (*Chelonoidis denticulatus* and *C. carbonarius*, respectively), and on the success of rewilding efforts with Aldabra giant tortoises, the authors argued that introducing these *Chelonoidis* spp. in fragments of the northern Atlantic Forest would be a way to mitigate the negative cascading effects of defaunation. To support their argument, they employed niche modelling based on known occurrence of tortoises, and assessed food availability and conservation co-benefits, and found that fragments in the northern Atlantic Forest are suitable for these tortoises.

## X Conclusions

1. Chelonian FSD is geographically and taxonomically widespread. In contrast to other major classes of frugivorous reptiles, chelonian FSD is not mainly restricted to islands. However, and different to patterns of chelonian species richness, most FSD studies in turtles and tortoises come from the south-eastern USA and northern South America. Studies on chelonian FSD in south-east Asia, where chelonian species richness peaks, are notably scarce.
2. Likewise, chelonian FSD occurs widely across the angiosperm phylogeny, with at least one family represented in the major grades and clades. There is, however, an asymmetry of interactions, in which few plant families amass most of the unique pairwise interactions with chelonians.
3. Based on the studies reviewed here, we expect frugivorous chelonians to be, in most cases, efficient seed dispersers. Not only they can consume large quantities and a high diversity of fruit and/or seed species, but also damage by the mouth parts or after passage is minimal, resulting in many viable seeds. Moreover, compared to controls, passage of seeds through chelonian guts seldom causes negative impacts on seed germination, often resulting in neutral to positive effects, and can result in high seedling vigour.
4. Seed dispersal interactions do not occur in a vacuum, and the few studies that have investigated the role of chelonians from a community perspective have highlighted their importance in terms of not only the number and strength of interactions, but also the importance of their role as central species amongst frugivores in seed dispersal networks.
5. Large and giant tortoises (Testudinidae) were present on many islands worldwide, and were often amongst the largest vertebrates. It is in islands, especially, where they are/were prime dispersers of large-seeded plants. Nonetheless, the capacity of large testudinid species in continental ecosystems as megafaunal seed dispersers has also been demonstrated. Therefore, chelonians can act as megafaunal seed dispersers.
6. Finally, on the one hand, chelonians are amongst the most threatened taxa in the world. Not only they suffer from habitat loss and lack of protection, but they are also heavily exploited, and face an uncertain future due to pressures imposed by climate change. On the other hand, chelonians have a great potential to aid in the conservation of plant–frugivore mutualisms, which have vital implications for ecosystem functioning, and to be used as analogue species to restore lost interactions and functions.

## XI Acknowledgements

We thank Dr. Yuval Itescu (Tel Aviv University) for providing us with the data on the global distribution of chelonians. Funding was provided to DMH and WF by the Swiss National Science Foundation (grant number 31003A_143940), and the Zoological Museum and the Department of Evolutionary Biology and Environmental Studies (both at the University of Zurich), for which we are thankful. DM would like to thank Missouri State University, and especially the Biology Department, for financial and other support of research he has participated in related to frugivory and seed dispersal in chelonians.

## SUPPLEMENTARY MATERIALS

**S1:**
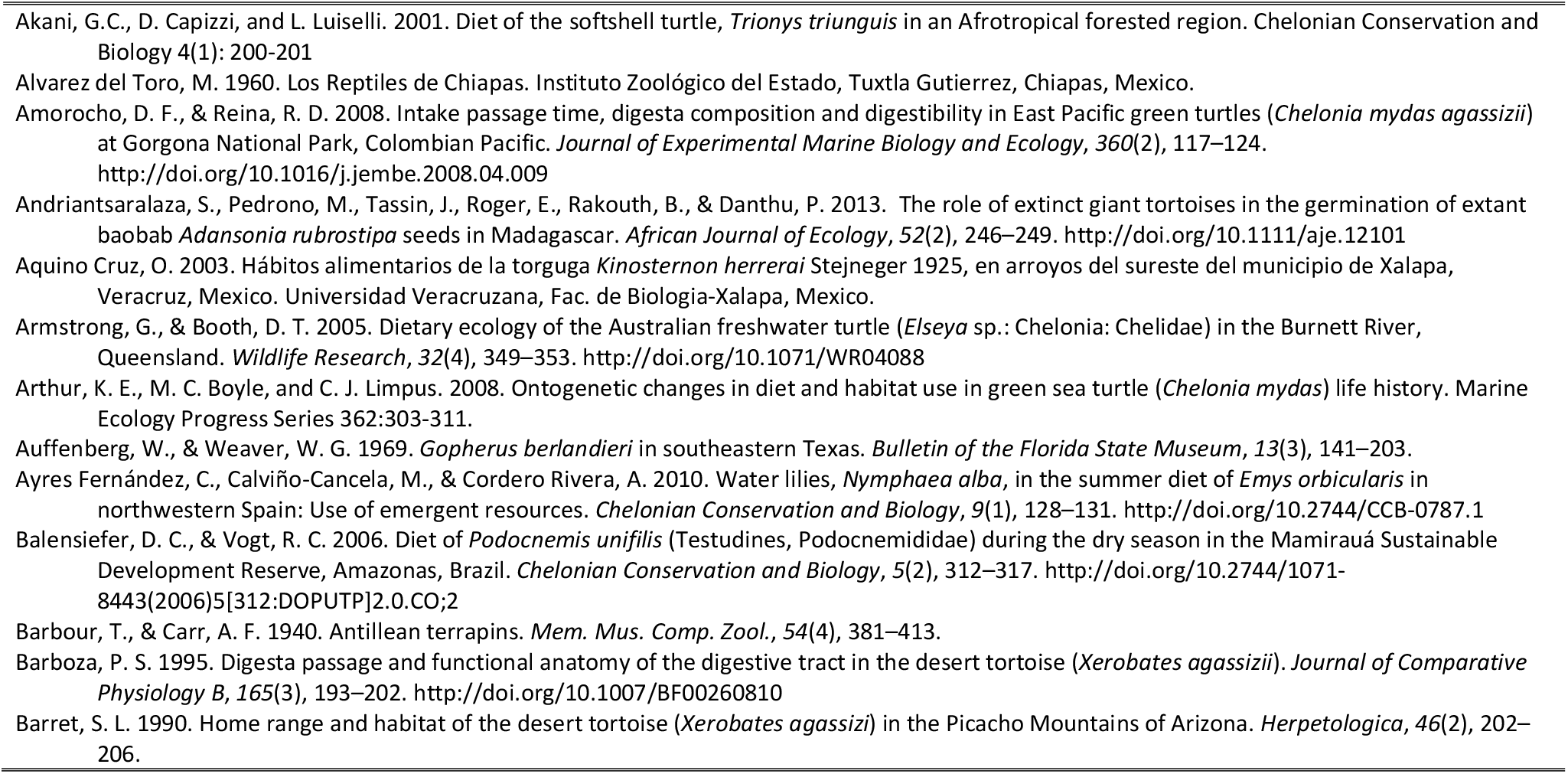
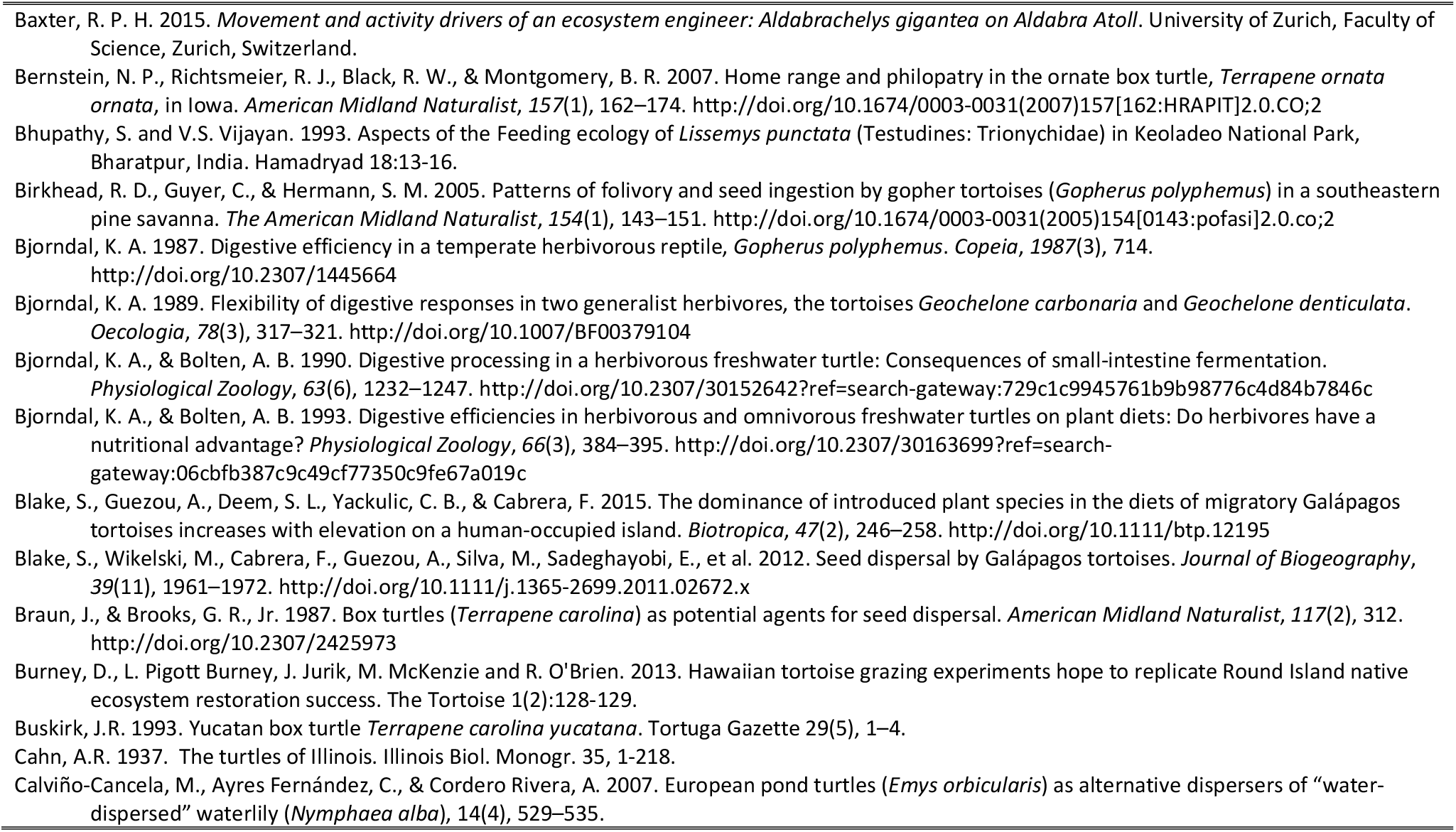
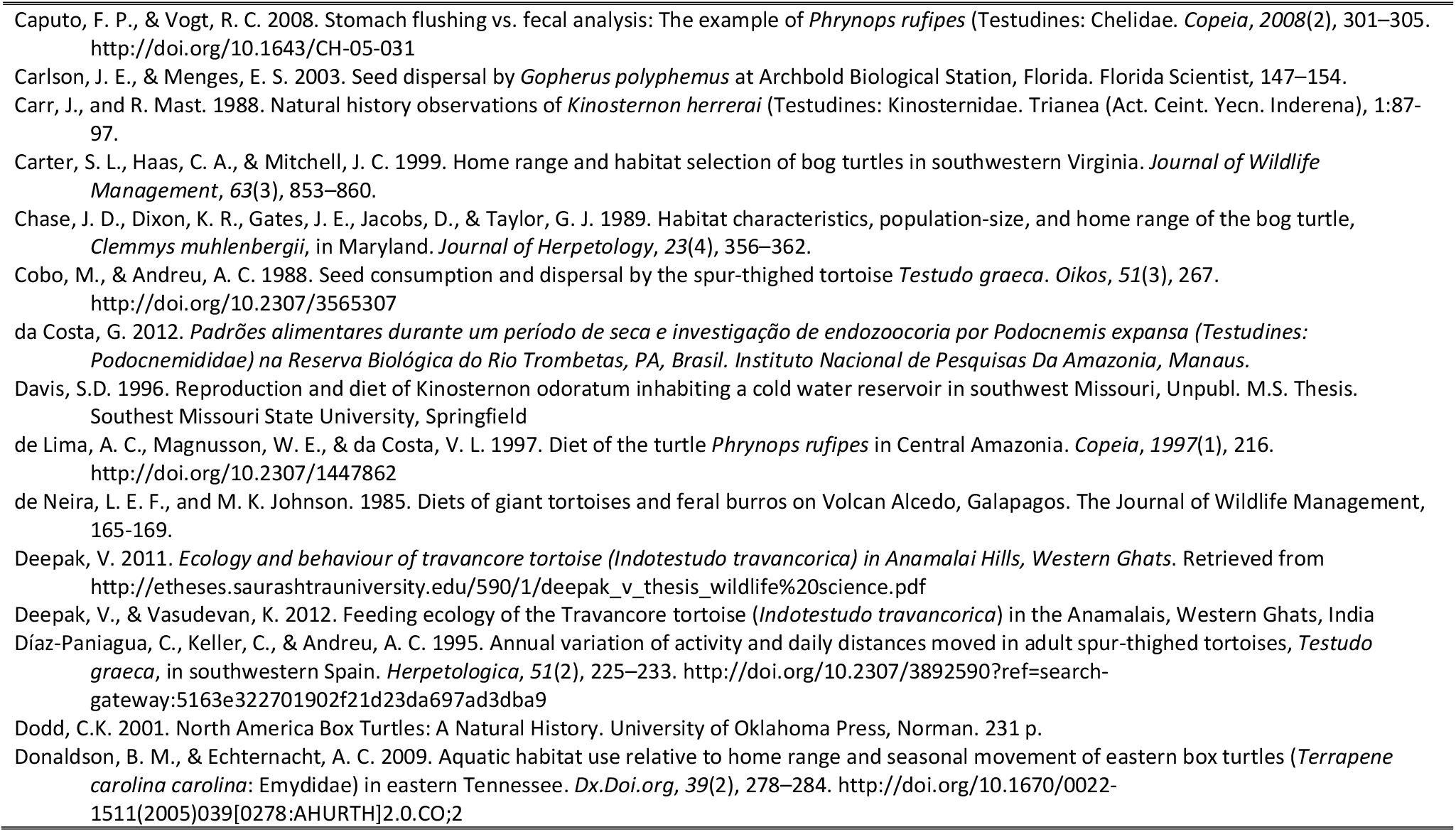
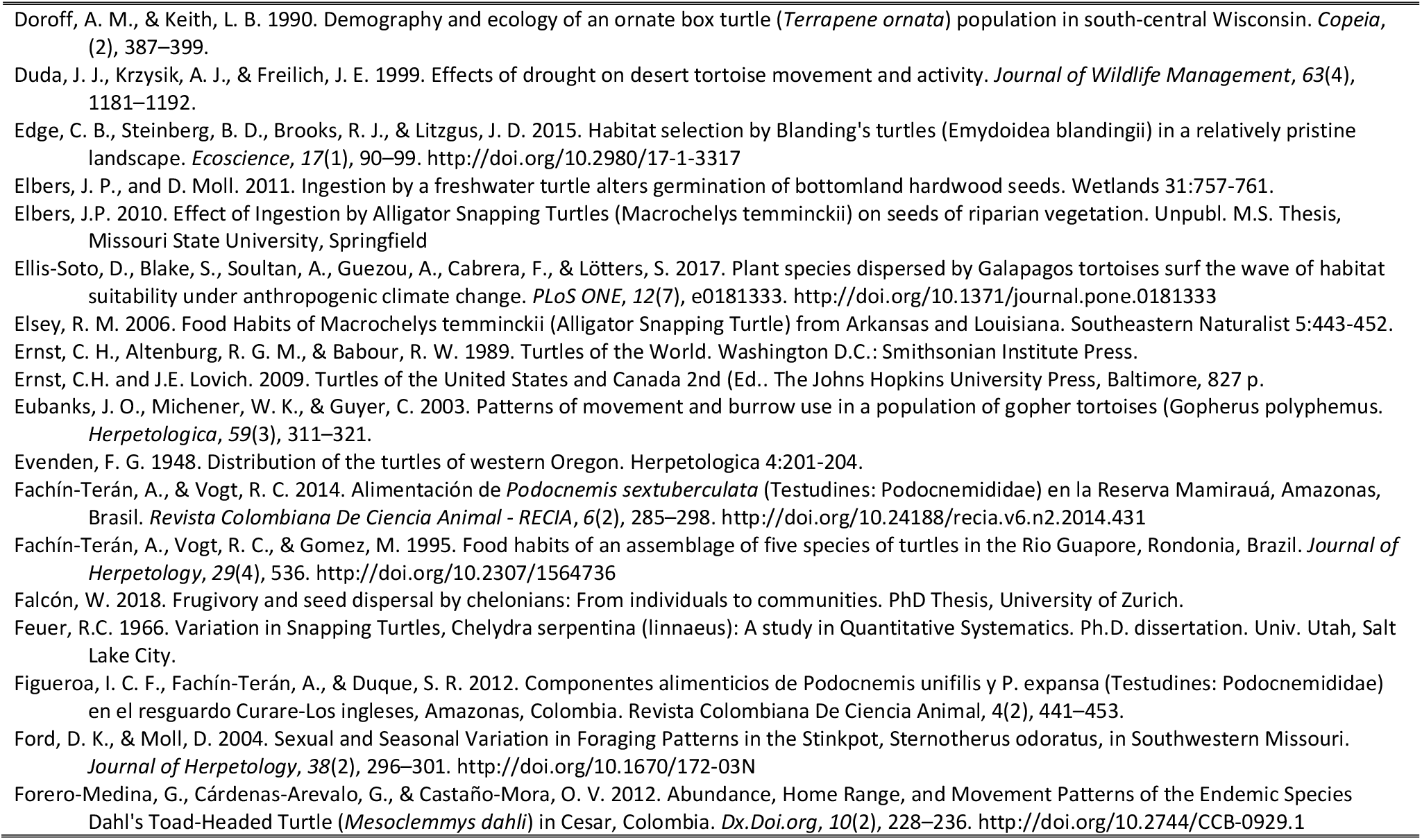
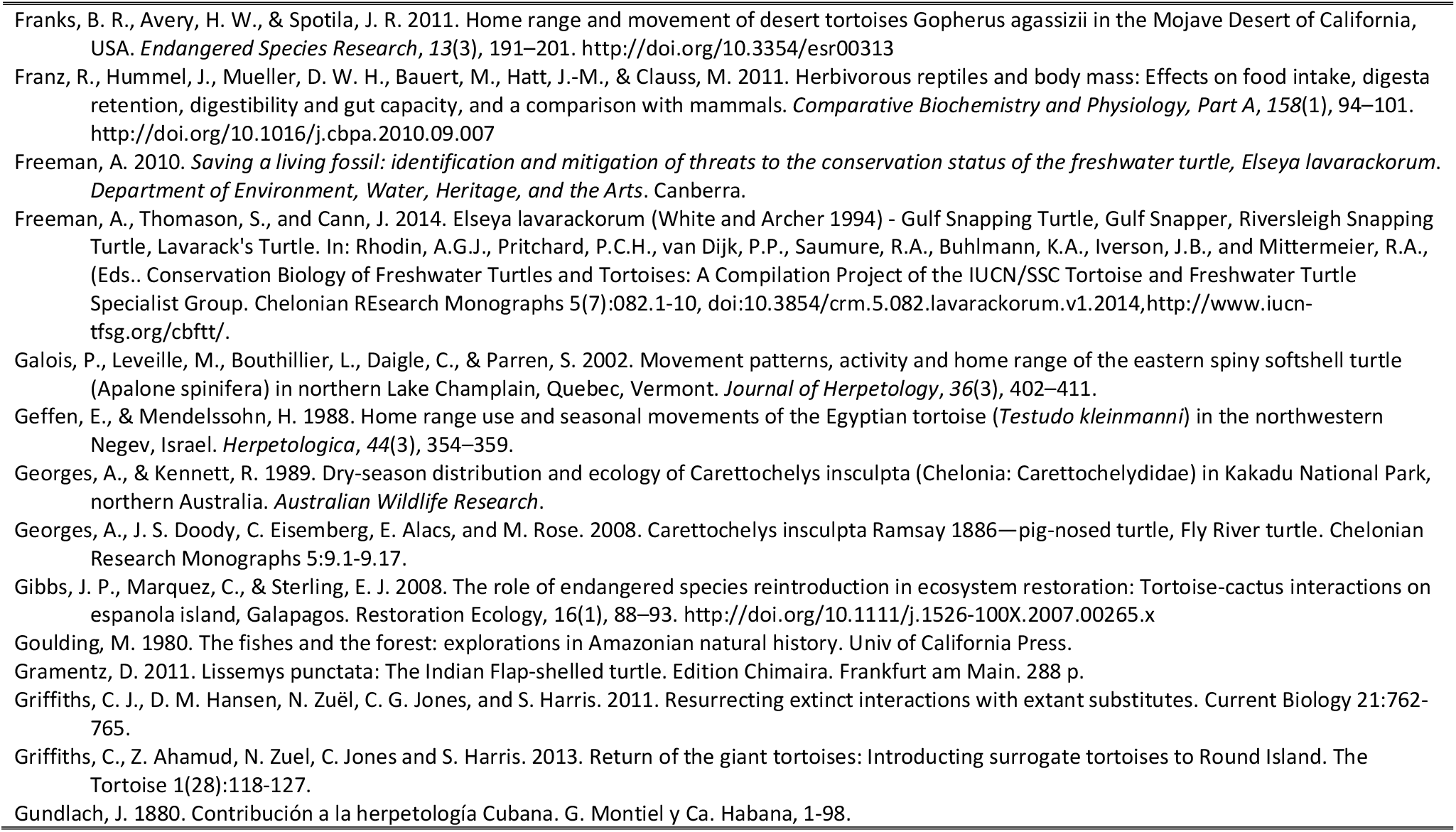
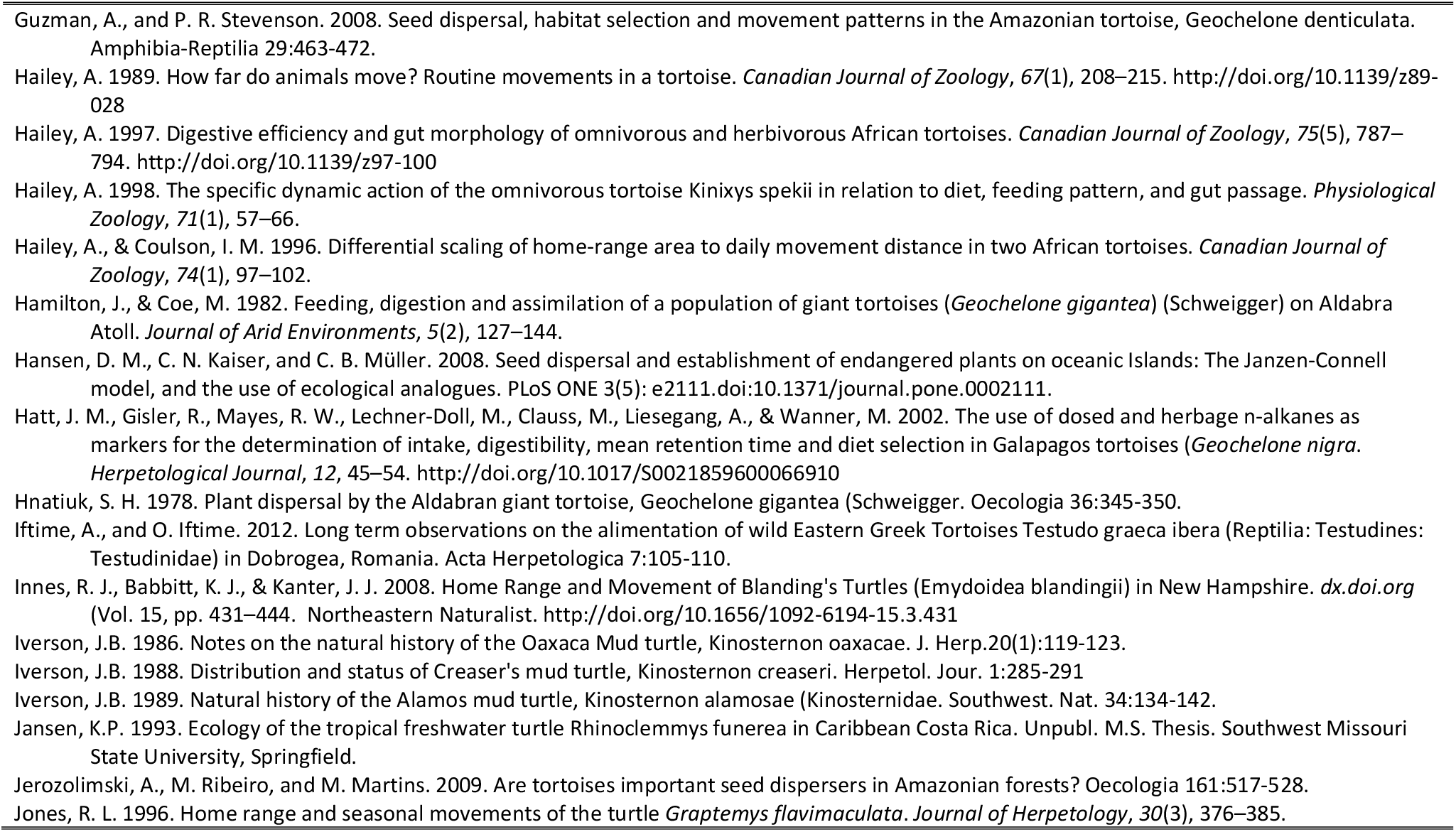
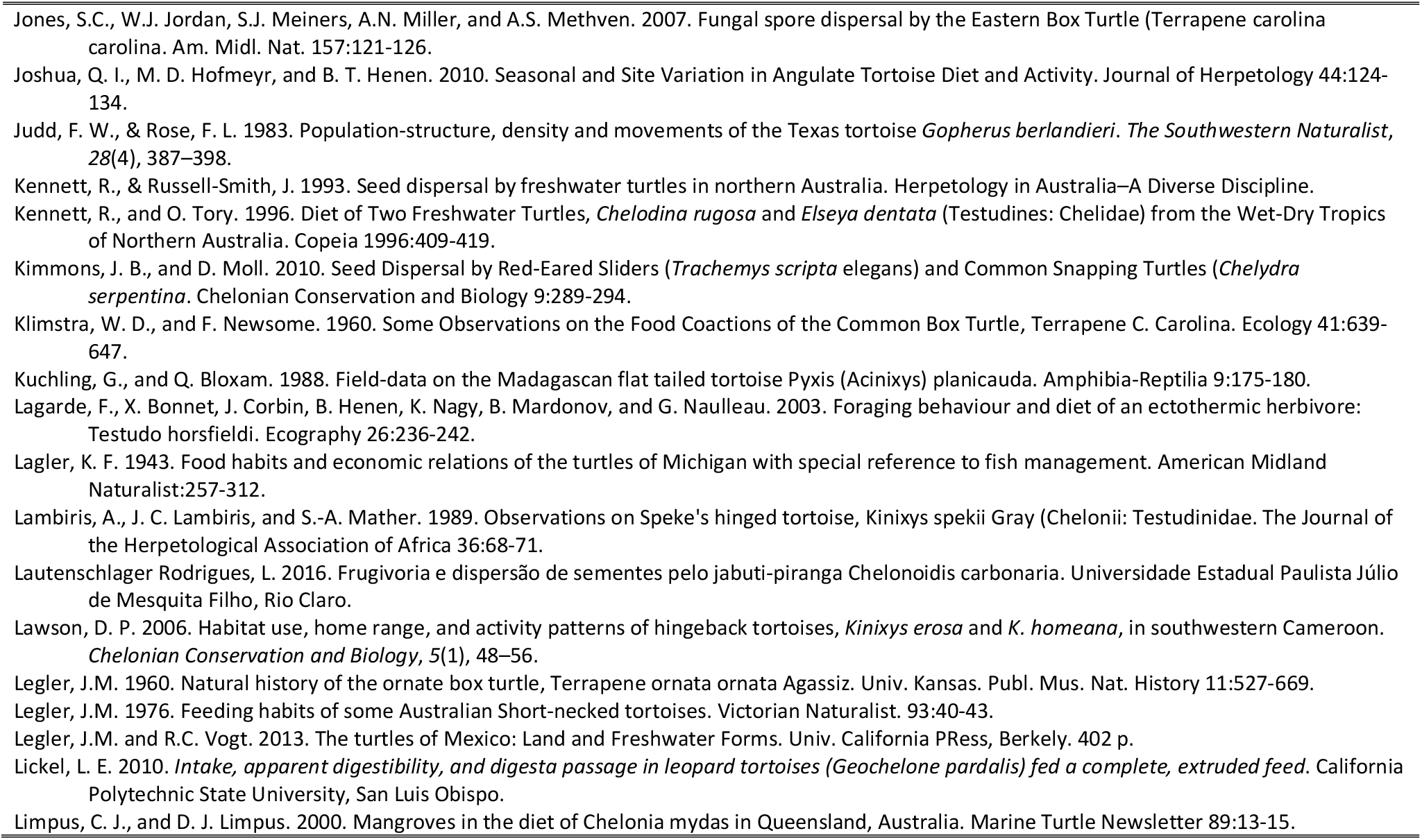
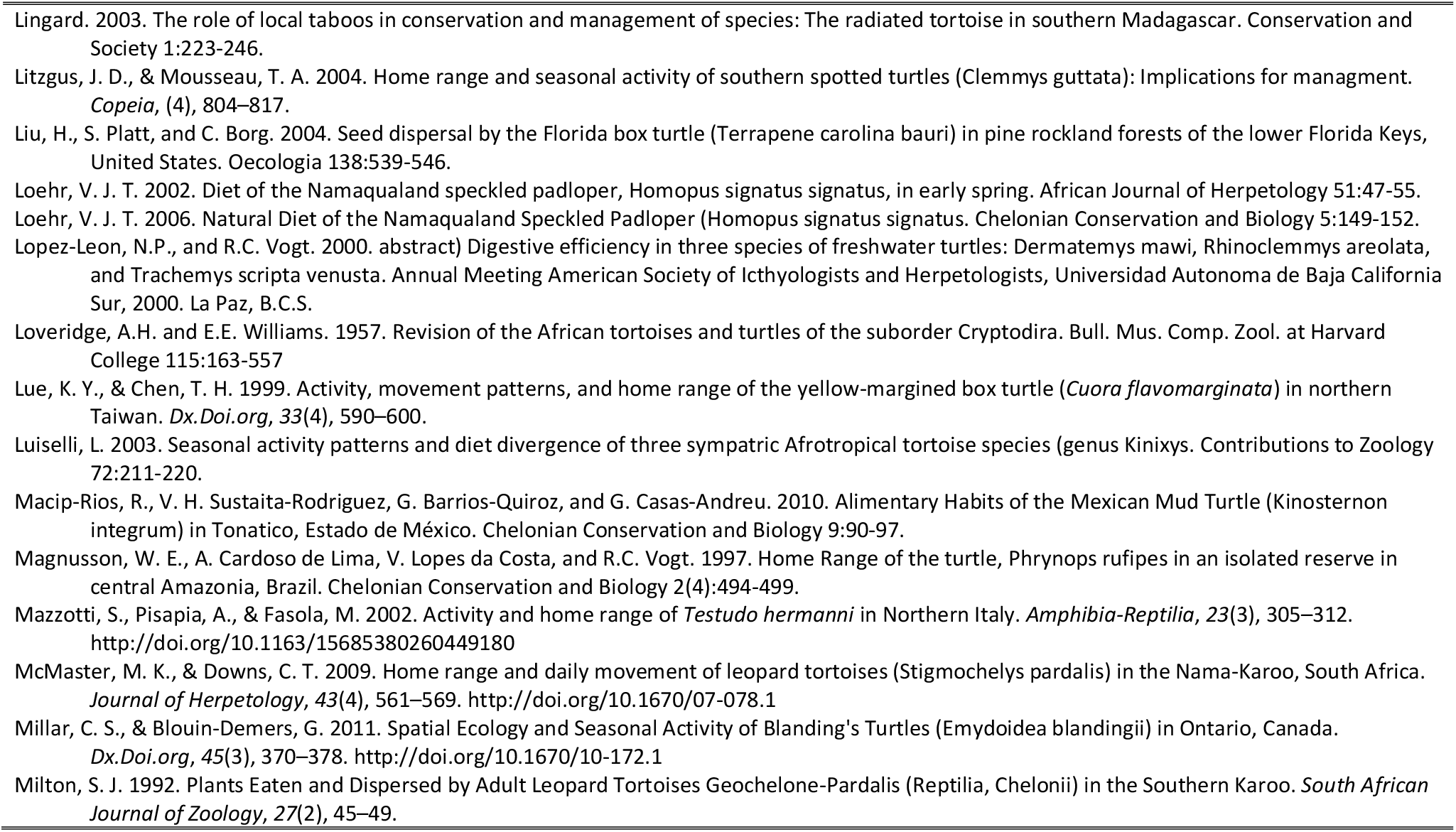
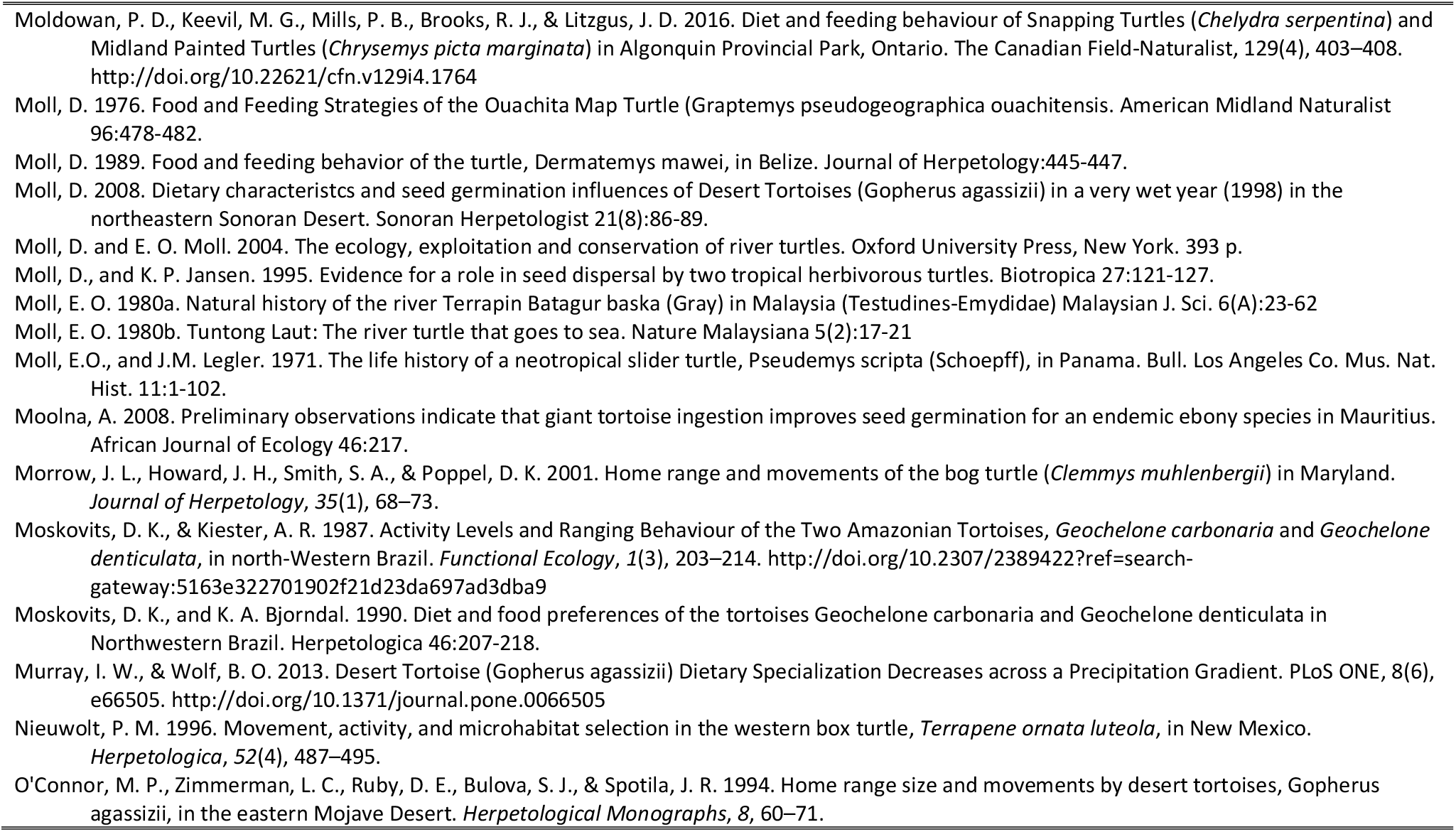
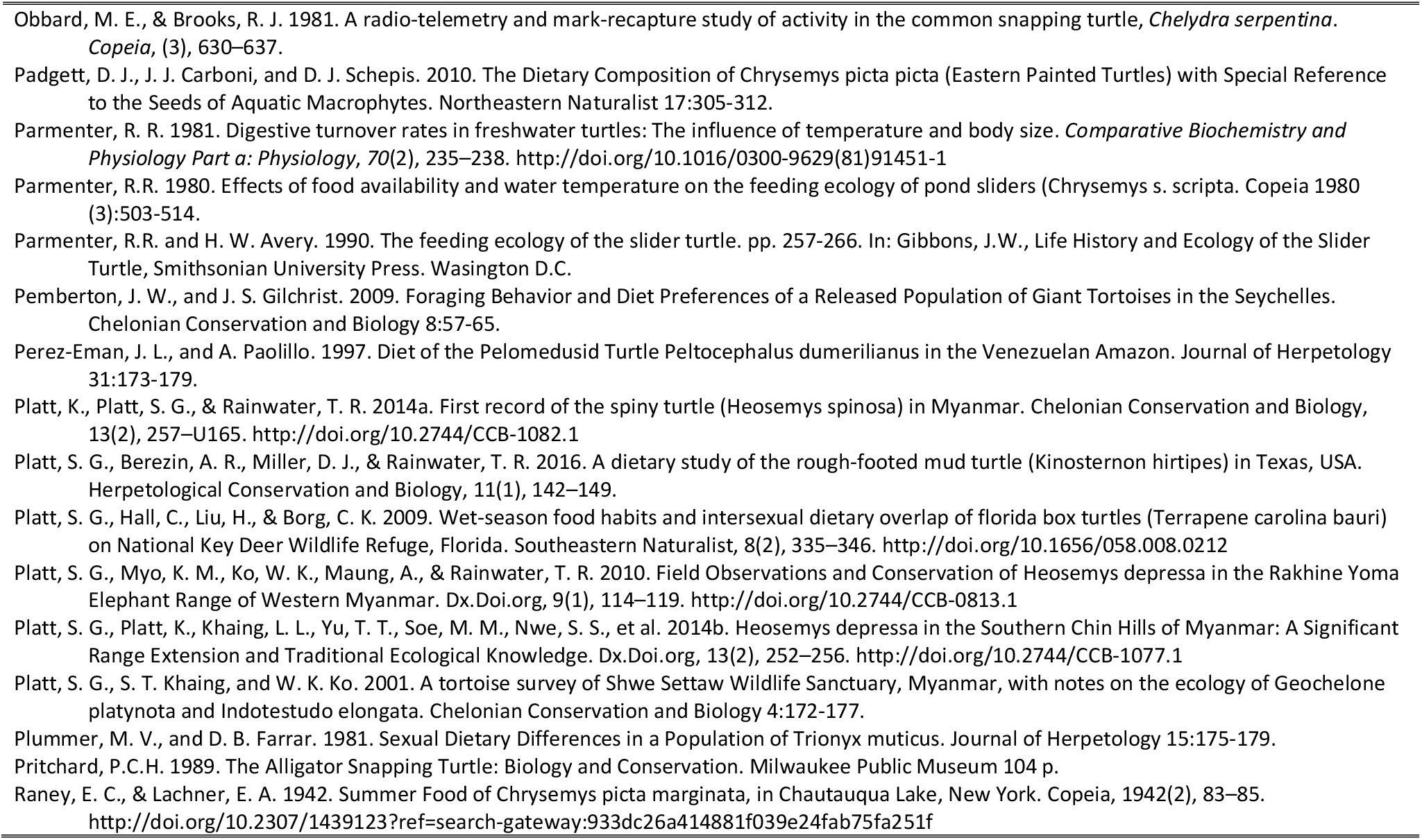
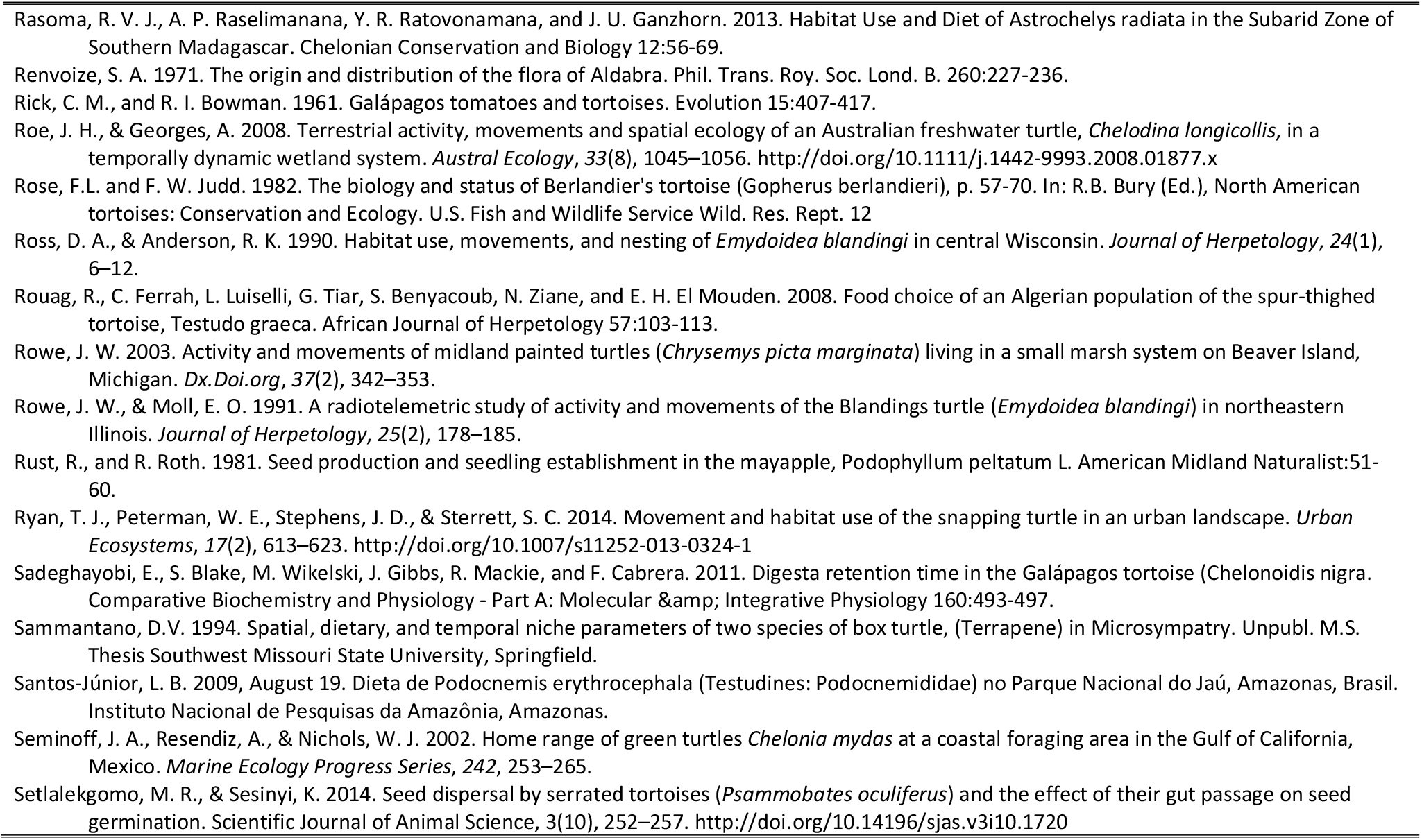
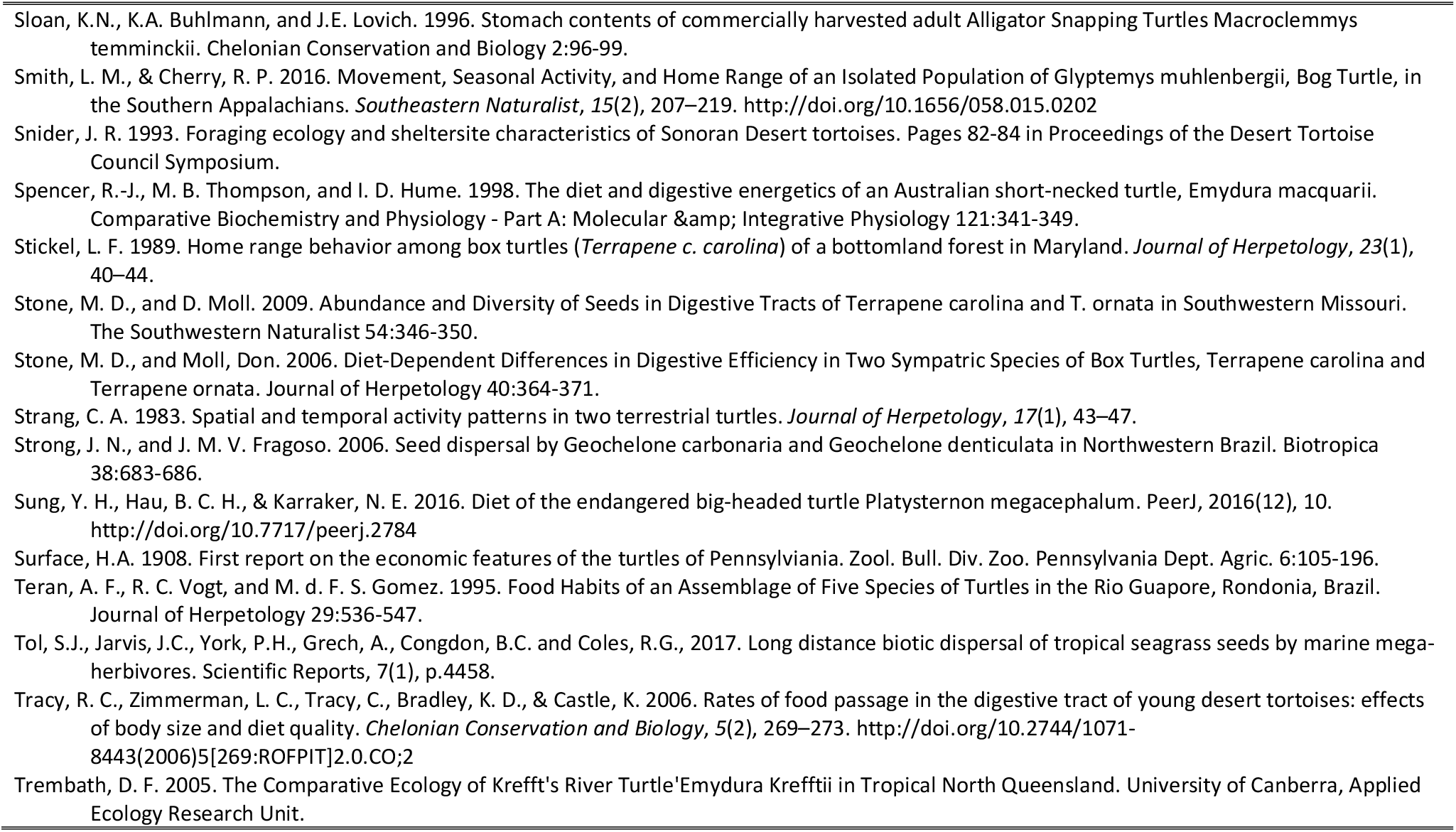
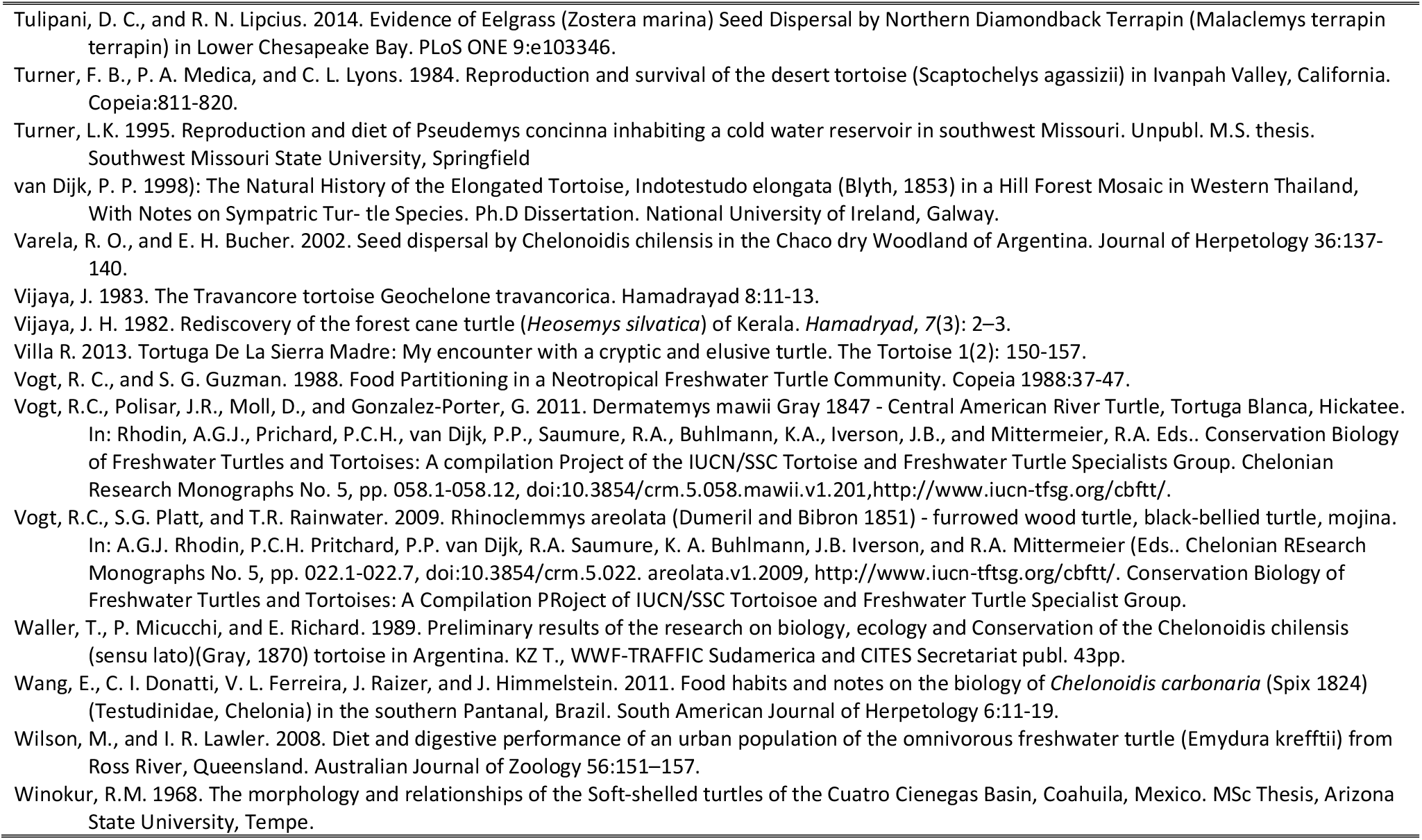
Full references for the literature and sources of information related to chelonian frugivory and seed dispersal, gut retention times, home range and movement, and germination success reviewed in this article.

**S2:** Chelonian species that engage in frugivory and seed dispersal, and the species of plants that they consumed and/or disperse.

| Reference | Chelonian species | Plant species |
| --- | --- | --- |
| Amorocho and Reina 2008 | *Chelonia mydas* | *Rhizophora mangle* |
| Andriantsaralaza et. al. 2013 | *Aldabrachelys gigantea* | *Adansonia fony* |
| Armstrong and Booth 2005 | *Elseya albagula* | *Costanospermum australe* |
| Arthur et. al. 2008 | *Chelonia mydas* | Unidentified 'mangrove' |
| Auffenberg and Weaver 1969 | *Gopherus berlandieri* | *Aristida* sp. |
| Auffenberg and Weaver 1969 | *Gopherus berlandieri* | *Aster* sp. |
| Auffenberg and Weaver 1969 | *Gopherus berlandieri* | *Buchloe* sp. |
| Auffenberg and Weaver 1969 | *Gopherus berlandieri* | *Celtis pallida* |
| Auffenberg and Weaver 1969 | *Gopherus berlandieri* | *Cenchrus* sp. |
| Auffenberg and Weaver 1969 | *Gopherus berlandieri* | *Chloris* sp. |
| Auffenberg and Weaver 1969 | *Gopherus berlandieri* | *Citharexylum* sp. |
| Auffenberg and Weaver 1969 | *Gopherus berlandieri* | *Plantago* sp. |
| Auffenberg and Weaver 1969 | *Gopherus berlandieri* | *Viola* sp. |
| Auffenberg and Weaver 1969 | *Gopherus berlandieri* | *Xanthophyllum* sp. |
| Auffenberg and Weaver 1969 | *Gopherus berlandieri* | *Cylindropuntia leptocaulis* |
| Auffenberg and Weaver 1969 | *Gopherus berlandieri* | *Opuntia engelmannii* |
| Ayres et. al. 2010 | *Emys orbicularis* | *Nymphaea alba* |
| Balensiefer and Vogt 2006 | *Podocnemis unifilis* | *Eichhornia* sp. |
| Balensiefer and Vogt 2006 | *Podocnemis unifilis* | *Pistia* sp. |
| Balensiefer and Vogt 2006 | *Podocnemis unifilis* | *Pseudobombax munguba* |
| Balensiefer and Vogt 2006 | *Podocnemis unifilis* | *Salvinia* sp. |
| Balensiefer and Vogt 2006 | *Podocnemis unifilis* | Unidentified Sapindaceae |
| Balensiefer and Vogt 2006 | *Podocnemis unifilis* | Unidentified Poaceae |
| Balensiefer and Vogt 2006 | *Podocnemis unifilis* | Unidentified Myrtaceae |
| Balensiefer and Vogt 2006 | *Podocnemis unifilis* | Unidentified Melastomataceae |
| Balensiefer and Vogt 2006 | *Podocnemis unifilis* | Unidentified Fabaceae |
| Balensiefer and Vogt 2006 | *Podocnemis unifilis* | Unidentified Fabaceae |
| Balensiefer and Vogt 2006 | *Podocnemis unifilis* | Unidentified Bombacaceae |
| Birkhead et. al. 2005 | *Gopherus polyphemus* | *Acalypha gracilens* |
| Birkhead et. al. 2005 | *Gopherus polyphemus* | *Ambrosia* sp. |
| Birkhead et. al. 2005 | *Gopherus polyphemus* | *Asclepias* sp. |
| Birkhead et. al. 2005 | *Gopherus polyphemus* | *Commelina erecta* |
| Birkhead et. al. 2005 | *Gopherus polyphemus* | *Crataegus* sp. |
| Birkhead et. al. 2005 | *Gopherus polyphemus* | *Digitaria* sp. |
| Birkhead et. al. 2005 | *Gopherus polyphemus* | *Dyschoriste oblongifolia* |
| Birkhead et. al. 2005 | *Gopherus polyphemus* | *Gaillardia aestivalis* |
| Birkhead et. al. 2005 | *Gopherus polyphemus* | *Hypericum* sp. |
| Birkhead et. al. 2005 | *Gopherus polyphemus* | *Licania michauxii* |
| Birkhead et. al. 2005 | *Gopherus polyphemus* | *Mollugo verticillata* |
| Birkhead et. al. 2005 | *Gopherus polyphemus* | *Oenothera* sp. |
| Birkhead et. al. 2005 | *Gopherus polyphemus* | *Opuntia humifusa* |
| Birkhead et. al. 2005 | *Gopherus polyphemus* | *Oxalis* sp. |
| Birkhead et. al. 2005 | *Gopherus polyphemus* | *Panicum* sp. |
| Birkhead et. al. 2005 | *Gopherus polyphemus* | *Paspalum* sp. |
| Birkhead et. al. 2005 | *Gopherus polyphemus* | *Physalis heterophylla* |
| Birkhead et. al. 2005 | *Gopherus polyphemus* | *Plantago* sp. |
| Birkhead et. al. 2005 | *Gopherus polyphemus* | *Polygala* sp. |
| Birkhead et. al. 2005 | *Gopherus polyphemus* | *Polygonum* sp. |
| Birkhead et. al. 2005 | *Gopherus polyphemus* | *Prunus angustifolia* |
| Birkhead et. al. 2005 | *Gopherus polyphemus* | *Prunus* sp. |
| Birkhead et. al. 2005 | *Gopherus polyphemus* | *Rhynchospora* sp. |
| Birkhead et. al. 2005 | *Gopherus polyphemus* | *Rubus* sp. |
| Birkhead et. al. 2005 | *Gopherus polyphemus* | *Rumex* sp. |
| Birkhead et. al. 2005 | *Gopherus polyphemus* | *Sclerodactylon macrostachyum* |
| Birkhead et. al. 2005 | *Gopherus polyphemus* | *Stellaria media* |
| Birkhead et. al. 2005 | *Gopherus polyphemus* | *Tradescantia ohiensis* |
| Birkhead et. al. 2005 | *Gopherus polyphemus* | *Tragia urens* |
| Birkhead et. al. 2005 | *Gopherus polyphemus* | *Veronica hederifolia* |
| Birkhead et. al. 2005 | *Gopherus polyphemus* | *Zornia bracteata* |
| Birkhead et. al. 2005 | *Gopherus polyphemus* | *Ipomoea* sp. |
| Birkhead et. al. 2005 | *Gopherus polyphemus* | *Stylosanthes biflora* |
| Birkhead et. al. 2005 | *Gopherus polyphemus* | *Asimina angustifolia* |
| Birkhead et. al. 2005 | *Gopherus polyphemus* | *Passiflora edulis* |
| Birkhead et. al. 2005 | *Gopherus polyphemus* | *Piriqueta cistoides* |
| Birkhead et. al. 2005 | *Gopherus polyphemus* | *Mimosa quadrivalvis* |
| Birkhead et. al. 2005 | *Gopherus polyphemus* | *Diodia* sp. |
| Birkhead et. al. 2005 | *Gopherus polyphemus* | *Richardia* sp. |
| Blake et. al. 2012 | *Chelonoidis nigra* | *Antheophora hermaphrodita* |
| Blake et. al. 2012 | *Chelonoidis nigra* | *Blainvillea dichotoma* |
| Blake et. al. 2012 | *Chelonoidis nigra* | *Brickellia diffusa* |
| Blake et. al. 2012 | *Chelonoidis nigra* | *Cenchrus platyacanthus* |
| Blake et. al. 2012 | *Chelonoidis nigra* | *Commelina diffusa* |
| Blake et. al. 2012 | *Chelonoidis nigra* | *Cordia lutea* |
| Blake et. al. 2012 | *Chelonoidis nigra* | *Crotalaria pumila* |
| Blake et. al. 2012 | *Chelonoidis nigra* | *Cynodon dactylon* |
| Blake et. al. 2012 | *Chelonoidis nigra* | *Cyperus ligularis* |
| Blake et. al. 2012 | *Chelonoidis nigra* | *Desmodium incanum* |
| Blake et. al. 2012 | *Chelonoidis nigra* | *Digitaria setigera* |
| Blake et. al. 2012 | *Chelonoidis nigra* | *Eleocharis maculosa* |
| Blake et. al. 2012 | *Chelonoidis nigra* | *Eleusine indica* |
| Blake et. al. 2012 | *Chelonoidis nigra* | *Eriochloa pacifica* |
| Blake et. al. 2012 | *Chelonoidis nigra* | *Hippomane mancinella* |
| Blake et. al. 2012 | *Chelonoidis nigra* | *Ipomoea triloba* |
| Blake et. al. 2012 | *Chelonoidis nigra* | *Kyllinga brevifolia* |
| Blake et. al. 2012 | *Chelonoidis nigra* | *Opuntia echios* |
| Blake et. al. 2012 | *Chelonoidis nigra* | *Panicum dichotomiflorum* |
| Blake et. al. 2012 | *Chelonoidis nigra* | *Panicum maximum* |
| Blake et. al. 2012 | *Chelonoidis nigra* | *Paspalum conjugatum* |
| Blake et. al. 2012 | *Chelonoidis nigra* | *Passiflora edulis* |
| Blake et. al. 2012 | *Chelonoidis nigra* | *Physalis pubescens* |
| Blake et. al. 2012 | *Chelonoidis nigra* | *Pisonia floribunda* |
| Blake et. al. 2012 | *Chelonoidis nigra* | *Polygonum opelousanum* |
| Blake et. al. 2012 | *Chelonoidis nigra* | *Portulaca oleracea* |
| Blake et. al. 2012 | *Chelonoidis nigra* | *Psidium galapageium* |
| Blake et. al. 2012 | *Chelonoidis nigra* | *Psidium guajava* |
| Blake et. al. 2012 | *Chelonoidis nigra* | *Rubus niveus* |
| Blake et. al. 2012 | *Chelonoidis nigra* | *Scleria distans* |
| Blake et. al. 2012 | *Chelonoidis nigra* | *Scleria hirtella* |
| Blake et. al. 2012 | *Chelonoidis nigra* | *Sida rhombifolia* |
| Blake et. al. 2012 | *Chelonoidis nigra* | *Sida spinosa* |
| Blake et. al. 2012 | *Chelonoidis nigra* | *Sida spinosa* |
| Blake et. al. 2012 | *Chelonoidis nigra* | *Solanum americanum* |
| Blake et. al. 2012 | *Chelonoidis nigra* | *Stachytarpheta cayennensis* |
| Blake et. al. 2012 | *Chelonoidis nigra* | *Synedrella nodiflora* |
| Blake et. al. 2012 | *Chelonoidis nigra* | *Tradescantia fluminensis* |
| Blake et. al. 2012 | *Chelonoidis nigra* | *Zanthoxylum fagara* |
| Blake et. al. 2012 | *Chelonoidis nigra* | *Acacia rorudia* |
| Blake et. al. 2012 | *Chelonoidis nigra* | *Bidens* sp. |
| Blake et. al. 2012 | *Chelonoidis nigra* | *Galactia striata* |
| Blake et. al. 2012 | *Chelonoidis nigra* | *Clerodendrum villosum* |
| Blake et. al. 2012 | *Chelonoidis nigra* | *Brachiaria multiculma* |
| Blake et. al. 2012 | *Chelonoidis nigra* | *Brachiaria mutica* |
| Blake et. al. 2012 | *Chelonoidis nigra* | *Hippomane mancinella* |
| Blake et. al. 2012 | *Chelonoidis nigra* | *Opuntia echios* |
| Blake et. al. 2012 | *Chelonoidis nigra* | *Passiflora edulis* |
| Blake et. al. 2012 | *Chelonoidis nigra* | *Psidium galapageium* |
| Blake et. al. 2012 | *Chelonoidis nigra* | *Psidium guajava* |
| Blake et. al. 2015 | *Chelonoidis nigra* | *Antheophora hermaphrodita* |
| Blake et. al. 2015 | *Chelonoidis nigra* | *Axonopus micay* |
| Blake et. al. 2015 | *Chelonoidis nigra* | *Blainvillea dichotoma* |
| Blake et. al. 2015 | *Chelonoidis nigra* | *Brickellia diffusa* |
| Blake et. al. 2015 | *Chelonoidis nigra* | *Cenchrus platyacanthus* |
| Blake et. al. 2015 | *Chelonoidis nigra* | *Cordia lutea* |
| Blake et. al. 2015 | *Chelonoidis nigra* | *Cyperus ligularis* |
| Blake et. al. 2015 | *Chelonoidis nigra* | *Desmodium glabrum* |
| Blake et. al. 2015 | *Chelonoidis nigra* | *Desmodium incanum* |
| Blake et. al. 2015 | *Chelonoidis nigra* | *Digitaria setigera* |
| Blake et. al. 2015 | *Chelonoidis nigra* | *Eleocharis maculosa* |
| Blake et. al. 2015 | *Chelonoidis nigra* | *Eleusine indica* |
| Blake et. al. 2015 | *Chelonoidis nigra* | *Eragrostis cilianensis* |
| Blake et. al. 2015 | *Chelonoidis nigra* | *Eriochloa pacifica* |
| Blake et. al. 2015 | *Chelonoidis nigra* | *Hippomane mancinella* |
| Blake et. al. 2015 | *Chelonoidis nigra* | *Ipomoea triloba* |
| Blake et. al. 2015 | *Chelonoidis nigra* | *Opuntia echios* |
| Blake et. al. 2015 | *Chelonoidis nigra* | *Panicum dichotomiflorum* |
| Blake et. al. 2015 | *Chelonoidis nigra* | *Panicum maximum* |
| Blake et. al. 2015 | *Chelonoidis nigra* | *Paspalum conjugatum* |
| Blake et. al. 2015 | *Chelonoidis nigra* | *Passiflora edulis* |
| Blake et. al. 2015 | *Chelonoidis nigra* | *Pennisetum purpureum* |
| Blake et. al. 2015 | *Chelonoidis nigra* | *Physalis pubescens* |
| Blake et. al. 2015 | *Chelonoidis nigra* | *Pisonia floribunda* |
| Blake et. al. 2015 | *Chelonoidis nigra* | *Polygonum opelousanum* |
| Blake et. al. 2015 | *Chelonoidis nigra* | *Portulaca oleracea* |
| Blake et. al. 2015 | *Chelonoidis nigra* | *Psidium galapageium* |
| Blake et. al. 2015 | *Chelonoidis nigra* | *Psidium guajava* |
| Blake et. al. 2015 | *Chelonoidis nigra* | *Rubus niveus* |
| Blake et. al. 2015 | *Chelonoidis nigra* | *Scleria distans* |
| Blake et. al. 2015 | *Chelonoidis nigra* | *Scleria hirtella* |
| Blake et. al. 2015 | *Chelonoidis nigra* | *Sida rhombifolia* |
| Blake et. al. 2015 | *Chelonoidis nigra* | *Sida salviifolia* |
| Blake et. al. 2015 | *Chelonoidis nigra* | *Silene dichotoma* |
| Blake et. al. 2015 | *Chelonoidis nigra* | *Solanum ochraceo-ferrugineum* |
| Blake et. al. 2015 | *Chelonoidis nigra* | *Stachytarpheta cayennensis* |
| Blake et. al. 2015 | *Chelonoidis nigra* | *Synedrella nodiflora* |
| Blake et. al. 2015 | *Chelonoidis nigra* | *Tradescantia fluminensis* |
| Blake et. al. 2015 | *Chelonoidis nigra* | *Zanthoxylum fagara* |
| Blake et. al. 2015 | *Chelonoidis nigra* | *Acacia rorudia* |
| Blake et. al. 2015 | *Chelonoidis nigra* | *Bidens* sp. |
| Blake et. al. 2015 | *Chelonoidis nigra* | *Clerodendrum villosum* |
| Bonin et. al. 2006 | *Batagur baska* | *Sorocea* sp. |
| Bonin et. al. 2006 | *Batagur borneonensis* | Unidentified 'mangrove' |
| Bonin et. al. 2006 | *Carettochelys insculpta* | *Pandanus aquaticus* |
| Bonin et. al. 2006 | *Carettochelys insculpta* | *Syzygium forte* |
| Bonin et. al. 2006 | *Chelonoidis denticulata* | *Jacaratia spinosa* |
| Bonin et. al. 2006 | *Kinosternon baurii* | Unidentified Arecaceae |
| Bonin et. al. 2006 | *Kinosternon scorpioides* | Unidentified Arecaceae |
| Bonin et. al. 2006 | *Macrochelys temminckii* | *Quercus* sp. |
| Bonin et. al. 2006 | *Mesoclemmys nasuta* | *Philodendron* sp. |
| Bonin et. al. 2006 | *Trionyx triunguis* | *Phoenix* sp. |
| Bonin et. al. 2006 | *Emydura subglobosa* | *Pandanus* sp. |
| Braun and Brooks 1987 | *Terrapene carolina* | *Podophyllum peltatum* |
| Braun and Brooks 1987 | *Terrapene carolina* | *Podophyllum peltatum* |
| Braun and Brooks 1987 | *Terrapene carolina* | *Rubus* sp. |
| Braun and Brooks 1987 | *Terrapene carolina* | *Vaccinium* sp. |
| Braun and Brooks 1987 | *Terrapene carolina* | *Viburnum* sp. |
| Braun and Brooks 1987 | *Terrapene carolina* | *Vitis rotundifolia* |
| Braun and Brooks 1987 | *Terrapene carolina* | *Arisaema triphyllum* |
| Braun and Brooks 1987 | *Terrapene carolina* | *Duchesnea indica* |
| Braun and Brooks 1987 | *Terrapene carolina* | *Fragaria virginiana* |
| Braun and Brooks 1987 | *Terrapene carolina* | *Gaylussacia baccata* |
| Braun and Brooks 1987 | *Terrapene carolina* | *Morus alba* |
| Braun and Brooks 1987 | *Terrapene carolina* | *Phytolacca americana* |
| Braun and Brooks 1987 | *Terrapene carolina* | *Podophyllum peltatum* |
| Braun and Brooks 1987 | *Terrapene carolina* | *Podophyllum peltatum* |
| Braun and Brooks 1987 | *Terrapene carolina* | *Prunus* sp. |
| Braun and Brooks 1987 | *Terrapene carolina* | *Rosa multiflora* |
| Braun and Brooks 1987 | *Terrapene carolina* | *Rubus phoenicolasius* |
| Braun and Brooks 1987 | *Terrapene carolina* | *Rubus* sp. |
| Braun and Brooks 1987 | *Terrapene carolina* | *Vaccinium vacillans* |
| Braun and Brooks 1987 | *Terrapene carolina* | *Vitis aestivalis* |
| Braun and Brooks 1987 | *Terrapene carolina* | *Vitis rotundifolia* |
| Braun and Brooks 1987 | *Terrapene carolina* | *Vitis vulpina* |
| Braun and Brooks 1987 | *Terrapene carolina* | *Passiflora edulis* |
| Burgin and Renshaw 2008 | *Chelodina longicollis* | *Eleocharis acuta* |
| Burgin and Renshaw 2008 | *Chelodina longicollis* | *Eleocharis* sp. |
| Burgin and Renshaw 2008 | *Chelodina longicollis* | *Gahnia* sp. |
| Burgin and Renshaw 2008 | *Chelodina longicollis* | *Juncus* sp. |
| Burgin and Renshaw 2008 | *Chelodina longicollis* | *Paspalum dilatatum* |
| Burgin and Renshaw 2008 | *Chelodina longicollis* | *Polygonum* sp. |
| Burgin and Renshaw 2008 | *Chelodina longicollis* | *Potamogeton* sp. |
| Burgin and Renshaw 2008 | *Chelodina longicollis* | *Sagittaria graminea* |
| Burgin and Renshaw 2008 | *Chelodina longicollis* | *Scirpus* sp. |
| Calviño-Cancela et. al. 2007 | *Emys orbicularis* | *Nymphaea alba* |
| Caputo and Vogt 2008 | *Phrynops rufipes* | *Bactris* sp. |
| Caputo and Vogt 2008 | *Phrynops rufipes* | *Iriartella setigera* |
| Caputo and Vogt 2008 | *Phrynops rufipes* | *Mauritia flexuosa* |
| Caputo and Vogt 2008 | *Phrynops rufipes* | *Mauritia flexuosa* |
| Caputo and Vogt 2008 | *Phrynops rufipes* | *Oenocarpus bataua* |
| Caputo and Vogt 2008 | *Phrynops rufipes* | *Socratea exorrhiza* |
| Caputo and Vogt 2008 | *Phrynops rufipes* | *Socratea exorrhiza* |
| Caputo and Vogt 2008 | *Phrynops rufipes* | Unidentified Sapotaceae |
| Caputo and Vogt 2008 | *Phrynops rufipes* | Unidentified Rubiaceae |
| Caputo and Vogt 2008 | *Phrynops rufipes* | Unidentified Malvaceae |
| Caputo and Vogt 2008 | *Phrynops rufipes* | Unidentified Fabaceae |
| Caputo and Vogt 2008 | *Phrynops rufipes* | Unidentified Clusiaceae |
| Caputo and Vogt 2008 | *Phrynops rufipes* | Unidentified Arecaceae |
| Caputo and Vogt 2008 | *Phrynops rufipes* | *Euterpe precatoria* |
| Carlson et. al. 2003 | *Gopherus polyphemus* | *Digitaria* sp. |
| Carlson et. al. 2003 | *Gopherus polyphemus* | *Diodella teres* |
| Carlson et. al. 2003 | *Gopherus polyphemus* | *Euphorbia maculata* |
| Carlson et. al. 2003 | *Gopherus polyphemus* | *Licania michauxii* |
| Carlson et. al. 2003 | *Gopherus polyphemus* | *Paspalum notatum* |
| Carlson et. al. 2003 | *Gopherus polyphemus* | *Paspalum setaceum* |
| Carlson et. al. 2003 | *Gopherus polyphemus* | *Quercus geminata* |
| Carlson et. al. 2003 | *Gopherus polyphemus* | *Paspalum notatum* |
| Carlson et. al. 2003 | *Gopherus polyphemus* | *Paspalum setaceum* |
| Cobo and Reu 1988 | *Testudo graeca* | *Agrostis* sp. |
| Cobo and Reu 1988 | *Testudo graeca* | *Anthoxanthum ovatum* |
| Cobo and Reu 1988 | *Testudo graeca* | *Briza maxima* |
| Cobo and Reu 1988 | *Testudo graeca* | *Briza minor* |
| Cobo and Reu 1988 | *Testudo graeca* | *Carduus meonanthus* |
| Cobo and Reu 1988 | *Testudo graeca* | *Carduus* sp. |
| Cobo and Reu 1988 | *Testudo graeca* | *Cerastium glomeratum* |
| Cobo and Reu 1988 | *Testudo graeca* | *Corynephorus* sp. |
| Cobo and Reu 1988 | *Testudo graeca* | *Cynodon dactylon* |
| Cobo and Reu 1988 | *Testudo graeca* | *Erodium* sp. |
| Cobo and Reu 1988 | *Testudo graeca* | *Halimium halimifolium* |
| Cobo and Reu 1988 | *Testudo graeca* | *Hypochaeris glabra* |
| Cobo and Reu 1988 | *Testudo graeca* | *Isolepis* sp. |
| Cobo and Reu 1988 | *Testudo graeca* | *Juncus* sp. |
| Cobo and Reu 1988 | *Testudo graeca* | *Leontodon taraxacoides* |
| Cobo and Reu 1988 | *Testudo graeca* | *Malcolmia lacera* |
| Cobo and Reu 1988 | *Testudo graeca* | *Moenchia erecta* |
| Cobo and Reu 1988 | *Testudo graeca* | *Ononis* sp. |
| Cobo and Reu 1988 | *Testudo graeca* | *Panicum repens* |
| Cobo and Reu 1988 | *Testudo graeca* | *Paspalum* sp. |
| Cobo and Reu 1988 | *Testudo graeca* | *Polypogon maritimus* |
| Cobo and Reu 1988 | *Testudo graeca* | *Ranunculus sardous* |
| Cobo and Reu 1988 | *Testudo graeca* | *Ranunculus* sp. |
| Cobo and Reu 1988 | *Testudo graeca* | *Reseda media* |
| Cobo and Reu 1988 | *Testudo graeca* | *Rubus ulmifolius* |
| Cobo and Reu 1988 | *Testudo graeca* | *Rumex bucephalophorus* |
| Cobo and Reu 1988 | *Testudo graeca* | *Simaba* sp. |
| Cobo and Reu 1988 | *Testudo graeca* | *Spergula arvensis* |
| Cobo and Reu 1988 | *Testudo graeca* | *Sporobolus* sp. |
| Cobo and Reu 1988 | *Testudo graeca* | *Vulpia* sp. |
| Cobo and Reu 1988 | *Testudo graeca* | *Lotus subbiflorus* |
| Cobo and Reu 1988 | *Testudo graeca* | *Ornithophus sativus* |
| Cobo and Reu 1988 | *Testudo graeca* | *Astragalus pelecinus* |
| Cobo and Reu 1988 | *Testudo graeca* | *Bromus rigidus* |
| Cobo and Reu 1988 | *Testudo graeca* | *Briza maxima* |
| Cobo and Reu 1988 | *testudo graeca* | *Hypochaeris glabra* |
| Cobo and Reu 1988 | *Testudo graeca* | *Rumex bucephalophorus* |
| Cobo and Reu 1988 | *Testudo graeca* | *Spergula arvensis* |
| Cobo and Reu 1988 | *Testudo graeca* | *Anagallis arvensis* |
| da Costa 2012 | *Podocnemis expansa* | *Pouteria elegans* |
| da Costa 2012 | *Podocnemis expansa* | *Psidium* sp. |
| da Costa 2012 | *Podocnemis expansa* | *Oryza* sp. |
| da Costa 2012 | *Podocnemis expansa* | *Duroia* sp. |
| da Costa 2012 | *Podocnemis expansa* | Unidentified |
| da Costa 2012 | *Podocnemis expansa* | Unidentified |
| da Costa 2012 | *Podocnemis expansa* | Unidentified |
| Deepak 2011 | *Indotestudo travancorica* | *Artocarpus* sp. |
| Deepak 2011 | *Indotestudo travancorica* | *Dillenia pentagyna* |
| Deepak 2011 | *Indotestudo travancorica* | *Dillenia pentagyna* |
| Deepak 2011 | *Indotestudo travancorica* | *Ficus virens* |
| Deepak 2011 | *Indotestudo travancorica* | *Gomphandra* sp. |
| Deepak 2011 | *Indotestudo travancorica* | *Grewia tiliifolia* |
| Deepak 2011 | *Indotestudo travancorica* | *Lantana camara* |
| de Lima et. al. 1997 | *Phrynops rufipes* | *Euterpe precatoria* |
| de Lima et. al. 1997 | *Phrynops rufipes* | *Iriartella* sp. |
| de Lima et. al. 1997 | *Phrynops rufipes* | *Oenocarpus bacaba* |
| de Lima et. al. 1997 | *Phrynops rufipes* | "Munbaca" sp. |
| de Lima et. al. 1997 | *Phrynops rufipes* | "Pupunharana" sp. |
| de Lima et. al. 1997 | *Phrynops rufipes* | *Socratea exorrhiza* |
| de Lima et. al. 1997 | *Phrynops rufipes* | *Oenocarpus bataua* |
| de Lima et. al. 1997 | *Phrynops rufipes* | Unidentified Humiriaceae |
| de Lima et. al. 1997 | *Phrynops rufipes* | Unidentified Fabaceae |
| de Lima et. al. 1997 | *Phrynops rufipes* | Unidentified Euphorbiaceae |
| de Lima et. al. 1997 | *Phrynops rufipes* | Unidentified [Annonaceae] |
| del Vecchio et. al. 2011 | *Testudo hermanni* | *Carduus pycnocephalus* |
| del Vecchio et. al. 2011 | *Testudo hermanni* | *Hedera helix* |
| del Vecchio et. al. 2011 | *Testudo hermanni* | *Rubus ulmifolius* |
| del Vecchio et. al. 2011 | *Testudo hermanni* | *Ruscus aculeatus* |
| del Vecchio et. al. 2011 | *Testudo hermanni* | Unidentified Fabaceae |
| de Neira and Johnson 1985 | *Chelonoidis nigra* | *Psidium galapageium* |
| Elbers and Moll 2011 | *Macrochelys temminckii* | *Diospyros virginiana* |
| Elbers and Moll 2011 | *Macrochelys temminckii* | *Nyssa aquatica* |
| Elbers and Moll 2011 | *Macrochelys temminckii* | *Quercus phellos* |
| Ellis-Soto et. al. 2017 | *Chelonoidis donfaustoi* | *Passiflora edulis* |
| Ellis-Soto et. al. 2017 | *Chelonoidis donfaustoi* | *Psidium guajava* |
| Ellis-Soto et. al. 2017 | *Chelonoidis porteri* | *Passiflora edulis* |
| Ellis-Soto et. al. 2017 | *Chelonoidis porteri* | *Psidium guajava* |
| Elsey 2006 | *Macrochelys temminckii* | *Quercus sp.* |
| Evenden 1948 | *Actinemys marmorata* | *Nuphar polysepala* |
| Fachín-Terán 1995 et. al. | *Mesoclemmys raniceps* | Unidentified Fabaceae |
| Fachín-Terán 1995 et. al. | *Mesoclemmys raniceps* | Unidentified Myrtaceae |
| Fachín-Terán 1995 et. al. | *Mesoclemmys raniceps* | Unidentified Sapotaceae |
| Fachín-Terán 1995 et. al. | *Podocnemis unifilis* | *Diospyros sp.* |
| Fachín-Terán 1995 et. al. | *Podocnemis unifilis* | *Margaritaria sp.* |
| Fachín-Terán 1995 et. al. | *Podocnemis unifilis* | *Maripa sp.* |
| Fachín-Terán 1995 et. al. | *Podocnemis unifilis* | *Pouteria sp.* |
| Fachín-Terán 1995 et. al. | *Podocnemis unifilis* | Unidentified Fabaceae |
| Falcón 2018 | *Aldabrachelys gigantea* | *Allophylus aldabricus* |
| Falcón 2018 | *Aldabrachelys gigantea* | *Apodytes dimidiata* |
| Falcón 2018 | *Aldabrachelys gigantea* | *Azima tetraacantha* |
| Falcón 2018 | *Aldabrachelys gigantea* | *Capparis cartilaginea* |
| Falcón 2018 | *Aldabrachelys gigantea* | *Volkameria glabra* |
| Falcón 2018 | *Aldabrachelys gigantea* | *Cordia subcordata* |
| Falcón 2018 | *Aldabrachelys gigantea* | *Ehretia cymosa* |
| Falcón 2018 | *Aldabrachelys gigantea* | *Ficus sundaica* |
| Falcón 2018 | *Aldabrachelys gigantea* | *Ficus lutea* |
| Falcón 2018 | *Aldabrachelys gigantea* | *Ficus reflexa* |
| Falcón 2018 | *Aldabrachelys gigantea* | *Flacourtia ratmonchii* |
| Falcón 2018 | *Aldabrachelys gigantea* | *Guettarda speciosa* |
| Falcón 2018 | *Aldabrachelys gigantea* | *Aloe aldabrensis* |
| Falcón 2018 | *Aldabrachelys gigantea* | *Cassine aethiopica* |
| Falcón 2018 | *Aldabrachelys gigantea* | *Ochna ciliata* |
| Falcón 2018 | *Aldabrachelys gigantea* | *Pandanus tectorius* |
| Falcón 2018 | *Aldabrachelys gigantea* | *Scaevola taccada* |
| Falcón 2018 | *Aldabrachelys gigantea* | *Solanum aldabrensis* |
| Falcón 2018 | *Aldabrachelys gigantea* | *Terminalia boivinii* |
| Falcón 2018 | *Aldabrachelys gigantea* | *Thespesia populnea* |
| Falcón 2018 | *Aldabrachelys gigantea* | *Thespesia populneides* |
| Falcón 2018 | *Aldabrachelys gigantea* | *Colubrina asiatica* |
| Falcón 2018 | *Aldabrachelys gigantea* | *Pemphis acidula* |
| Falcón 2018 | *Aldabrachelys gigantea* | *Casuarina equisetifolia* |
| Falcón 2018 | *Aldabrachelys gigantea* | *Tournefortia argentata* |
| Falcón 2018 | *Aldabrachelys gigantea* | *Abrus prectorius* |
| Falcón 2018 | *Aldabrachelys gigantea* | *Acalypha claoxyoides* |
| Figueroa et. al. 2012 | *Podocnemis expansa* | *Erythrina* sp. |
| Figueroa et. al. 2012 | *Podocnemis expansa* | *Ficus* sp. |
| Figueroa et. al. 2012 | *Podocnemis expansa* | *Macrolobium acaciifolium* |
| Ford and Moll 2004 | *Sternotherus odoratus* | *Ludwigia peploides* |
| Freeman 2010 | *Elseya lavarackorum* | *Ficus racemosa* |
| Freeman 2010 | *Elseya lavarackorum* | *Livistona rigida* |
| Freeman 2010 | *Elseya lavarackorum* | *Nauclea orientalis* |
| Freeman 2010 | *Elseya lavarackorum* | *Pandanus aquaticus* |
| Freeman 2010 | *Elseya lavarackorum* | *Passiflora foetida* |
| Georges and Kennet 1989 | *Carettochelys insculpta* | *Ficus racemosa* |
| Georges and Kennet 1989 | *Carettochelys insculpta* | *Pandanus aquaticus* |
| Georges and Kennet 1989 | *Carettochelys insculpta* | *Syzygium forte* |
| Georges et. al. 2008 | *Carettochelys insculpta* | *Artocarpus altilis* |
| Georges et. al. 2008 | *Carettochelys insculpta* | *Canarium indicum* |
| Georges et. al. 2008 | *Carettochelys insculpta* | *Ficus racemosa* |
| Georges et. al. 2008 | *Carettochelys insculpta* | *Nypa* sp. |
| Georges et. al. 2008 | *Carettochelys insculpta* | *Pandanus aquaticus* |
| Georges et. al. 2008 | *Carettochelys insculpta* | *Saccharum robustum* |
| Georges et. al. 2008 | *Carettochelys insculpta* | *Syzygium forte* |
| Georges et. al. 2008 | *Carettochelys insculpta* | *Xylocarpus* sp. |
| Gibbs et. al. 2008 | *Chelonoidis nigra* | *Opuntia megasperma* |
| Goulding 1980 | *Podocnemis expansa* | *Hevea spruceana* |
| Goulding 1980 | *Podocnemis expansa* | *Macrolobium acaciifolium* |
| Griffiths et. al. 2011 | *Aldabrachelys gigantea* | *Diospyros egrettarum* |
| Gundlach 1880 | *Trachemys decussata* | *Annona glabra* |
| Gundlach 1880 | *Trachemys decussata* | *Spondias mombin* |
| Guzmán and Stevenson 2008 | *Chelonoidis denticulata* | *Annona* sp. |
| Guzmán and Stevenson 2008 | *Chelonoidis denticulata* | *Astrocaryum murumuru* |
| Guzmán and Stevenson 2008 | *Chelonoidis denticulata* | *Casearia macrocarpa* |
| Guzmán and Stevenson 2008 | *Chelonoidis denticulata* | *Cayaponia ophthalmica* |
| Guzmán and Stevenson 2008 | *Chelonoidis denticulata* | *Cecropia membranacea* |
| Guzmán and Stevenson 2008 | *Chelonoidis denticulata* | *Clarisia racemosa* |
| Guzmán and Stevenson 2008 | *Chelonoidis denticulata* | *Combretum* sp. |
| Guzmán and Stevenson 2008 | *Chelonoidis denticulata* | *Coussapoa* sp. |
| Guzmán and Stevenson 2008 | *Chelonoidis denticulata* | *Duguetia* sp. |
| Guzmán and Stevenson 2008 | *Chelonoidis denticulata* | *Ficus* sp. |
| Guzmán and Stevenson 2008 | *Chelonoidis denticulata* | *Ficus insipida* |
| Guzmán and Stevenson 2008 | *Chelonoidis denticulata* | *Ficus maxima* |
| Guzmán and Stevenson 2008 | *Chelonoidis denticulata* | *Ficus* sp. |
| Guzmán and Stevenson 2008 | *Chelonoidis denticulata* | *Genipa americana* |
| Guzmán and Stevenson 2008 | *Chelonoidis denticulata* | *Geophila repens* |
| Guzmán and Stevenson 2008 | *Chelonoidis denticulata* | *Guatteria* sp. |
| Guzmán and Stevenson 2008 | *Chelonoidis denticulata* | *Helicostylis tomentosa* |
| Guzmán and Stevenson 2008 | *Chelonoidis denticulata* | *Inga* sp. |
| Guzmán and Stevenson 2008 | *Chelonoidis denticulata* | *Iriartea deltoidea* |
| Guzmán and Stevenson 2008 | *Chelonoidis denticulata* | *Jacaratia digitata* |
| Guzmán and Stevenson 2008 | *Chelonoidis denticulata* | *Lecointea amazonica* |
| Guzmán and Stevenson 2008 | *Chelonoidis denticulata* | *Loreya strigosa* |
| Guzmán and Stevenson 2008 | *Chelonoidis denticulata* | *Miconia* sp. |
| Guzmán and Stevenson 2008 | *Chelonoidis denticulata* | *Pourouma cecropiifolia* |
| Guzmán and Stevenson 2008 | *Chelonoidis denticulata* | *Pourouma minor* |
| Guzmán and Stevenson 2008 | *Chelonoidis denticulata* | *Pourouma* sp. |
| Guzmán and Stevenson 2008 | *Chelonoidis denticulata* | *Pouteria* sp. |
| Guzmán and Stevenson 2008 | *Chelonoidis denticulata* | *Pseudolmedia laevis* |
| Guzmán and Stevenson 2008 | *Chelonoidis denticulata* | *Quiina peruviana* |
| Guzmán and Stevenson 2008 | *Chelonoidis denticulata* | *Rollinia* sp. |
| Guzmán and Stevenson 2008 | *Chelonoidis denticulata* | *Salacia gigantea* |
| Guzmán and Stevenson 2008 | *Chelonoidis denticulata* | *Salacia* sp. |
| Guzmán and Stevenson 2008 | *Chelonoidis denticulata* | *Sorocea* sp. |
| Guzmán and Stevenson 2008 | *Chelonoidis denticulata* | *Spondias mombin* |
| Guzmán and Stevenson 2008 | *Chelonoidis denticulata* | *Strychnos* sp. |
| Guzmán and Stevenson 2008 | *Chelonoidis denticulata* | *Tetragastris* sp. |
| Guzmán and Stevenson 2008 | *Chelonoidis denticulata* | *Viola surinamensis* |
| Guzmán and Stevenson 2008 | *Chelonoidis denticulata* | *Mendoncia bivalvis* |
| Guzmán and Stevenson 2008 | *Chelonoidis denticulata* | Unidentified Poaceae 1 |
| Guzmán and Stevenson 2008 | *Chelonoidis denticulata* | Unidentified Poaceae 2 |
| Guzmán and Stevenson 2008 | *Chelonoidis denticulata* | *Brosimum lactescens* |
| Guzmán and Stevenson 2008 | *Chelonoidis denticulata* | *Cecropia sciadophylla* |
| Guzmán and Stevenson 2008 | *Chelonoidis denticulata* | *Ficus* sp1. |
| Guzmán and Stevenson 2008 | *Chelonoidis denticulata* | *Ficus* sp2. |
| Hansen et. al. 2008 | *Aldabrachelys gigantea* | *Rauvolfia micrantha* |
| Hnatiuk 1978 | *Aldabrachelys gigantea* | *Syzygium mamillatum* |
| Hnatiuk 1978 | *Aldabrachelys gigantea* | *Bulbostylis basalis* |
| Hnatiuk 1978 | *Aldabrachelys gigantea* | *Colubrina asiatica* |
| Hnatiuk 1978 | *Aldabrachelys gigantea* | *Cyperus ligularis* |
| Hnatiuk 1978 | *Aldabrachelys gigantea* | *Dactyloctenium ctenoides* |
| Hnatiuk 1978 | *Aldabrachelys gigantea* | *Eragrostis decumbens* |
| Hnatiuk 1978 | *Aldabrachelys gigantea* | *Eragrostis subaequiglumis* |
| Hnatiuk 1978 | *Aldabrachelys gigantea* | *Euphorbia* sp. |
| Hnatiuk 1978 | *Aldabrachelys gigantea* | *Fimbristylis ferruginea* |
| Hnatiuk 1978 | *Aldabrachelys gigantea* | *Guettarda speciosa* |
| Hnatiuk 1978 | *Aldabrachelys gigantea* | *Lepturus repens* |
| Hnatiuk 1978 | *Aldabrachelys gigantea* | *Ochna ciliata* |
| Hnatiuk 1978 | *Aldabrachelys gigantea* | *Phyllanthus maderaspatensis* |
| Hnatiuk 1978 | *Aldabrachelys gigantea* | *Portulaca mauritiensis* |
| Hnatiuk 1978 | *Aldabrachelys gigantea* | *Scaevola taccada* |
| Hnatiuk 1978 | *Aldabrachelys gigantea* | *Serenoa repens* |
| Hnatiuk 1978 | *Aldabrachelys gigantea* | *Solanum americanum* |
| Hnatiuk 1978 | *Aldabrachelys gigantea* | *Sporobolus testudinum* |
| Hnatiuk 1978 | *Aldabrachelys gigantea* | *Sporobolus virginicus* |
| Hnatiuk 1978 | *Aldabrachelys gigantea* | *Stachytarpheta jamaicensis* |
| Hnatiuk 1978 | *Aldabrachelys gigantea* | *Terminalia boivinii* |
| Hnatiuk 1978 | *Aldabrachelys gigantea* | *Thespesia populnea* |
| Hnatiuk 1978 | *Aldabrachelys gigantea* | *Dactyloctenium pilosum* |
| Hnatiuk 1978 | *Aldabrachelys gigantea* | *Hedyotis prolifera* |
| Hnatiuk 1978 | *Aldabrachelys gigantea* | *Pycnus pumilus* |
| Hnatiuk 1978 | *Aldabrachelys gigantea* | *Ficus sundaica* |
| Hnatiuk 1978 | *Aldabrachelys gigantea* | *Gouania tiliifolia* |
| Hnatiuk 1978 | *Aldabrachelys gigantea* | *Cassine aethiopica* |
| Iftime and Iftime 2012 | *Testudo graeca* | *Nesogenes prostrata* |
| Iftime and Iftime 2012 | *Testudo graeca* | *Cornus mas* |
| Iftime and Iftime 2012 | *Testudo graeca* | *Prunus* sp. |
| Jerozolimski et. al. 2009 | *Chelonoidis denticulata* | *Pyrus* sp. |
| Jerozolimski et. al. 2009 | *Chelonoidis denticulata* | Unidentified Sapotaceae |
| Jerozolimski et. al. 2009 | *Chelonoidis denticulata* | *Ananas ananassoides* |
| Jerozolimski et. al. 2009 | *Chelonoidis denticulata* | *Attalea maripa* |
| Jerozolimski et. al. 2009 | *Chelonoidis denticulata* | *Brosimum lactescens* |
| Jerozolimski et. al. 2009 | *Chelonoidis denticulata* | *Castilla ulei* |
| Jerozolimski et. al. 2009 | *Chelonoidis denticulata* | *Cecropia* sp. |
| Jerozolimski et. al. 2009 | *Chelonoidis denticulata* | *Celtis* sp. |
| Jerozolimski et. al. 2009 | *Chelonoidis denticulata* | *Clarisia ilicifolia* |
| Jerozolimski et. al. 2009 | *Chelonoidis denticulata* | *Eugenia* sp. |
| Jerozolimski et. al. 2009 | *Chelonoidis denticulata* | *Ficus* sp1. |
| Jerozolimski et. al. 2009 | *Chelonoidis denticulata* | *Ficus* sp2. |
| Jerozolimski et. al. 2009 | *Chelonoidis denticulata* | *Genipa americana* |
| Jerozolimski et. al. 2009 | *Chelonoidis denticulata* | *Geophila cordifolia* |
| Jerozolimski et. al. 2009 | *Chelonoidis denticulata* | *Guettarda* sp. |
| Jerozolimski et. al. 2009 | *Chelonoidis denticulata* | *Helicostylis tomentosa* |
| Jerozolimski et. al. 2009 | *Chelonoidis denticulata* | *Inga* sp. |
| Jerozolimski et. al. 2009 | *Chelonoidis denticulata* | *Jacaratia spinosa* |
| Jerozolimski et. al. 2009 | *Chelonoidis denticulata* | *Mouriri* sp. |
| Jerozolimski et. al. 2009 | *Chelonoidis denticulata* | *Pourouma guianensis* |
| Jerozolimski et. al. 2009 | *Chelonoidis denticulata* | *Pourouma* sp. |
| Jerozolimski et. al. 2009 | *Chelonoidis denticulata* | *Pouteria macrophylla* |
| Jerozolimski et. al. 2009 | *Chelonoidis denticulata* | *Protium* sp. |
| Jerozolimski et. al. 2009 | *Chelonoidis denticulata* | *Psidium* sp. |
| Jerozolimski et. al. 2009 | *Chelonoidis denticulata* | *Quiina paraensis* |
| Jerozolimski et. al. 2009 | *Chelonoidis denticulata* | *Scleria* sp. |
| Jerozolimski et. al. 2009 | *Chelonoidis denticulata* | *Spondias mombin* |
| Jerozolimski et. al. 2009 | *Chelonoidis denticulata* | *Tetragastris altissima* |
| Jerozolimski et. al. 2009 | *Chelonoidis denticulata* | *Xylopia amazonica* |
| Jerozolimski et. al. 2009 | *Chelonoidis denticulata* | Unidentified Myrtaceae |
| Joshua et. al. 2010 | *Chersina angulata* | *Genipa americana* |
| Kennet and Russel-Smith 1993 | *Elseya dentata* | *Nylandtia spinosa* |
| Kennet and Russel-Smith 1993 | *Elseya dentata* | *Ficus racemosa* |
| Kennet and Russel-Smith 1993 | *Elseya dentata* | *Nauclea orientalis* |
| Kennet and Russel-Smith 1993 | *Elseya dentata* | *Pandanus aquaticus* |
| Kennet and Russel-Smith 1993 | *Elseya dentata* | *Syzygium forte* |
| Kennet and Russel-Smith 1993 | *Elseya dentata* | *Terminalia erythrocarpa* |
| Kennet and Russel-Smith 1993 | *Elseya dentata* | *Terminalia microcarpa* |
| Kennet and Tory 1996 | *Elseya dentata* | *Cyclophyllum schultzei* |
| Kennet and Tory 1996 | *Elseya dentata* | *Acacia auriculiformis* |
| Kennet and Tory 1996 | *Elseya dentata* | *Carallia brachiata* |
| Kennet and Tory 1996 | *Elseya dentata* | *Ficus racemosa* |
| Kennet and Tory 1996 | *Elseya dentata* | *Morinda citrifolia* |
| Kennet and Tory 1996 | *Elseya dentata* | *Nauclea orientalis* |
| Kennet and Tory 1996 | *Elseya dentata* | *Pandanus aquaticus* |
| Kennet and Tory 1996 | *Elseya dentata* | *Terminalia erythrocarpa* |
| Kimmons and Moll 2010 | *Chelydra serpentina* | *Ambrosia* sp. |
| Kimmons and Moll 2010 | *Chelydra serpentina* | *Echinochloa crus-galli* |
| Kimmons and Moll 2010 | *Chelydra serpentina* | *Elymus repens* |
| Kimmons and Moll 2010 | *Chelydra serpentina* | *Morus* sp. |
| Kimmons and Moll 2010 | *Chelydra serpentina* | *Panicum* sp. |
| Kimmons and Moll 2010 | *Chelydra serpentina* | *Polygonum* sp. |
| Kimmons and Moll 2010 | *Chelydra serpentina* | *Ranunculus sceleratus* |
| Kimmons and Moll 2010 | *Trachemys scripta* | *Rumex crispus* |
| Kimmons and Moll 2010 | *Trachemys scripta* | *Echinochloa crus-galli* |
| Kimmons and Moll 2010 | *Trachemys scripta* | *Morus* sp. |
| Kimmons and Moll 2010 | *Trachemys scripta* | *Polygonum* sp. |
| Kimmons and Moll 2010 | *Trachemys scripta* | *Ranunculus sceleratus* |
| Kimmons and Moll 2010 | *Trachemys scripta* | *Rumex crispus* |
| Kimmons and Moll 2010 | *Trachemys scripta* | *Rumex obtusifolius* |
| Kimmons and Moll 2010 | *Trachemys scripta* | *Setaria verticillata* |
| Kimmons and Moll 2010 | *Chelydra serpentina* | *Silene nocturna* |
| Kimmons and Moll 2010 | *Chelydra serpentina* | *Echinochloa crus-galli* |
| Kimmons and Moll 2010 | *Chelydra serpentina* | *Morus* sp. |
| Kimmons and Moll 2010 | *Trachemys scripta* | *Rumex crispus* |
| Kimmons and Moll 2010 | *Trachemys scripta* | *Echinochloa crus-galli* |
| Kimmons and Moll 2010 | *Trachemys scripta* | *Morus* sp. |
| Klimstra and Newsome 1960 | *Terrapene carolina* | *Rumex crispus* |
| Klimstra and Newsome 1960 | *Terrapene carolina* | *Hordeum* sp. |
| Klimstra and Newsome 1960 | *Terrapene carolina* | *Bromus* sp. |
| Klimstra and Newsome 1960 | *Terrapene carolina* | *Paspalum* sp. |
| Klimstra and Newsome 1960 | *Terrapene carolina* | *Rubus* sp. |
| Klimstra and Newsome 1960 | *Terrapene carolina* | *Prunus* sp. |
| Klimstra and Newsome 1960 | *Terrapene carolina* | *Fragaria* sp. |
| Klimstra and Newsome 1960 | *Terrapene carolina* | *Diospyros virginiana* |
| Klimstra and Newsome 1960 | *Terrapene carolina* | *Morus rubra* |
| Klimstra and Newsome 1960 | *Terrapene carolina* | *Polygonum* sp. |
| Klimstra and Newsome 1960 | *Terrapene carolina* | Unidentified Caryophyllaceae |
| Klimstra and Newsome 1960 | *Terrapene carolina* | *Galium* sp. |
| Klimstra and Newsome 1960 | *Terrapene carolina* | *Ambrosia* sp. |
| Klimstra and Newsome 1960 | *Terrapene carolina* | Unidentified Chenopodiaceae |
| Klimstra and Newsome 1960 | *Terrapene carolina* | Unidentified Cyperaceae |
| Klimstra and Newsome 1960 | *Terrapene carolina* | *Vitis* sp. |
| Kuchling and Bloxam 1988 | *Pyxis planicauda* | *Breonia perrieri* |
| Kuchling and Bloxam 1988 | *Pyxis planicauda* | *Broussonetia greveana* |
| Lagler 1943 | *Chelydra serpentina* | *Nymphaea odorata* |
| Lagler 1943 | *Chelydra serpentina* | *Bidens* sp. |
| Lagler 1943 | *Chrysemys picta* | *Nymphaea odorata* |
| Lagler 1943 | *Chrysemys picta* | *Triticum* sp. |
| Lagler 1943 | *Chrysemys picta* | *Zea mays* |
| Lagler 1943 | *Emys blandingii* | *Cornus amomum* |
| Lagler 1943 | *Emys blandingii* | *Bidens* sp. |
| Lagler 1943 | *Graptemys geographica* | *Potamogeton* sp. |
| Lagler 1943 | *Sternotherus odoratus* | *Cornus amomum* |
| Lagler 1943 | *Sternotherus odoratus* | *Nuphar* sp. |
| Lagler 1943 | *Sternotherus odoratus* | *Nymphaea odorata* |
| Lagler 1943 | *Sternotherus odoratus* | *Bidens* sp. |
| Lambiris et. al. 1989 | *Kinixys spekii* | *Uapaca kirkiana* |
| Lautenschlager-Rodrigues 2016 | *Chelonoidis carbonaria* | *Abuta selloana* |
| Lautenschlager-Rodrigues 2016 | *Chelonoidis carbonaria* | *Acrocomia aculeata* |
| Lautenschlager-Rodrigues 2016 | *Chelonoidis carbonaria* | *Archontophoenix cunninghamiana* |
| Lautenschlager-Rodrigues 2016 | *Chelonoidis carbonaria* | *Artocarpus heterophyllus* |
| Lautenschlager-Rodrigues 2016 | *Chelonoidis carbonaria* | *Casearia sylvestris* |
| Lautenschlager-Rodrigues 2016 | *Chelonoidis carbonaria* | *Cecropia pachystachya* |
| Lautenschlager-Rodrigues 2016 | *Chelonoidis carbonaria* | *Clusia criuva* |
| Lautenschlager-Rodrigues 2016 | *Chelonoidis carbonaria* | *Cordia ecalyculata* |
| Lautenschlager-Rodrigues 2016 | *Chelonoidis carbonaria* | *Cryptocarya mandioccana* |
| Lautenschlager-Rodrigues 2016 | *Chelonoidis carbonaria* | *Eriobotrya japonica* |
| Lautenschlager-Rodrigues 2016 | *Chelonoidis carbonaria* | *Eugenia uniflora* |
| Lautenschlager-Rodrigues 2016 | *Chelonoidis carbonaria* | *Euterpe edulis* |
| Lautenschlager-Rodrigues 2016 | *Chelonoidis carbonaria* | *Genipa americana* |
| Lautenschlager-Rodrigues 2016 | *Chelonoidis carbonaria* | *Hymenaea courbaril* |
| Lautenschlager-Rodrigues 2016 | *Chelonoidis carbonaria* | *Inga* sp. |
| Lautenschlager-Rodrigues 2016 | *Chelonoidis carbonaria* | *Jacaratia spinosa* |
| Lautenschlager-Rodrigues 2016 | *Chelonoidis carbonaria* | *Licuala grandis* |
| Lautenschlager-Rodrigues 2016 | *Chelonoidis carbonaria* | *Litchi chinensis* |
| Lautenschlager-Rodrigues 2016 | *Chelonoidis carbonaria* | *Malpighia* sp. |
| Lautenschlager-Rodrigues 2016 | *Chelonoidis carbonaria* | *Melia azedarach* |
| Lautenschlager-Rodrigues 2016 | *Chelonoidis carbonaria* | *Ocotea catharinensis* |
| Lautenschlager-Rodrigues 2016 | *Chelonoidis carbonaria* | *Philodendron bipinnatifidum* |
| Lautenschlager-Rodrigues 2016 | *Chelonoidis carbonaria* | *Plinia cauliflora* |
| Lautenschlager-Rodrigues 2016 | *Chelonoidis carbonaria* | *Psidium cattleianum* |
| Lautenschlager-Rodrigues 2016 | *Chelonoidis carbonaria* | *Quiina glaziovii* |
| Lautenschlager-Rodrigues 2016 | *Chelonoidis carbonaria* | *Sabal maritima* |
| Lautenschlager-Rodrigues 2016 | *Chelonoidis carbonaria* | *Spondias purpurea* |
| Lautenschlager-Rodrigues 2016 | *Chelonoidis carbonaria* | *Syagrus oleracea* |
| Lautenschlager-Rodrigues 2016 | *Chelonoidis carbonaria* | *Syagrus romanzoffiana* |
| Lautenschlager-Rodrigues 2016 | *Chelonoidis carbonaria* | *Syzygium cumini* |
| Lautenschlager-Rodrigues 2016 | *Chelonoidis carbonaria* | *Talisia esculenta* |
| Lautenschlager-Rodrigues 2016 | *Chelonoidis carbonaria* | *Hyophorbe indica* |
| Lautenschlager-Rodrigues 2016 | *Chelonoidis carbonaria* | *Cordia africana* |
| Leuteritz 2003 | *Astrochelys radiata* | *Clerodendrum perrieri* |
| Leuteritz 2003 | *Astrochelys radiata* | *Diospyros myriophylla* |
| Leuteritz 2003 | *Astrochelys radiata* | *Operculicarya pachypus* |
| Leuteritz 2003 | *Astrochelys radiata* | *Opuntia* sp. |
| Leuteritz 2003 | *Astrochelys radiata* | *Lycium acutifolium* |
| Leuteritz 2003 | *Astrochelys radiata* | *Sclerocarya birrea* |
| Leuteritz 2003 | *Astrochelys radiata* | *Tetraena madagascariensis* |
| Limpus and Limpus 2000 | *Chelonia mydas* | *Avicennia marina* |
| Limpus and Limpus 2000 | *Chelonia mydas* | *Rhizophora* sp. |
| Lingard et. al. 2003 | *Astrochelys radiata* | *Operculicarya decarya* |
| Lingard et. al. 2003 | *Astrochelys radiata* | *Opuntia ficus-indica* |
| Lingard et. al. 2003 | *Astrochelys radiata* | *Opuntia monacantha* |
| Liu et. al. 2004 | *Terrapene carolina* | *Annona glabra* |
| Liu et. al. 2004 | *Terrapene carolina* | *Coccoloba uvifera* |
| Liu et. al. 2004 | *Terrapene carolina* | *Coccothrinax argentata* |
| Liu et. al. 2004 | *Terrapene carolina* | *Ficus* sp. |
| Liu et. al. 2004 | *Terrapene carolina* | *Manilkara zapota* |
| Liu et. al. 2004 | *Terrapene carolina* | *Morinda royoc* |
| Liu et. al. 2004 | *Terrapene carolina* | *Mosiera longipes* |
| Liu et. al. 2004 | *Terrapene carolina* | *Paspalum* sp. |
| Liu et. al. 2004 | *Terrapene carolina* | *Smilax havanensis* |
| Liu et. al. 2004 | *Terrapene carolina* | Unidentified Fabaceae |
| Liu et. al. 2004 | *Terrapene carolina* | *Byrsonima lucida* |
| Liu et. al. 2004 | *Terrapene carolina* | *Leucothrinax morrisii* |
| Liu et. al. 2004 | *Terrapene carolina* | *Setaria* sp. |
| Loehr 2002 | *Homopus signatus* | *Antizoma* sp. |
| Loehr 2002 | *Homopus signatus* | *Crassula thunbergiana* |
| Loehr 2002 | *Homopus signatus* | *Grielum humifusum* |
| Loehr 2002 | *Homopus signatus* | *Heliophila variabilis* |
| Loehr 2002 | *Homopus signatus* | *Oxalis* sp. |
| Macip-Rios et. al. 2010 | *Kinosternon integrum* | *Argemone ochroleuca* |
| Macip-Rios et. al. 2010 | *Kinosternon integrum* | *Lemna* sp. |
| Macip-Rios et. al. 2010 | *Kinosternon integrum* | *Psidium* sp. |
| Macip-Rios et. al. 2010 | *Kinosternon integrum* | Unidentified Poaceae |
| Milton 1992 | *Stigmochelys pardalis* | *Amaranthus* sp. |
| Milton 1992 | *Stigmochelys pardalis* | *Aptosimum indivisum* |
| Milton 1992 | *Stigmochelys pardalis* | *Argemone mexicana* |
| Milton 1992 | *Stigmochelys pardalis* | *Aristida* sp. |
| Milton 1992 | *Stigmochelys pardalis* | *Atriplex lindleyi* |
| Milton 1992 | *Stigmochelys pardalis* | *Atriplex semibaccata* |
| Milton 1992 | *Stigmochelys pardalis* | *Chamaesyce inequilatera* |
| Milton 1992 | *Stigmochelys pardalis* | *Chenopodium* sp. |
| Milton 1992 | *Stigmochelys pardalis* | *Chrysocoma ciliata* |
| Milton 1992 | *Stigmochelys pardalis* | *Crassula subaphylla* |
| Milton 1992 | *Stigmochelys pardalis* | *Cuspidia cernua* |
| Milton 1992 | *Stigmochelys pardalis* | *Enneapogon desvauxii* |
| Milton 1992 | *Stigmochelys pardalis* | *Enneapogon scaber* |
| Milton 1992 | *Stigmochelys pardalis* | *Eragrostis obtusa* |
| Milton 1992 | *Stigmochelys pardalis* | *Euphorbia* sp. |
| Milton 1992 | *Stigmochelys pardalis* | *Galenia papulosa* |
| Milton 1992 | *Stigmochelys pardalis* | *Heliophila* sp. |
| Milton 1992 | *Stigmochelys pardalis* | *Hermannia* sp. |
| Milton 1992 | *Stigmochelys pardalis* | *Hypertelis salsoloides* |
| Milton 1992 | *Stigmochelys pardalis* | *Lepidium* sp. |
| Milton 1992 | *Stigmochelys pardalis* | *Lessertia annularis* |
| Milton 1992 | *Stigmochelys pardalis* | *Leysera tenella* |
| Milton 1992 | *Stigmochelys pardalis* | *Limeum aethiopicum* |
| Milton 1992 | *Stigmochelys pardalis* | *Lolium* sp. |
| Milton 1992 | *Stigmochelys pardalis* | *Lotononis* sp. |
| Milton 1992 | *Stigmochelys pardalis* | *Malva parviflora* |
| Milton 1992 | *Stigmochelys pardalis* | *Medicago polymorpha* |
| Milton 1992 | *Stigmochelys pardalis* | *Nemesia* sp. |
| Milton 1992 | *Stigmochelys pardalis* | *Osteospermum calendulaceum* |
| Milton 1992 | *Stigmochelys pardalis* | *Pleiospilos compactus* |
| Milton 1992 | *Stigmochelys pardalis* | *Polygonum* sp. |
| Milton 1992 | *Stigmochelys pardalis* | *Sida hederifolia* |
| Milton 1992 | *Stigmochelys pardalis* | *Tetragonia echinata* |
| Milton 1992 | *Stigmochelys pardalis* | *Tetragonia spicata* |
| Milton 1992 | *Stigmochelys pardalis* | *Thesium lineatum* |
| Milton 1992 | *Stigmochelys pardalis* | *Tragus* sp. |
| Milton 1992 | *Stigmochelys pardalis* | *Trianthema triquetra* |
| Milton 1992 | *Stigmochelys pardalis* | *Tribulus terrestris* |
| Milton 1992 | *Stigmochelys pardalis* | *Walafrida* sp. |
| Milton 1992 | *Stigmochelys pardalis* | *Eriocephalus* sp. |
| Milton 1992 | *Stigmochelys pardalis* | *Ursinia* sp. |
| Milton 1992 | *Stigmochelys pardalis* | *Tribolium purpureum* |
| Milton 1992 | *Stigmochelys pardalis* | Unidentified Cyperaceae |
| Moldowan et. al. 2016 | *Chelydra serpentina* | *Nuphar variegata* |
| Moll 1976 | *Graptemys ouachitensis* | *Ulmus americana* |
| Moll 1989 | *Dermatemys mawii* | *Ficus* sp. |
| Moll 1989 | *Dermatemys mawii* | *Ficus obtusiuscula* |
| Moll and Jansen 1995 | *Rhinoclemmys annulata* | *Astrocaryum alatum* |
| Moll and Jansen 1995 | *Rhinoclemmys annulata* | *Faramea suerrensis* |
| Moll and Jansen 1995 | *Rhinoclemmys annulata* | *Jacaratia dolichaula* |
| Moll and Jansen 1995 | *Rhinoclemmys annulata* | *Solanum siparunoides* |
| Moll and Jansen 1995 | *Rhinoclemmys funerea* | *Artocarpus altilis* |
| Moll and Jansen 1995 | *Rhinoclemmys funerea* | *Eichhornia crassipes* |
| Moll and Jansen 1995 | *Rhinoclemmys funerea* | *Ficus* sp. |
| Moll and Jansen 1995 | *Rhinoclemmys funerea* | *Miconia* sp. |
| Moll and Jansen 1995 | *Rhinoclemmys funerea* | *Passiflora foetida* |
| Moll and Jansen 1995 | *Rhinoclemmys annulata* | *Faramea suerrensis* |
| Moll and Jansen 1995 | *Rhinoclemmys annulata* | *Ficus* sp. |
| Moll and Jansen 1995 | *Rhinoclemmys annulata* | *Jacaratia dolichaula* |
| Moll and Jansen 1995 | *Rhinoclemmys annulata* | *Miconia affinis* |
| Moll and Jansen 1995 | *Rhinoclemmys annulata* | *Sonneratia* sp. |
| Moll and Jansen 1995 | *Rhinoclemmys funerea* | *Cecropia* sp. |
| Moll and Jansen 1995 | *Rhinoclemmys funerea* | *Dieffenbachia longispatha* |
| Moll and Jansen 1995 | *Rhinoclemmys funerea* | *Ficus insipida* |
| Moll and Jansen 1995 | *Rhinoclemmys funerea* | *Ipomoea trifida* |
| Moll and Jansen 1995 | *Rhinoclemmys funerea* | *Solanum pimpinellifolium* |
| Moll and Jansen 1995 | *Rhinoclemmys funerea* | *Spondias mombin* |
| Moolna 2007 | *Aldabrachelys gigantea* | *Diospyros egrettarum* |
| Moskovits and Bjorndal 1990 | *Chelonoidis carbonaria* | *Annona* sp1. |
| Moskovits and Bjorndal 1990 | *Chelonoidis carbonaria* | *Annona* sp2. |
| Moskovits and Bjorndal 1990 | *Chelonoidis carbonaria* | *Bagassa guianensis* |
| Moskovits and Bjorndal 1990 | *Chelonoidis carbonaria* | *Duguetia surinamensis* |
| Moskovits and Bjorndal 1990 | *Chelonoidis carbonaria* | *Ecclinusa guianensis* |
| Moskovits and Bjorndal 1990 | *Chelonoidis carbonaria* | *Ficus* sp. |
| Moskovits and Bjorndal 1990 | *Chelonoidis carbonaria* | *Genipa americana* |
| Moskovits and Bjorndal 1990 | *Chelonoidis carbonaria* | *Geophila repens* |
| Moskovits and Bjorndal 1990 | *Chelonoidis carbonaria* | *Guettarda argentea* |
| Moskovits and Bjorndal 1990 | *Chelonoidis carbonaria* | *Mauritia flexuosa* |
| Moskovits and Bjorndal 1990 | *Chelonoidis carbonaria* | *Myriaspore egensis* |
| Moskovits and Bjorndal 1990 | *Chelonoidis carbonaria* | *Passiflora* sp. |
| Moskovits and Bjorndal 1990 | *Chelonoidis carbonaria* | *Philodendron* sp. |
| Moskovits and Bjorndal 1990 | *Chelonoidis carbonaria* | *Spondias mombin* |
| Moskovits and Bjorndal 1990 | *Chelonoidis carbonaria* | *Trattinnickia ravifolia* |
| Moskovits and Bjorndal 1990 | *Chelonoidis carbonaria* | *Anacardium giganteum* |
| Moskovits and Bjorndal 1990 | *Chelonoidis carbonaria* | *Bagassa guianensis* |
| Moskovits and Bjorndal 1990 | *Chelonoidis carbonaria* | *Bromelia* sp. |
| Moskovits and Bjorndal 1990 | *Chelonoidis carbonaria* | *Brosimum potabile* |
| Moskovits and Bjorndal 1990 | *Chelonoidis carbonaria* | *Clavija* sp. |
| Moskovits and Bjorndal 1990 | *Chelonoidis carbonaria* | *Desmoncus polyacanthos* |
| Moskovits and Bjorndal 1990 | *Chelonoidis carbonaria* | *Duguetia surinamensis* |
| Moskovits and Bjorndal 1990 | *Chelonoidis carbonaria* | *Duroia eriopila* |
| Moskovits and Bjorndal 1990 | *Chelonoidis carbonaria* | *Ficus* sp. |
| Moskovits and Bjorndal 1990 | *Chelonoidis carbonaria* | *Guettarda argentea* |
| Moskovits and Bjorndal 1990 | *Chelonoidis carbonaria* | *Licania kunthiana* |
| Moskovits and Bjorndal 1990 | *Chelonoidis carbonaria* | *Mauritia flexuosa* |
| Moskovits and Bjorndal 1990 | *Chelonoidis carbonaria* | *Myriaspore egensis* |
| Moskovits and Bjorndal 1990 | *Chelonoidis carbonaria* | *Passiflora coccinea* |
| Moskovits and Bjorndal 1990 | *Chelonoidis carbonaria* | *Passiflora vespertilio* |
| Moskovits and Bjorndal 1990 | *Chelonoidis carbonaria* | *Philodendron* sp. |
| Moskovits and Bjorndal 1990 | *Chelonoidis carbonaria* | *Posoqueria* sp. |
| Moskovits and Bjorndal 1990 | *Chelonoidis carbonaria* | *Pouteria* sp. |
| Moskovits and Bjorndal 1990 | *Chelonoidis carbonaria* | *Pradosia* sp. |
| Moskovits and Bjorndal 1990 | *Chelonoidis carbonaria* | *Spondias mombin* |
| Moskovits and Bjorndal 1990 | *Chelonoidis carbonaria* | *Richardela* sp. |
| Moskovits and Bjorndal 1990 | *Chelonoidis carbonaria* | *Ecclinusa guianensis* |
| Moskovits and Bjorndal 1990 | *Chelonoidis carbonaria* | Unidentified Lecythidaceae |
| Moskovits and Bjorndal 1990 | *Chelonoidis denticulata* | *Annona* sp1. |
| Moskovits and Bjorndal 1990 | *Chelonoidis denticulata* | *Annona* sp2. |
| Moskovits and Bjorndal 1990 | *Chelonoidis denticulata* | *Bagassa guianensis* |
| Moskovits and Bjorndal 1990 | *Chelonoidis denticulata* | *Duguetia surinamensis* |
| Moskovits and Bjorndal 1990 | *Chelonoidis denticulata* | *Ecclinusa guianensis* |
| Moskovits and Bjorndal 1990 | *Chelonoidis denticulata* | *Ficus* sp. |
| Moskovits and Bjorndal 1990 | *Chelonoidis denticulata* | *Genipa americana* |
| Moskovits and Bjorndal 1990 | *Chelonoidis denticulata* | *Geophila repens* |
| Moskovits and Bjorndal 1990 | *Chelonoidis denticulata* | *Guettarda argentea* |
| Moskovits and Bjorndal 1990 | *Chelonoidis denticulata* | *Mauritia flexuosa* |
| Moskovits and Bjorndal 1990 | *Chelonoidis denticulata* | *Myriaspore egensis* |
| Moskovits and Bjorndal 1990 | *Chelonoidis denticulata* | *Passiflora* sp. |
| Moskovits and Bjorndal 1990 | *Chelonoidis denticulata* | *Philodendron* sp. |
| Moskovits and Bjorndal 1990 | *Chelonoidis denticulata* | *Spondias mombin* |
| Moskovits and Bjorndal 1990 | *Chelonoidis denticulata* | *Trattinnickia* sp. |
| Moskovits and Bjorndal 1990 | *Chelonoidis denticulata* | *Anacardium giganteum* |
| Moskovits and Bjorndal 1990 | *Chelonoidis denticulata* | *Bagassa guianensis* |
| Moskovits and Bjorndal 1990 | *Chelonoidis denticulata* | *Bromelia* sp. |
| Moskovits and Bjorndal 1990 | *Chelonoidis denticulata* | *Brosimum potabile* |
| Moskovits and Bjorndal 1990 | *Chelonoidis denticulata* | *Clavijsa* sp. |
| Moskovits and Bjorndal 1990 | *Chelonoidis denticulata* | *Desmoncus polyacanthos* |
| Moskovits and Bjorndal 1990 | *Chelonoidis denticulata* | *Duguetia surinamensis* |
| Moskovits and Bjorndal 1990 | *Chelonoidis denticulata* | *Duroia eriopila* |
| Moskovits and Bjorndal 1990 | *Chelonoidis denticulata* | *Ficus* sp. |
| Moskovits and Bjorndal 1990 | *Chelonoidis denticulata* | *Guettarda argentea* |
| Moskovits and Bjorndal 1990 | *Chelonoidis denticulata* | *Licania kunthiana* |
| Moskovits and Bjorndal 1990 | *Chelonoidis denticulata* | *Mauritia flexuosa* |
| Moskovits and Bjorndal 1990 | *Chelonoidis denticulata* | *Myriasporea egensis* |
| Moskovits and Bjorndal 1990 | *Chelonoidis denticulata* | *Passiflora coccinea* |
| Moskovits and Bjorndal 1990 | *Chelonoidis denticulata* | *Passiflora vespertilio* |
| Moskovits and Bjorndal 1990 | *Chelonoidis denticulata* | *Philodendron* sp. |
| Moskovits and Bjorndal 1990 | *Chelonoidis denticulata* | *Posoqueria* sp. |
| Moskovits and Bjorndal 1990 | *Chelonoidis denticulata* | *Pouteria* sp. |
| Moskovits and Bjorndal 1990 | *Chelonoidis denticulata* | *Pradosia* sp. |
| Moskovits and Bjorndal 1990 | *Chelonoidis denticulata* | *Spondias mombin* |
| Moskovits and Bjorndal 1990 | *Chelonoidis denticulata* | *Richardella* sp. |
| Moskovits and Bjorndal 1990 | *Chelonoidis denticulata* | *Ecclinusa guianensis* |
| Moskovits and Bjorndal 1990 | *Chelonoidis denticulata* | Unidentified Lecythydaceae |
| Murray and Wolf 2013 | *Gopherus agassizii* | *Opuntia* sp. |
| Padgett et. al. 2012 | *Chrysemys picta* | *Carex* sp. |
| Padgett et. al. 2012 | *Chrysemys picta* | *Decodon verticillatus* |
| Padgett et. al. 2012 | *Chrysemys picta* | *Najas flexilis* |
| Padgett et. al. 2012 | *Chrysemys picta* | *Nuphar variegata* |
| Padgett et. al. 2012 | *Chrysemys picta* | *Nymphaea odorata* |
| Padgett et. al. 2012 | *Chrysemys picta* | *Potamogeton* sp. |
| Padgett et. al. 2012 | *Chrysemys picta* | *Bidens* sp. |
| Padgett et. al. 2012 | *Chrysemys picta* | Unidentified Poaceae |
| Pemberton and Gilchrist 2009 | *Aldabrachelys gigantea* | *Artocarpus altilis* |
| Pemberton and Gilchrist 2009 | *Aldabrachelys gigantea* | *Xylocarpus moluccensis* |
| Perez-Eman and Paolillo 1997 | *Peltocephalus dumerilianus* | *Annona glabra* |
| Perez-Eman and Paolillo 1997 | *Peltocephalus dumerilianus* | *Combretum laxum* |
| Perez-Eman and Paolillo 1997 | *Peltocephalus dumerilianus* | *Eperua purpurea* |
| Perez-Eman and Paolillo 1997 | *Peltocephalus dumerilianus* | *Hevea benthamiana* |
| Perez-Eman and Paolillo 1997 | *Peltocephalus dumerilianus* | *Leopoldinia piassaba* |
| Perez-Eman and Paolillo 1997 | *Peltocephalus dumerilianus* | *Leopoldinia pulchra* |
| Perez-Eman and Paolillo 1997 | *Peltocephalus dumerilianus* | *Macrolobium multijugum* |
| Perez-Eman and Paolillo 1997 | *Peltocephalus dumerilianus* | *Macrolobium* sp. |
| Perez-Eman and Paolillo 1997 | *Peltocephalus dumerilianus* | *Maripa paniculata* |
| Perez-Eman and Paolillo 1997 | *Peltocephalus dumerilianus* | *Mauritia flexuosa* |
| Perez-Eman and Paolillo 1997 | *Peltocephalus dumerilianus* | *Mauritiella aculeata* |
| Perez-Eman and Paolillo 1997 | *Peltocephalus dumerilianus* | *Parahancornia negroensis* |
| Perez-Eman and Paolillo 1997 | *Peltocephalus dumerilianus* | *Parinari campestris* |
| Perez-Eman and Paolillo 1997 | *Peltocephalus dumerilianus* | *Rauvolfia polyphylla* |
| Perez-Eman and Paolillo 1997 | *Peltocephalus dumerilianus* | *Swartzia sericea* |
| Perez-Eman and Paolillo 1997 | *Peltocephalus dumerilianus* | *Compsiandra comosa* |
| Perez-Eman and Paolillo 1997 | *Peltocephalus dumerilianus* | *Macrolobium angustifolium* |
| Perez-Eman and Paolillo 1997 | *Peltocephalus dumerilianus* | Unidentified Lauraceae |
| Platt et. al. 2009 | *Terrapene carolina bauri* | *Annona glabra* |
| Platt et. al. 2009 | *Terrapene carolina bauri* | *Byrsonima lucida* |
| Platt et. al. 2009 | *Terrapene carolina bauri* | *Coccoloba uvifera* |
| Platt et. al. 2009 | *Terrapene carolina bauri* | *Coccothrinax argentata* |
| Platt et. al. 2009 | *Terrapene carolina bauri* | *Ficus* sp. |
| Platt et. al. 2009 | *Terrapene carolina bauri* | *Morinda royoc* |
| Platt et. al. 2009 | *Terrapene carolina bauri* | *Mosiera longipes* |
| Platt et. al. 2009 | *Terrapene carolina bauri* | *Socratea exorrhiza* |
| Platt et. al. 2009 | *Terrapene carolina bauri* | *Leucothrinax morrisii* |
| Platt et. al. 2009 | *Terrapene carolina bauri* | Unidentified Fabaceae |
| Platt et. al. 2010 | *Heosemys depressa* | *Ficus hispida* |
| Platt et. al. 2010 | *Heosemys depressa* | *Erythrina suberosa* |
| Platt et. al. 2010 | *Heosemys depressa* | *Grewia nervosa* |
| Platt et. al. 2014a | *Vijayachelys silvatica* | *Dillenia pentagyna* |
| Platt et. al. 2014a | *Vijayachelys silvatica* | *Salacca* sp. |
| Platt et. al. 2014b | *Heosemys depressa* | *Dillenia pentagyna* |
| Platt et. al. 2016 | *Kinosternon hirtipes* | *Paspalum distichum* |
| Platt et. al. 2016 | *Kinosternon hirtipes* | *Prosopis glandulosa* |
| Plummer and Farrar 1981 | *Apalone mutica* | *Morus* sp. |
| Plummer and Farrar 1981 | *Apalone mutica* | *Populus deltoides* |
| Plummer and Farrar 1981 | *Apalone mutica* | *Morus* sp. |
| Plummer and Farrar 1981 | *Apalone mutica* | *Populus* sp. |
| Raney and Rachner 194 2 | *Chrysemys picta* | *Nuphar variegata* |
| Rasoma et. al. 2013 | *Astrochelys radiata* | *Gyrocarpus americanus* |
| Rasoma et. al. 2013 | *Astrochelys radiata* | *Paederia grandidieri* |
| Rasoma et. al. 2013 | *Astrochelys radiata* | *Radamaea montana* |
| Rasoma et. al. 2013 | *Astrochelys radiata* | *Salvadora angustifolia* |
| Rasoma et. al. 2013 | *Astrochelys radiata* | *Olex dissitiflora* |
| Renvoize 1971 | *Aldabrachelys gigantea* | *Pandanus tectorius* |
| Rick and Bowman 1961 | *Chelonoidis porteri* | *Solanum siparunoides* |
| Rouag et. al. 2008 | *Testudo graeca* | *Anagallis minima* |
| Rouag et. al. 2008 | *Testudo graeca* | *Linaria pinifolia* |
| Rouag et. al. 2008 | *Testudo graeca* | *Tuberaria guttata* |
| Rouag et. al. 2008 | *Testudo graeca* | *Coronilla scorpioides* |
| Rust and Roth 1981 | *Terrapene carolina* | *Podophyllum peltatum* |
| Santos-Júnior 2009 | *Podocnemis erythrocephala* | *Pouteria* sp. |
| Santos-Júnior 2009 | *Podocnemis erythrocephala* | *Posoqueria* sp. |
| Santos-Júnior 2009 | *Podocnemis erythrocephala* | *Smilax coriacea* |
| Santos-Júnior 2009 | *Podocnemis erythrocephala* | Unidentified Fabaceae |
| Santos-Júnior 2009 | *Podocnemis erythrocephala* | Unidentified Poaceae |
| Setlalekgomo and Sesiny 2014 | *Psammobates oculifer* | *Grewia flavescens* |
| Setlalekgomo and Sesiny 2014 | *Psammobates oculifer* | Unidentified |
| Snider 1993 | *Gopherus agassizii* | *Opuntia engelmannii* |
| Stone and Moll 2006 | *Terrapene carolina* | *Fragaria* sp. |
| Stone and Moll 2006 | *Terrapene carolina* | *Podophyllum peltatum* |
| Stone and Moll 2006 | *Terrapene carolina* | *Rubus allegheniensis* |
| Stone and Moll 2006 | *Terrapene ornata* | *Fragaria* sp. |
| Stone and Moll 2006 | *Terrapene ornata* | *Podophyllum peltatum* |
| Stone and Moll 2006 | *Terrapene ornata* | *Rubus allegheniensis* |
| Stone and Moll 2009 | *Terrapene carolina* | *Ambrosia artemisiifolia* |
| Stone and Moll 2009 | *Terrapene carolina* | *Celtis* sp. |
| Stone and Moll 2009 | *Terrapene carolina* | *Cornus* sp. |
| Stone and Moll 2009 | *Terrapene carolina* | *Fragaria virginiana* |
| Stone and Moll 2009 | *Terrapene carolina* | *Morus* sp. |
| Stone and Moll 2009 | *Terrapene carolina* | *Passiflora* sp. |
| Stone and Moll 2009 | *Terrapene carolina* | *Phytolacca americana* |
| Stone and Moll 2009 | *Terrapene carolina* | *Platanus occidentalis* |
| Stone and Moll 2009 | *Terrapene carolina* | *Podophyllum peltatum* |
| Stone and Moll 2009 | *Terrapene carolina* | *Rubus* sp. |
| Stone and Moll 2009 | *Terrapene carolina* | *Vaccinium* sp. |
| Stone and Moll 2009 | *Terrapene carolina* | *Vitis* sp. |
| Stone and Moll 2009 | *Terrapene carolina* | Unidentified Polygonaceae |
| Stone and Moll 2009 | *Terrapene carolina* | Unidentified |
| Stone and Moll 2009 | *Terrapene ornata* | *Fragaria virginiana* |
| Stone and Moll 2009 | *Terrapene ornata* | *Galium* sp. |
| Stone and Moll 2009 | *Terrapene ornata* | *Morus* sp. |
| Stone and Moll 2009 | *Terrapene ornata* | *Prunus* sp. |
| Stone and Moll 2009 | *Terrapene ornata* | *Rubus* sp. |
| Stone and Moll 2009 | *Terrapene ornata* | Unidentified Polygonaceae |
| Stone and Moll 2009 | *Terrapene ornata* | Unidentified Poaceae |
| Stone and Moll 2009 | *Terrapene ornata* | Unidentified Cyperaceae |
| Strong and Fragoso 2006 | *Chelonoidis carbonaria* | *Aechmea* sp. |
| Strong and Fragoso 2006 | *Chelonoidis carbonaria* | *Ficus* sp. |
| Strong and Fragoso 2006 | *Chelonoidis carbonaria* | *Genipa americana* |
| Strong and Fragoso 2006 | *Chelonoidis denticulata* | *Aechmea* sp. |
| Strong and Fragoso 2006 | *Chelonoidis denticulata* | *Ficus* sp. |
| Strong and Fragoso 2006 | *Chelonoidis denticulata* | *Genipa americana* |
| Sung et. al. 2016 | *Platysternon megacephalum* | *Ficus* sp. |
| Sung et. al. 2016 | *Platysternon megacephalum* | *Machilus breviflora* |
| Sung et. al. 2016 | *Platysternon megacephalum* | *Machilus thunbergii* |
| Sung et. al. 2016 | *Platysternon megacephalum* | *Turpinia arguta* |
| Teran et. al. 1995 | *Phrynops geoffroanus* | *Diospyros* sp. |
| Teran et. al. 1995 | *Phrynops geoffroanus* | *Margaritaria nobilis* |
| Teran et. al. 1995 | *Phrynops geoffroanus* | *Maripa* sp. |
| Teran et. al. 1995 | *Phrynops geoffroanus* | *Pouteria* sp. |
| Teran et. al. 1995 | *Phrynops geoffroanus* | Unidentified Fabaceae |
| Tol et. al. 2017 | *Chelonia mydas* | *Halodule uninervis* |
| Tol et. al. 2017 | *Chelonia mydas* | *Halophila decipiens* |
| Tol et. al. 2017 | *Chelonia mydas* | *Zostera muelleri* |
| Tulipani and Lipcius 2014 | *Malaclemys terrapin* | *Zostera marina* |
| Turner et. al. 1984 | *Gopherus agassizii* | *Opuntia* sp. |
| van Dijk 1998 | *Indotestudo elongata* | *Cyanotis cristata* |
| van Dijk 1998 | *Indotestudo elongata* | *Dillenia* sp. |
| van Dijk 1998 | *Indotestudo elongata* | *Ficus racemosa* |
| van Dijk 1998 | *Indotestudo elongata* | *Olax scandens* |
| Varela and Bucher 2002 | *Chelonoidis chilensis* | *Celtis pallida* |
| Varela and Bucher 2002 | *Chelonoidis chilensis* | *Prosopis elata* |
| Varela and Bucher 2002 | *Chelonoidis chilensis* | *Prosopis nigra* |
| Varela and Bucher 2002 | *Chelonoidis chilensis* | *Prosopis torquata* |
| Varela and Bucher 2002 | *Chelonoidis chilensis* | *Ziziphus mistol* |
| Varela and Bucher 2002 | *Chelonoidis chilensis* | *Celtis pallida* |
| Varela and Bucher 2002 | *Chelonoidis chilensis* | *Ziziphus mistol* |
| Veerappan and Vasudevan 2012 | *Indotestudo travancorica* | *Dillenia pentagyna* |
| Vijaya 1982 | *Indotestudo forsteni* | *Artocarpus heterophyllus* |
| Vijaya 1982 | *Vijayachelys silvatica* | *Artocarpus heterophyllus* |
| Vijaya 1982 | *Vijayachelys silvatica* | *Dillenia pentagyna* |
| Vijaya 1982 | *Vijayachelys silvatica* | *Cordia peruviana* |
| Vijaya 1983 | *Indotestudo travancorica* | *Artocarpus heterophyllus* |
| Vijaya 1983 | *Indotestudo travancorica* | *Dillenia pentagyna* |
| Vogt and Guzmán 1988 | *Kinosternon leucostomum* | *Ficus* sp. |
| Vogt and Guzmán 1988 | *Kinosternon leucostomum* | *Piper* sp. |
| Vogt and Guzmán 1988 | *Kinosternon leucostomum* | *Pulcheni armata* |
| Vogt and Guzmán 1988 | *Staurotypus triporcatus* | *Diospyros nigra* |
| Vogt et. al. 2009 | *Rhinoclemmys aerolata* | *Byrsonima crassifolia* |
| Vogt et. al. 2009 | *Rhinoclemmys aerolata* | *Eugenia* sp. |
| Vogt et. al. 2009 | *Rhinoclemmys aerolata* | *Miconia* sp. |
| Waibel et. al. 2012 | *Aldabrachelys gigantea* | *Lantana camara* |
| Waibel et. al. 2012 | *Aldabrachelys gigantea* | *Mimusops coriacea* |
| Waibel et. al. 2012 | *Aldabrachelys gigantea* | *Wikstroemia indica* |
| Waibel et. al. 2012 | *Aldabrachelys gigantea* | *Adonidia merrillii* |
| Waller et. al. 1989 | *Chelonoidis chilensis* | *Goldmanceggea glauca* |
| Waller et. al. 1989 | *Chelonoidis chilensis* | *Cereus aethiops* |
| Waller et. al. 1989 | *Chelonoidis chilensis* | *Daucus pusillus* |
| Waller et. al. 1989 | *Chelonoidis chilensis* | *Geoffroea decorticans* |
| Waller et. al. 1989 | *Chelonoidis chilensis* | *Monttea aphylla* |
| Waller et. al. 1989 | *Chelonoidis chilensis* | *Plantago patagonica* |
| Waller et. al. 1989 | *Chelonoidis chilensis* | *Prosopis alpataco* |
| Waller et. al. 1989 | *Chelonoidis chilensis* | *Schismus barbatus* |
| Wang et. al. 2011 | *Chelonoidis carbonaria* | *Acrocomia aculeata* |
| Wang et. al. 2011 | *Chelonoidis carbonaria* | *Agonandra brasiliensis* |
| Wang et. al. 2011 | *Chelonoidis carbonaria* | *Annona cornifolia* |
| Wang et. al. 2011 | *Chelonoidis carbonaria* | *Annona dioica* |
| Wang et. al. 2011 | *Chelonoidis carbonaria* | *Ficus* sp. |
| Wang et. al. 2011 | *Chelonoidis carbonaria* | *Genipa americana* |
| Wang et. al. 2011 | *Chelonoidis carbonaria* | *Hancornia speciosa* |
| Wang et. al. 2011 | *Chelonoidis carbonaria* | *Mouriri elliptica* |
| Wang et. al. 2011 | *Chelonoidis carbonaria* | *Pouteria gardneri* |
| Wang et. al. 2011 | *Chelonoidis carbonaria* | *Protium heptaphyllum* |
| Wang et. al. 2011 | *Chelonoidis carbonaria* | *Psidium nutans* |
| Wang et. al. 2011 | *Chelonoidis carbonaria* | *Psidium guajava* |
| Wang et. al. 2011 | *Chelonoidis carbonaria* | *Syagrus flexuosa* |
| Wang et. al. 2011 | *Chelonoidis carbonaria* | *Syzygium cumini* |
| Wang et. al. 2011 | *Chelonoidis carbonaria* | *Vitex cymosa* |
| Wang et. al. 2011 | *Chelonoidis carbonaria* | *Cordia sessilis* |
| Wang et. al. 2011 | *Chelonoidis carbonaria* | *Byrsonima cydoniifolia* |
| Whitaker 2009 | *Vijayachelys silvatica* | *Ficus pertusa* |
| Wilson and Lawler 2008 | *Emydura macquarii krefftii* | *Ficus* sp. |

**S3:** Studies from which data on the gut retention times (GRT) of chelonians were extracted, and chelonian mean GRT and mass. *‘NA’* indicates that the mean GRT was not available (only the range; see Fig. 6).

| Reference | Chelonian species | Mean GRT (d) | Mass (kg) |
| --- | --- | --- | --- |
| Amorocho & Reina 2008 | *Chelonia mydas* | 23.7 | 160.0 |
| Andriantsaralaza et al. 2013 | *Aldabrachelys gigantea* | NA | 117.2 |
| Barboza 1995 | *Gopherus agassizii* | 10.8 | 2.8 |
| Bjorndal 1987 | *Gopherus polyphemus* | 13.0 | 4.1 |
| Bjorndal 1989 | *Chelonoidis carbonaria* | 2.6 | 2.0 |
| Bjorndal 1989 | *Chelonoidis denticulata* | 3.6 | 2.0 |
| Bjorndal 1990 | *Pseudemys nelsoni* | 2.8 | 3.8 |
| Bjorndal and Bolten 1993 | *Pseudemys nelsoni* | 3.1 | 3.8 |
| Bjorndal and Bolten 1993 | *Trachemys scripta* | 4.9 | 1.9 |
| Blake et al. 2012 | *Chelonoidis nigra* | 12.0 | 175.0 |
| Braun & Brooks 1987 | *Terrapene carolina* | NA | 0.4 |
| Davenport et al. 1992 | *Batagur baska* | NA | 17.9 |
| Elbers 2010 | *Macrochelys temminckii* | NA | 78.9 |
| Falcón et al. unpubl. | *Aldabrachelys gigantea* | 15.0 | 117.2 |
| Franz et al. 2011 | *Aldabrachelys gigantea* | 6.8 | 117.2 |
| Franz et al. 2011 | *Centrochelys sulcata* | 15.9 | 43.0 |
| Franz et al. 2011 | *Chelonoidis nigra* | 8.6 | 175.0 |
| Franz et al. 2011 | *Testudo graeca* | 6.8 | 1.4 |
| Franz et al. 2011 | *Testudo hermanni* | 5.0 | 1.3 |
| Guzmán & Stevenson 2008 | *Chelonoidis denticulata* | 21.0 | 2.0 |
| Hailey 1997 | *Kinixys spekii* | 5.6 | 0.6 |
| Hailey 1997 | *Stigmochelys pardalis* | 5.2 | 20.0 |
| Hailey 1998 | *Kinixys spekii* | 5.5 | 0.6 |
| Hamilton and Coe 1982 | *Aldabrachelys gigantea* | 12.2 | 117.2 |
| Hansen et al. 2008 | *Aldabrachelys gigantea* | 14.0 | 117.2 |
| Hatt et al. 2002 | *Chelonoidis nigra* | 10.2 | 175.0 |
| Jansen & Moll 1995 | *Rhinoclemmys annulata* | 1.5 | 1.4 |
| Jansen & Moll 1995 | *Rhinoclemmys funerea* | 1.8 | 0.9 |
| Jerozolinski et al. 2009 | *Chelonoidis denticulata* | 8.3 | 2.0 |
| Kimmons & Moll 2010 | *Chelydra serpentina* | 2.0 | 5.2 |
| Kimmons & Moll 2010 | *Chelydra serpentina* | 2.0 | 5.2 |
| Kimmons & Moll 2010 | *Chelydra serpentina* | 2.3 | 5.2 |
| Kimmons & Moll 2010 | *Trachemys scripta* | 2.8 | 1.9 |
| Kimmons & Moll 2010 | *Trachemys scripta* | 2.9 | 1.9 |
| Kimmons & Moll 2010 | *Trachemys scripta* | 3.7 | 1.9 |
| Lautenschlager-Rodrigues 2016 | *Chelonoidis carbonaria* | 6.9 | 2.0 |
| Legler & Vogt 2013 | *Rhinoclemmys aereolata* | 3.0 | 0.7 |
| Lickel 2010 | *Stigmochelys pardalis* | 15.8 | 20.0 |
| Meienberger et al. 1993 | *Gopherus agassizii* | 21.5 | 2.8 |
| Parmenter 1981 | *Chelydra serpentina* | 1.3 | 5.2 |
| Parmenter 1981 | *Chrysemys picta* | 2.5 | 0.4 |
| Parmenter 1981 | *Chrysemys scripta* | 2.5 | 0.4 |
| Parmenter 1981 | *Sternotherus minor* | 2.4 | 0.2 |
| Parmenter 1981 | *Sternotherus odoratus* | 2.0 | 0.1 |
| Rick & Bowman 1961 | *Chelonoidis porteri* | NA | 175.0 |
| Sadeghayobi et al. 2011 | *Chelonoidis nigra* | 10.1 | 175.0 |
| Setlalekgomo & Sesiny 2014 | *Psammobates oculifer* | NA | 0.3 |
| Setlalekgomo and Sesinyi 2014 | *Psammobates oculifer* | 4.0 | 0.3 |
| Stone & Moll 2006 | *Terrapene carolina* | NA | 0.4 |
| Stone & Moll 2006 | *Terrapene ornata* | NA | 0.4 |
| Tracy et al. 2006 | *Gopherus agassizii* | 9.5 | 2.8 |
| Valente et al. 2008 | *Caretta caretta* | 11.5 | 109.2 |
| Varela & Bucher 2002 | *Chelonoidis chilensis* | 8.0 | 3.2 |
| Waibel et al. 2012 | *Aldabrachelys gigantea* | 15.5 | 117.2 |

**S4:** Studies from which data on the home range size of chelonians were extracted, and chelonian mean home range and mass. See Figure 7a for ranges (minimum and maximum home range size).

| Reference | Chelonian species | Home range (ha) | Mass (kg) |
| --- | --- | --- | --- |
| Barret 1990 | *Gopherus agassizii* | 19.0 | 2.8 |
| Baxter 2015 | *Aldabrachelys gigantea* | 10.5 | 117.2 |
| Bernstein et al. 2007 | *Terrapene ornata* | 5.8 | 0.4 |
| Bridget and Echternacht 2009 | *Terrapene carolina* | 2.3 | 0.4 |
| Carter et al. 1999 | *Clemmys muhlenbergii* | 0.5 | 0.2 |
| Chase et al. 1989 | *Clemmys muhlenbergii* | 0.1 | 0.2 |
| Diemer 1992 | *Gopherus polyphemus* | 0.9 | 4.1 |
| Doroff and Keith 1990 | *Terrapene ornata* | 8.7 | 0.4 |
| Duda et al. 1999 | *Gopherus agassizii* | 12.5 | 2.8 |
| Edge et al. 2015 | *Emydoidea blandingii* | 59.2 | 1.2 |
| Eubanks et al. 2003 | *Gopherus polyphemus* | 0.8 | 4.1 |
| Forero-Medina et al. 2012 | *Mesoclemmys dahli* | 15.3 | 0.8 |
| Franks et al. 2011 | *Gopherus agassizii* | 7.3 | 2.8 |
| Galois et al. 2002 | *Apalone spinifera* | 24.2 | 4.8 |
| Geffen and Mendelssohn 1988 | *Testudo kleinmanni* | 26.4 | 0.4 |
| Hailey and Coulson 1996 | *Kinixys spekii* | 1.9 | 0.6 |
| Hailey and Coulson 1996 | *Stigmochelys pardalis* | 26.0 | 20.0 |
| Innes et al. 2008 | *Emydoidea blandingii* | 4.9 | 1.2 |
| Jones 1996 | *Graptemys flavimaculata* | 3.5 | 0.9 |
| Judd and Rose 1983 | *Gopherus berlandieri* | 0.4 | 1.8 |
| Lawson 2006 | *Kinixys erosa* | 14.2 | 1.1 |
| Lawson 2006 | *Kinixys homeana* | 20.0 | 1.1 |
| Litzgus and Mousseau 2004 | *Clemmys guttata* | 7.5 | 0.2 |
| Lue and Chen 1999 | *Cuora flavomarginata* | 1.1 | 0.5 |
| Mazzotti et al. 2002 | *Testudo hermanni* | 6.0 | 1.3 |
| McMaster and Downs 2009 | *Stigmochelys pardalis* | 122.4 | 20.0 |
| Millar and Blouin-Demers 2011 | *Emydoidea blandingii* | 12.0 | 1.2 |
| Morrow et al. 2001 | *Clemmys muhlenbergii* | 0.6 | 0.2 |
| Moskovits and Kiester 1987 | *Chelonoidis carbonaria* | 26.3 | 2.0 |
| Moskovits and Kiester 1987 | *Chelonoidis denticulata* | 36.1 | 2.0 |
| Nieuwot 1996 | *Terrapene ornata* | 1.6 | 0.4 |
| O'connor et al. 1994 | *Gopherus agassizii* | 27.3 | 2.8 |
| Obbard and Brooks 1981 | *Chelydra serpentina* | 3.5 | 5.2 |
| Roe and Arthur 2008 | *Chelodina longicollis* | 11.5 | 1.3 |
| Ross and Anderson 1990 | *Emydoidea blandingii* | 0.6 | 1.2 |
| Rowe 2003 | *Chrysemys picta* | 1.2 | 0.4 |
| Rowe and Moll 1991 | *Emydoidea blandingii* | 84.5 | 1.2 |
| Seminoff et al. 2002 | *Chelonia mydas* | 1662.0 | 160.0 |
| Smith and Cherry 2016 | *Glyptemys muhlenbergii* | 0.8 | 0.1 |
| Stickel 1989 | *Terrapene carolina* | 1.2 | 0.4 |
| Strang 1983 | *Clemmys insculpta* | 0.0 | 0.2 |
| Strang 1983 | *Terrapene carolina* | 0.0 | 0.4 |

**S5:** Studies from which data on the displacement distances of chelonians were extracted, and chelonian mean displacement distance and mass. *‘NA’* indicates that the mean displacement distance was not available (only the range; see Fig. 7b).

| Reference | Chelonian species | Displacement (m d<sup>-1</sup>) | Mass (kg) |
| --- | --- | --- | --- |
| Baxter 2015 | *Aldabrachelys gigantea* | 191.8 | 117.2 |
| Birkhead et al. 2005 | *Gopherus polyphemus* | NA | 4.1 |
| Brown and Brooks 1993 | *Chelydra serpentina* | 300.5 | 5.2 |
| Díaz-Paniagua | *Testudo graeca* | 50.0 | 1.4 |
| Duda et al. 1999 | *Gopherus agassizii* | 41.7 | 2.8 |
| Geffen and Mendelssohn 1988 | *Testudo kleinmanni* | 26.0 | 0.4 |
| Guzmán and Stevenson 2008 | *Chelonoidis denticulata* | NA | 2.0 |
| Hailey 1989 | *Testudo hermanni* | 78.0 | 1.3 |
| Hailey and Coulson 1996 | *Kinixys spekii* | 172.0 | 0.6 |
| Hailey and Coulson 1996 | *Stigmochelys pardalis* | 435.0 | 20.0 |
| Innes et al. 2008 | *Emydoidea blandingii* | 30.9 | 1.2 |
| Kimmons & Moll 2010 | *Trachemys scripta* | NA | 1.9 |
| Kimmons & Moll 2010 | *Chelydra serpentina* | NA | 5.2 |
| Lambiris et al. 1989 | *Kinixys spekii* | NA | 0.6 |
| Mazzotti et al. 2002 | *Testudo hermanni* | 55.9 | 1.3 |
| Millar and Blouin-Demers 2011 | *Emydoidea blandingii* | 214.8 | 1.2 |
| Moll & Jansen 1995 | *Rhinoclemmys funerea* | NA | 0.9 |
| Moll & Jansen 1995 | *Rhinoclemmys annulata* | NA | 1.4 |
| Morrow et al. 2001 | *Clemmys muhlenbergii* | 3.3 | 0.2 |
| Moskovits and Kiester 1987 | *Chelonoidis carbonaria* | 0.4 | 2.0 |
| Moskovits and Kiester 1987 | *Chelonoidis denticulata* | 0.5 | 2.0 |
| Nieuwot 1996 | *Terrapene ornata* | 13.4 | 0.4 |
| Ross and Anderson 1990 | *Emydoidea blandingii* | 71.4 | 1.2 |
| Rowe 2003 | *Chrysemys picta* | 250.8 | 0.4 |
| Rowe and Moll 1991 | *Emydoidea blandingii* | 40.4 | 1.2 |
| Smith and Cherry 2016 | *Glyptemys muhlenbergii* | 14.1 | 0.1 |
| Strang 1983 | *Clemmys insculpta* | 108.0 | 0.2 |
| Strang 1983 | *Terrapene carolina* | 40.0 | 0.4 |
| Travis et al. 2014 | *Chelydra serpentina* | 147.5 | 5.2 |

**S6:**
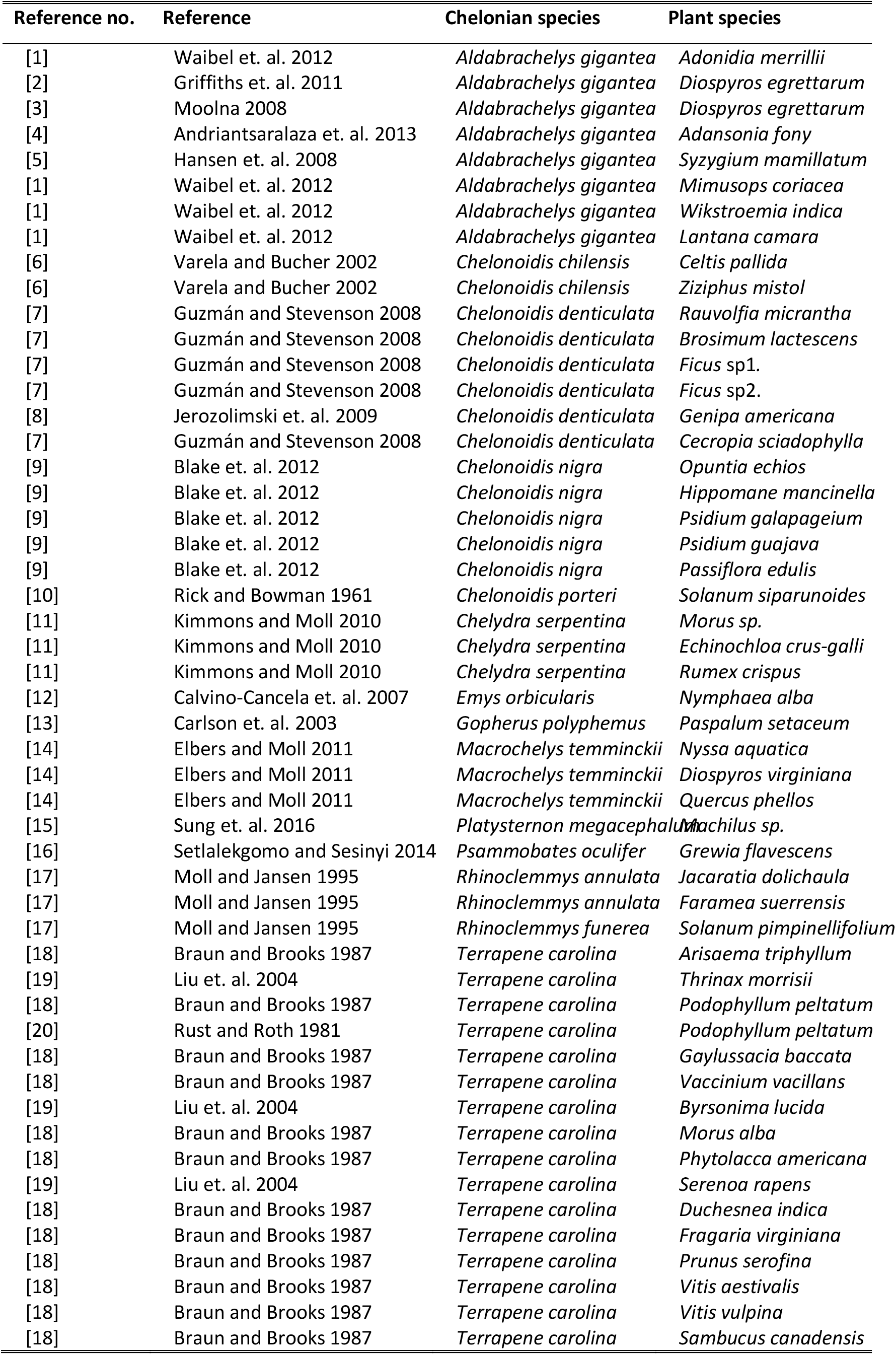
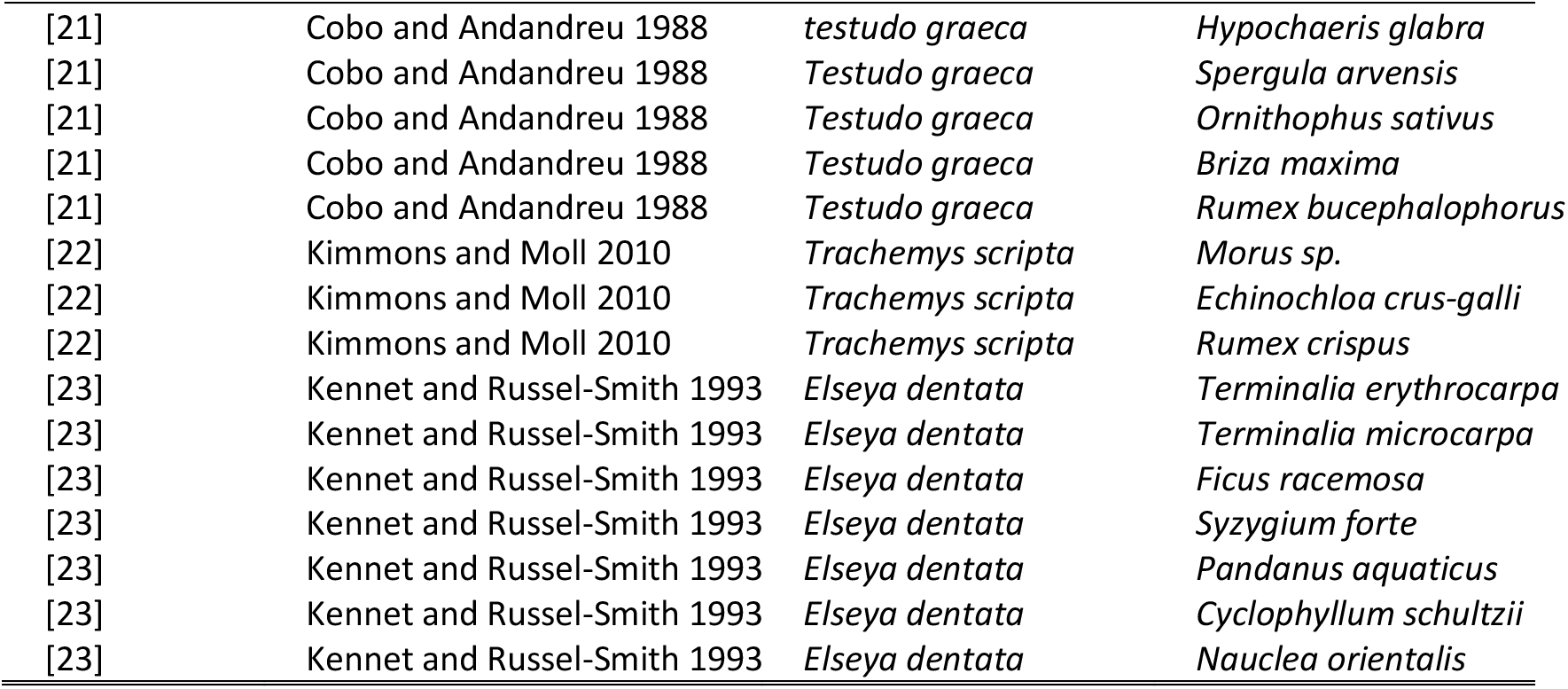
Studies from which data on the effect of chelonian gut passage on germination were extracted. References for Table 2 in the main text.

| Reference no. | Reference | Chelonian species | Plant species |
| --- | --- | --- | --- |
| [1] | Waibel et. al. 2012 | *Aldabrachelys gigantea* | *Adonidia merrillii* |
| [2] | Griffiths et. al. 2011 | *Aldabrachelys gigantea* | *Diospyros egrettarum* |
| [3] | Moolna 2008 | *Aldabrachelys gigantea* | *Diospyros egrettarum* |
| [4] | Andriantsaralaza et. al. 2013 | *Aldabrachelys gigantea* | *Adansonia fony* |
| [5] | Hansen et. al. 2008 | *Aldabrachelys gigantea* | *Syzygium mamillatum* |
| [1] | Waibel et. al. 2012 | *Aldabrachelys gigantea* | *Mimusops coriacea* |
| [1] | Waibel et. al. 2012 | *Aldabrachelys gigantea* | *Wikstroemia indica* |
| [1] | Waibel et. al. 2012 | *Aldabrachelys gigantea* | *Lantana camara* |
| [6] | Varela and Bucher 2002 | *Chelonoidis chilensis* | *Celtis pallida* |
| [6] | Varela and Bucher 2002 | *Chelonoidis chilensis* | *Ziziphus mistol* |
| [7] | Guzmán and Stevenson 2008 | *Chelonoidis denticulata* | *Rauvolfia micrantha* |
| [7] | Guzmán and Stevenson 2008 | *Chelonoidis denticulata* | *Brosimum lactescens* |
| [7] | Guzmán and Stevenson 2008 | *Chelonoidis denticulata* | *Ficus* sp1. |
| [7] | Guzmán and Stevenson 2008 | *Chelonoidis denticulata* | *Ficus* sp2. |
| [8] | Jerozolinski et. al. 2009 | *Chelonoidis denticulata* | *Genipa americana* |
| [7] | Guzmán and Stevenson 2008 | *Chelonoidis denticulata* | *Cecropia sciadophylla* |
| [9] | Blake et. al. 2012 | *Chelonoidis nigra* | *Opuntia echios* |
| [9] | Blake et. al. 2012 | *Chelonoidis nigra* | *Hippomane mancinella* |
| [9] | Blake et. al. 2012 | *Chelonoidis nigra* | *Psidium galapageium* |
| [9] | Blake et. al. 2012 | *Chelonoidis nigra* | *Psidium guajava* |
| [9] | Blake et. al. 2012 | *Chelonoidis nigra* | *Passiflora edulis* |
| [10] | Rick and Bowman 1961 | *Chelonoidis porteri* | *Solanum siparunoides* |
| [11] | Kimmons and Moll 2010 | *Chelydra serpentina* | *Morus* sp. |
| [11] | Kimmons and Moll 2010 | *Chelydra serpentina* | *Echinochloa crus-galli* |
| [11] | Kimmons and Moll 2010 | *Chelydra serpentina* | *Rumex crispus* |
| [12] | Calvino-Cancela et. al. 2007 | *Emys orbicularis* | *Nymphaea alba* |
| [13] | Carlson et. al. 2003 | *Gopherus polyphemus* | *Paspalum setaceum* |
| [14] | Elbers and Moll 2011 | *Macrochelys temminckii* | *Nyssa aquatica* |
| [14] | Elbers and Moll 2011 | *Macrochelys temminckii* | *Diospyros virginiana* |
| [14] | Elbers and Moll 2011 | *Macrochelys temminckii* | *Quercus phellos* |
| [15] | Sung et. al. 2016 | *Platysternon megacephalum* | *Machilus* sp. |
| [16] | Setlalekgomo and Sesinyi 2014 | *Psammobates oculifer* | *Grewia flavescens* |
| [17] | Moll and Jansen 1995 | *Rhinoclemmys annulata* | *Jacaratia dolichaula* |
| [17] | Moll and Jansen 1995 | *Rhinoclemmys annulata* | *Faramaea suerrensensis* |
| [17] | Moll and Jansen 1995 | *Rhinoclemmys funerea* | *Solanum pimpinellifolium* |
| [18] | Braun and Brooks 1987 | *Terrapene carolina* | *Arisaema triphyllum* |
| [19] | Liu et. al. 2004 | *Terrapene carolina* | *Thrinax morrisii* |
| [18] | Braun and Brooks 1987 | *Terrapene carolina* | *Podophyllum peltatum* |
| [20] | Rust and Roth 1981 | *Terrapene carolina* | *Podophyllum peltatum* |
| [18] | Braun and Brooks 1987 | *Terrapene carolina* | *Gaylussacia baccata* |
| [18] | Braun and Brooks 1987 | *Terrapene carolina* | *Vaccinium vacillans* |
| [19] | Liu et. al. 2004 | *Terrapene carolina* | *Byrsonima lucida* |
| [18] | Braun and Brooks 1987 | *Terrapene carolina* | *Morus alba* |
| [18] | Braun and Brooks 1987 | *Terrapene carolina* | *Phytolacca americana* |
| [19] | Liu et. al. 2004 | *Terrapene carolina* | *Serenoa rapens* |
| [18] | Braun and Brooks 1987 | *Terrapene carolina* | *Duchesnea indica* |
| [18] | Braun and Brooks 1987 | *Terrapene carolina* | *Fragaria virginiana* |
| [18] | Braun and Brooks 1987 | *Terrapene carolina* | *Prunus serotina* |
| [18] | Braun and Brooks 1987 | *Terrapene carolina* | *Vitis aestivalis* |
| [18] | Braun and Brooks 1987 | *Terrapene carolina* | *Vitis vulpina* |
| [18] | Braun and Brooks 1987 | *Terrapene carolina* | *Sambucus canadensis* |
| [21] | Cobo and Andandreu 1988 | *testudo graeca* | *Hypochaeris glabra* |
| [21] | Cobo and Andandreu 1988 | *Testudo graeca* | *Spergula arvensis* |
| [21] | Cobo and Andandreu 1988 | *Testudo graeca* | *Ornithophus sativus* |
| [21] | Cobo and Andandreu 1988 | *Testudo graeca* | *Briza maxima* |
| [21] | Cobo and Andandreu 1988 | *Testudo graeca* | *Rumex bucephalophorus* |
| [22] | Kimmons and Moll 2010 | *Trachemys scripta* | *Morus sp.* |
| [22] | Kimmons and Moll 2010 | *Trachemys scripta* | *Echinochloa crus-galli* |
| [22] | Kimmons and Moll 2010 | *Trachemys scripta* | *Rumex crispus* |
| [23] | Kennet and Russel-Smith 1993 | *Elseya dentata* | *Terminalia erythrocarpa* |
| [23] | Kennet and Russel-Smith 1993 | *Elseya dentata* | *Terminalia microcarpa* |
| [23] | Kennet and Russel-Smith 1993 | *Elseya dentata* | *Ficus racemosa* |
| [23] | Kennet and Russel-Smith 1993 | *Elseya dentata* | *Syzygium forte* |
| [23] | Kennet and Russel-Smith 1993 | *Elseya dentata* | *Pandanus aquaticus* |
| [23] | Kennet and Russel-Smith 1993 | *Elseya dentata* | *Cyclophyllum schultzii* |
| [23] | Kennet and Russel-Smith 1993 | *Elseya dentata* | *Nauclea orientalis* |

